# Characterization of starch metabolism pathways in dinoflagellates, chromerids and apicomplexans highlights the presence of a starch accumulating symbiont in their common ancestor

**DOI:** 10.64898/2026.09.25.754361

**Authors:** Cenci Ugo, Malika Chabi, Léa Fermont, Philippe Deschamps, Marie Leleu, Catherine Tirtiaux, John M. Archibald, Miroslav Oborník, Colleoni Christophe, Steven Ball

**Author notes:** Corresponding authors (S.B.). These authors contributed equally to this work (Cenci Ugo, Malika Chabi, Léa Fermont).

## Abstract

Unlike most other heterotrophic protist lineages which synthesize glycogen, alveolates accumulate α-glucan storage polysaccharides in the form of starch. Within Alveolata, ciliates do so by using a simple pathway analogous to that used for glycogen metabolism in eukaryotes. However, Myzozoa have evolved five additional enzymatic steps which are common to Dinoflagellata, Chromerids and Apicomplexa. Moreover, dinoflagellates have acquired seven additional enzymes that distinguish them from the Chromerids-Apicompexa lineages, while chromerids possess only a single enzyme not found in dinoflagellates. Such modifications are not found in other eukaryotic lineages save for those resulting directly from primary or secondary plastid endosymbiosis. The many differences affecting the starch pathway of Myzozoa can be explained by a single secondary or tertiary plastid endosymbiosis of a starch-containing alga followed by distinct compartmentalisation outcomes of starch metabolism in Dinoflagellata relative to chromerids and Apicomplexa.

## INTRODUCTION

Alveolata is a large monophyletic group of unicellular eukaryotes harbouring cortical alveoli below the membrane surface (*1*). The two major subgroups of alveolates are the ciliates (Ciliophora) and Myzozoa (*2*). Ciliates are primarily heterotrophic and mostly phagotrophic protists with little or no evidence supporting the hypothesis that they evolved from a plastid-bearing ancestor (*3*). Myzozoa are alveolates that feed by myzocytosis or whose ancestors displayed this capacity (*2*). Dinoflagellata on the one hand and the Chromerids/Apicomplexa parasite lineages on the other define the two major groups of Myzozoa. Within Myozoza, presence of plastid serves as evidence of a past endosymbiotic origin, which may represent a key factor in their diversification. (*4*, *5*). Photosynthetic and non-photosynthetic plastids are indeed common in these organisms and two major synapomorphies related to photosynthesis unite all photosynthetic Myzozoa (*6*, *7*): the acquisition by lateral gene transfer (LGT) from proteobacteria to the alveolate nuclear genome of a Rubisco II subunit gene (*6*) and the presence of RNA polyuridination in plastids (*7*). It was recently proposed that an ancestral myzozoan plastid likely originated from a quaternary endosymbiotic event with a stramenopile (*8–13*). However, myzozoans have experienced independent gains of temporary and permanent plastids, multiple plastid replacements, losses of photosynthesis, and, more rarely, complete plastid loss, all of which together further obscure efforts to reconstruct their ancient endosymbiotic history (*4*).

Alveolates, unlike most eukaryotes, save the Archaeplastida and Cryptophyceae, most often accumulate starch as storage material (*14*). α-1,4-linked glucose chains branched through α-1,6 linkages define the most widespread form of storage polysaccharides in living cells (*14*). These glucans come in two very distinct forms known as glycogen and starch. The small (< 50nm) water-soluble glycogen particles distribute widely in Archaea, Bacteria, and heterotrophic eukaryotes. However, synthesis of semicrystalline solid starch granules of unlimited size (from 50 nm to several µm) seems to be restricted to photosynthetic eukaryotes resulting from Archaeplastida, Cryptista and alveolates including their photosynthetic derivates that include Cryptophyceae and Myzozoa. Starch accumulating prokaryotes are limited to a few rare single-celled diazotrophic Cyanobacteria, and starch-like granulose particles accumulating Clostridia as well as some Verrucomicrobia(*14–17*). In eukaryotes synthesizing starch, the presence of semi-crystalline polysaccharide dramatically affects the accessibility of the polysaccharide to the enzymes of starch breakdown. In all eukaryotes, polysaccharide mobilisation of starch requires the prior phosphorylation of glucose residues present in the crystallites through GWD (glucan-water dikinases) thereby opening the crystallites and rendering the hydrophobic structures accessible to degradation (*18–20*).

All descendants from primary plastid endosymbiosis (the Archaeplastida) accumulate starch either in the cytosol (red algae and glaucophytes) or in their plastids (green algae and plants) (*21*). The pathway of starch biosynthesis in Archaeplastida includes all enzymes of eukaryotic glycogen metabolism present in most Amoebozoa and metamonads (*16*, *22*). This includes β-amylase and amylomaltases which have been lost from the more intensively studied glycogen accumulating fungi and animals as well as in a few parasitic metamonads.

The differences between the syntheses of glycogen and starch have been demonstrated to be dependent on the presence of a prokaryotic isoamylase in all green algae and plants tested (*20*, *23*). In the absence of this enzyme, mutants switch to glycogen synthesis with a variable but consequent decrease (10 to 95%) of storage polysaccharide content(*24–28*). Mutants of the red alga *Cyanidioschizon merolae* defective in one of two of its cytosolic isoamylases behaves similarly (*29*). In addition, mutants of the starch accumulating *Cyanobacterium* sp. Clg1 were equally demonstrated to switch back to the decreased synthesis of glycogen if one of its two isoamylase-like genes was inactivated or altered by mutations (*30*). Because the *Cyanobacterium* sp. Clg1 isoamylase is only very distantly related to the archaeplastidal enzymes, and because the cyanobacterial glycogen metabolism enzyme network is very different from that of eukaryotes, this implies that isoamylase recruitment is mandatory whenever a eukaryotic or prokaryotic lineage is evolving from glycogen to starch metabolism. It is thought that isoamylase displays a dual function (*31*). The first function would consist of catalytic removal by enzymatic debranching of those chains that prevent crystallisation of neighbouring glucans. The second function is thought to be non-catalytic and due to the interaction of the complex quaternary structure of the multi-subunit isoamylase with the surface of the growing polysaccharide, further speeding up its aggregation into semi-crystalline granules (*31*). However, the mandatory nature of isoamylase to generate starch structures in biological systems may be questioned by several observations. Among these, quadruple mutants of *Arabidopsis* affected in debranching enzymes with no traces of isoamylase and pullulanase can be partly suppressed by inactivation of AMY3, one of the 3 plastidial α-amylases thereby allowing for the synthesis of starch-like granular structures (*28*).

In addition, a yeast semi-*in vitro* system expressing enzymes of either the maize endosperm or Arabidopsis leaves in the absence of functional yeast glycogen metabolism resulted in significant synthesis of starch-like structures even in the absence of isoamylase (*32*). Nevertheless, the addition of isoamylase to these semi-*in vitro* systems significantly stimulated the heterologous production of starch in yeast.

To clarify the mandatory nature of polysaccharide debranching to obtain starch in living cells, we are studying starch metabolism in starch accumulating organisms apparently lacking isoamylase. Alveolates, with the noticeable exception of dinoflagellates, are such organisms. We have concentrated our efforts on the chromerid *Chromera velia* and have reported that the major determinants of polysaccharide aggregation into starch-like granules surprisingly reside in the concerted action of phosphorylase and indirect debranching enzymes (Chabi et al. submitted). During this study we also showed that two previously unreported enzymes were required to obtain significant starch synthesis in *Chromera velia*. To understand the emergence of such novel features of storage polysaccharide metabolism we undertook a study of its evolution in alveolates. We show that alveolates have undergone considerable evolution of their storage polysaccharide metabolism. Starting from a relatively simple platform of starch synthesis derived from eukaryotic cytosolic glycogen metabolism that we now report for ciliates, we demonstrate the presence of five additional enzymes common to all Myzozoa that were introduced by LGT/EGT. In addition, we show the presence in dinoflagellates of another set of seven enzymes not found in the chromerid apicomplexa lineage including the presence of an isoamylase of red alga origin. These results can be interpreted as evidence for integration of a starch synthesizing symbiont common to all Myzozoa of either a red alga or cryptophyte origin.

Furthermore, we propose a scenario which accounts for the selective presence of isoamylase in the dinoflagellate lineage within Alveolata.

## RESULTS

### The ciliates possess a simple glycogen-like metabolism enzyme network

Depending on the species (Fig. 1A), alveolates have been reported to contain glycogen, starch granules, or “amylopectin granules” (also referred to as paraglycogen or zooamylon) (*33–35*). Starch granules can be distinguished from amylopectin granules by the additional presence of amylose, a primarily linear α-1,4 linked polyglucan synthesized within the amylopectin matrix by the elongation enzyme granule bound starch synthase (GBSS) (*36*, *37*). To identify candidate enzymes potentially responsible for the semi-crystalline nature of the insoluble polymers (amylopectin or starch), the genomes of species in the alveolate clade were surveyed for presence of known enzymes previously demonstrated to be involved in the production or catabolism of both glycogen and starch and compared to the corresponding enzyme content of various eukaryotes (Fig. 1B). The role of the major enzymes of starch/glycogen metabolism is summarized in Table 1. The glycogen-accumulating clades in Fig. 1B belong to the Opisthokonta (yeast) the Amoebozoa (*D. discoideum*), the metamonads (*T. vaginalis*, *G. intestinalis*), the Chloroplastida (*A. thaliana*, *C. reinhardtii*, *O. taurii*), the Rhodophycea (*C. crispus*, *P. purpureum*, *C. merolae*), Glaucophyta (*C. paradoxa*), Cryptista (*G. theta*, *G. avonlea*), the Myzozoa (*V. brassicaformis*, *C. velia*, *C. parvum*, *T. gondii*, *P. glacialis*, *S. microadriaticum*, *O. marina*) and the ciliates (*O. trifallax*, *P. tetraurelia*, *T. thermophila*). The eukaryotic glycogen metabolism networks present in the cytosol where the polysaccharide is accumulated contain ad minima one form of glycogen synthase, one form of branching enzyme, one form of indirect debranching enzyme (iDBE) and one form of glycogen phosphorylase. In addition to this, eukaryotes have acquired enzymes of glycogen/starch digestion present in lysosomes or in various acidic vacuoles involved in phagotrophic prey digestion. These “phagotrophic” vacuolar enzymes include α- and β-amylases as well as various glucosidases but not the minimal pathway enzyme suite described above which is distinctively cytosolic. These phagotrophic enzymes are thus often found in protist phagotrophs that feed on glycogen/starch contained in their preys or in extracellular debris, but which do not, themselves, store glycogen nor starch in the cytosol. However, some of these glucan hydrolases may still have been recruited in the cytosol for breakdown of starch/glycogen accumulating protists not reported to contain plastids. Depending on the glycogen accumulating clades, we find either a glycogen synthase of the GT3 (Glycosyl Transferase) CAZy (Carbohydrate Active Enzyme) family (all fungi and animals and a minority of protists) or of the GT5 family (many protists) or sometimes both (a few protists). Another major difference between different networks resides in the presence or absence of β-amylase often accompanied by the DPE2 amylomaltase of CAZy family GH77 (Glycosyl hydrolase). The latter two enzymes are absent from most fungi and all animals and from some parasitic metamonads (*G. intestinalis*).

**Figure 1:**
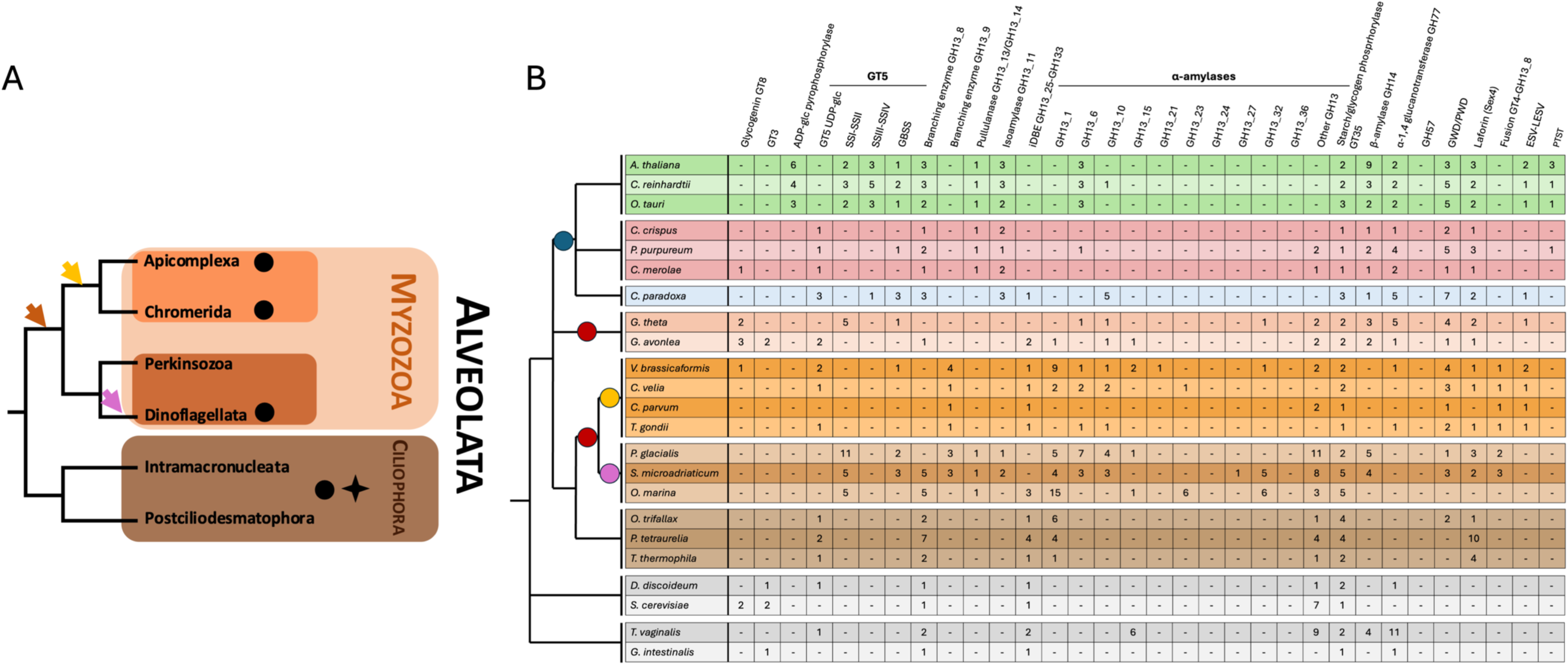
**(A) Schematic Tree Alveolata.** Polysaccharide diversity in Alveolates. Alveolates diversity is represented with Myzozoa in orange, and ciliophora in brown as the latter can be considered an outgroup when studying plastid endosymbiosis in Alveolates. The Dark orange arrow represents an hypothetical plastid acquisition at the base of the Myzozoa. The yellow and purple arrows represent putative separate endosymbioses in this clade as proposed by Ševčíková et al. 2015. Black dots represent the accumulation of crystalline storage polysaccharides while the star represents the accumulation of water-soluble storage polysaccharides (glycogen) that occurred in late diverging ciliates (Paramecium, Tetrahymena). **(B)** Comparative composition of storage polysaccharide metabolism enzyme networks across major eukaryotic lineages. The number of enzyme isoforms is listed for each eukaryotic clade, organized according to the phylogeny of eukaryotes. Chloroplastida in green; Rhodophycea in red; Glaucophyta are represented in blue; Cryptophyceae in light orange; Myzozoa in orange to dark orange (Apicomplexa and Chromerida in orange, Dinophyceae in dark orange); Ciliophora in brown; and major heterotrophic glycogen accumulating eukaryotic clades (Amoebozoa, Fungi, or Metamonada) in grey. The blue circle represents the primary plastid endosymbiosis and red circles secondary endosymbiosis (endosymbiosis of a red alga) or higher order plastid endosymbiosis (endosymbiosis of a cryptophyte). Finally yellow and pink circles represent other higher order endosymbioses that followed selectively in the dinoflagellate and chromerid/apicomplexa lineages as suggested in (Ševčíková et al. 2015). The isoforms for each enzyme class involved in storage polysaccharide metabolism were identified using BLAST and hmm searches using dbCAN databases on genome gathered from various sources (see methods) and classified according to the CAZy database. We should stress that sequence data of *P. glacialis*, *S. microadriaticum* and *O. marina* while being the best we could find, remain likely incomplete, because of the huge dinoflagellate genome sizes. In addition, we indicate two GT35 phosphorylase isoforms in *C. velia* and *V. brassicaformis*, however we were never able to visualize on zymogram nor measure the activity of the second chromerid-specific GT35 (see Chabi et al. submitted).

**Table 1:** Proteins involved in starch metabolism with name functions, classification and description.

| ABBREVIATION | COMMON NAME | EC NUMBER | CAZY<br>CLASSIFICATION | DESCRIPTION |
| --- | --- | --- | --- | --- |
| **GLG** | Glycogenin | 2.4.1.186 | GT8 | This glycosyl transferase can self-glucosylate to initiate synthesis of storage polysaccharide |
| **GT5-SS** | Starch synthase | 2.4.1.- | GT5 | Produces chains of glucose linked in α-1,4 (α-1,4 linked glucan) |
| **SSIII/SSIV** | Starch synthase | 2.4.1.- | GT5 | Produces chains of glucan linked in α-1,4 |
| **GBSS** | Granule bound Starch synthase | 2.4.1.- | GT5 | Produces chains of α-1,4 linked glucan, only within crystalline starch granules |
| **BE** | Branching enzyme | 2.4.1.18 | GH13_8 or GH13_9 | Produces α-1,6 branched glucans |
| **IDBE** | Indirect Debranching enzyme | 2.4.1.25 and 3.2.1.33 | GH13_25-GH133 | Transfers 3 residual α-1,4 linked glucose from a 4 glucose residue outer polysaccharide chain to a neighbouring outer chain before hydrolysing the remaining glucose brached in α-1,6 position |
| **ISA** | Isoamylase | 3.2.1.68 | GH13_11 | Hydrolyses the α-1,6 linkage from polysaccharide outer chains |
| **ESV-LESV** | (Like-)Early starvation | - | - | Protein associated with starch granules and likely used in scaffolding and other non-enzymatic functions |
| **DPE1, DPE2** | Disproportionating enzyme 1 and 2 | 2.4.1.25 | GH77 | Uses maltooligosaccharides to elongate chains of glucose linked in α-1,4 through oligosaccharide transfers |
| **PHO** | Glycogen/starch phosphorylase | 2.4.1.1 | GT35 | Releases glucose-1-phosphate from the non-reducing end of branched polysaccharides or oligosaccharides |
| **B-AMY** | β-amylase | 3.2.1.2 | GH14 | Hydrolyses outer α-1,4 linked chains releasing maltose |
| **A-AMY** | α-amylase | 3.2.1.1 | GH13 | Hydrolyses outer and inner α-1,4 linked glucose chains releasing smaller oligosaccharides, maltose and glucose |
| **LAFORIN** | laforin | 3.1.3.16 | CBM20+phosphatase | Phosphatase preventing hyperphosphorylation of α-glucan storage polysaccharide |
| **GWD/PWD** | (Phospho)Glucan water dikinase | 2.7.9.4 | CBM45+PPDK or CBM20+PPDK | Mediates the incorporation of phosphate through a dikinase mechanism in semi-crystalline polysaccharides |
| **MEX** | Maltose Excess | - | - | Probable maltose transporter |

The ciliates contain a simple storage polysaccharide network similar to glycogen metabolism found in fungi, animals and *G. intestinalis*. However, the glycogen synthase is of the GT5 type, as in many other glycogen-accumulating protist lineages. Not surprisingly for such phagotrophs, a large number of candidate α-amylase genes are observed in ciliates (predominantly of CAZy subfamily GH13_1).

In our previous work on *C. velia* (Chabi et al. submitted) we have shown that the phylogeny of the BE2 branching enzyme is robustly monophyletic in all alveolates including ciliates with strong support (Fig. 2A). Here we analyze the phylogenies of the GT5 glycogen/starch synthase (Figs. 2D and S1), of iDBE (Figs. 2B and S2) and of glycogen/starch phosphorylase (Figs. 2C and S3) The bootstrap for alveolate monophyly is lower, albeit significant for both glycogen/starch phosphorylase (BS=60) and for iDBE (BS=54). The phylogeny of the GT5 glycogen synthase on the other hand displays a complex topology which nevertheless remains compatible with a monophyletic origin of this enzyme for the ciliates and chromerid/apicomplexan clade, albeit with very low or no support (BS=13) (Fig. S1). Several starch phosphorylase isoforms (2 to 4) of common origin have diversified among the monophyletic alveolate enzymes of ciliates. We have recently produced genetic evidence from *C. velia* mutant studies supporting a function of the monophyletic alveolate starch phosphorylase in the aggregation of semi-crystalline starch granules (Chabi et al. submitted). The subfunctionalization of phosphorylase in ciliates may suggest that one or several phosphorylase subunits may be devoted to this novel function while others may be devoted to the standard polysaccharide degradation function evidenced for such enzymes in fungi, animals and Amoebozoa.

**Figure 2:**
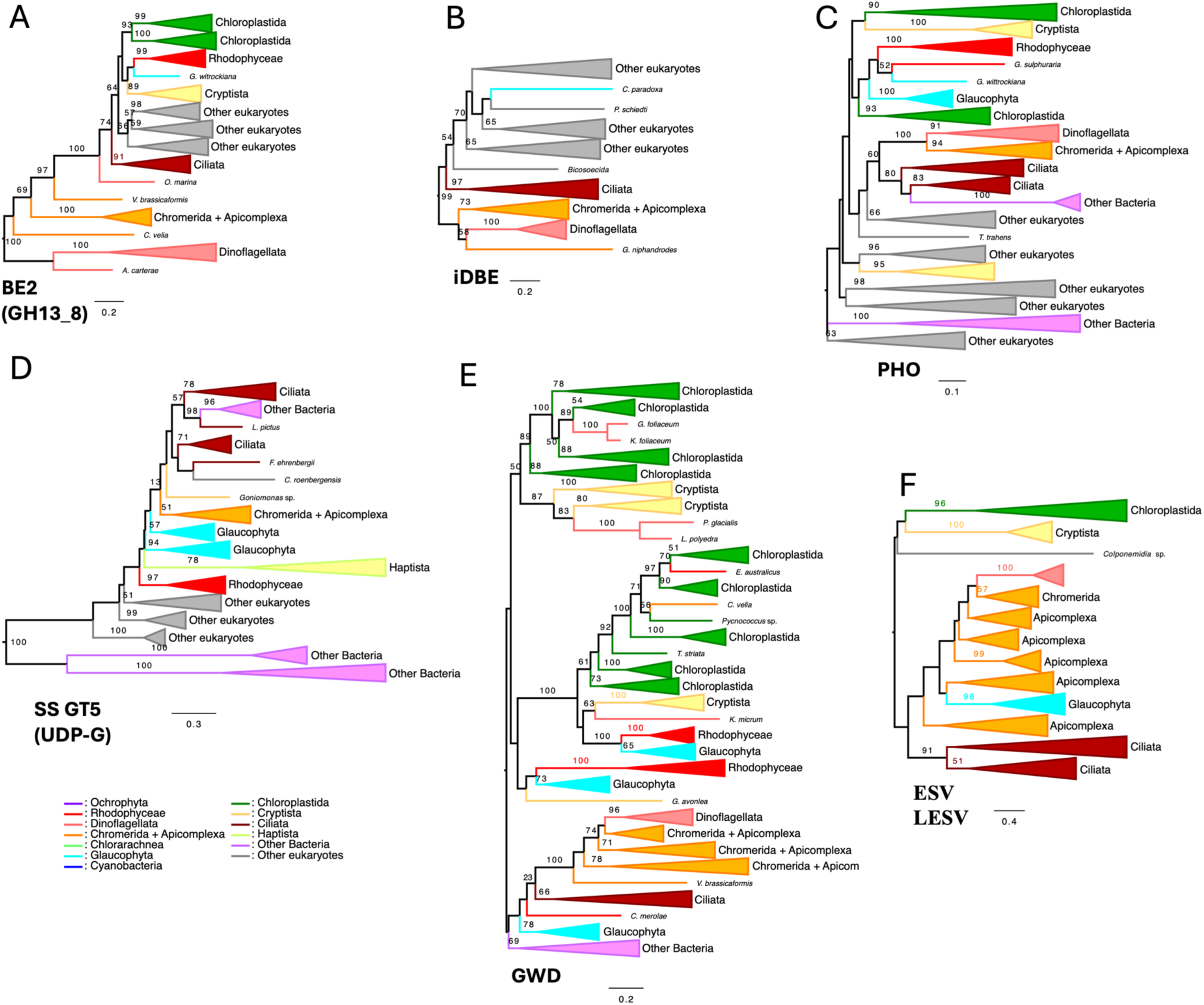
Synapomorphies in alveolate storage polysaccharide metabolism. Phylogenetic trees were all build using IQTREE with LG4X. The bootstrap values are indicated on the branch and are drawn from 100 bootstrap repetition. Sequences are colored depending on their origin with color legend indicated in the figures and the trees are midpoint rooted. (A) BE2 branching enzyme (GH13_8) is monophyletic in all alveolates including ciliates with bootstraps BS=64. (B) iDBE of all alveolate group together with (BS=54) as well (C) sequences of glycogen/starch phosphorylase (PHO) group with a BS=60. (D) while glycogen/starch synthase (SS GT5) displays a complex topology it remains compatible with a monophyletic origin of this enzyme for the ciliates and chromerid/apicomplexa albeit with very low support (BS=13). (E) GWD/PWD isoforms are hallmarks of starch metabolism separating ciliates into two groups those with dikinases synthesizing starch stand out from the simplest glycogen metabolism eukaryotic pathways, here we observe the presence of an alveolate specific GWD enzyme present in all starch accumulating alveolates. Two nodes unite the ciliates, on the one hand, and myzozoa, on another, with BS values of respectively 66 and 100 while the node uniting all of them remained low (BS=23). (F) ESV (early starvation) phylogeny confirms the absence of these polysaccharide chaperones in all red algae. Interestingly these proteins are found in Chromerids, Apicomplexa, dinoflagellates and ciliates and were likely acquired in all alveolates from Glaucophyta. In addition, we observe the presence of more distantly related ESV-LESV in Cryptophyceae which still qualify as a possible source for these genes.

In the chromerids and Apicomplexa only one isoform of the monophyletic alveolate starch phosphorylase of eukaryotic origin is found. We now document the presence of a second isoform of prokaryotic (delta-proteobacterial) origin which is selectively found in both *C. velia* and *V. brassicaformis,* but which is absent from all other alveolates including apicomplexa (Fig. S4). It is thus possible that the delta-proteobacterial isoform is taking over in chromerids some of the functions assumed by the subfunctionalized ciliate isoforms.

Within the ciliate clade, the only noticeable difference between their pathway and the enzyme suites characterizing the simplest eukaryotic glycogen metabolism pathways, are the presence of GWD/PWD isoforms (Figs. 1B, 2 and S5) as well as that of ESV/LESV proteins. We now report the phylogeny of GWD enzymes (Fig. 2E and S5). We confirm our previous finding that glucan water dikinases were lost from some late diverging ciliates that accumulate glycogen, including *Paramecium* and *Tetrahymena* (*16*) . GWD appears to be a requisite factor in starch producing eukaryotic systems, as it exhibits a perfect correlation in eukaryotes with the presence of semi-crystalline α-polyglucans. The phylogeny displayed in Fig. S3 suggests the presence of an alveolate specific GWD enzyme present in all starch accumulating alveolates. Both nodes uniting the ciliates and Myzozoa displayed BS values of respectively 66 and 100 while the node uniting all of them remained low (BS=23). Nevertheless, in a previous study with another sample of alveolates, we have obtained a BS value of 100 for the node uniting all alveolates and a BS value of 80 uniting them to glaucophytes and cyanidiales further strengthening the case for a monophyletic alveolate GWD enzyme origin (*38*).

Our phylogenetic analysis of genes encoding ESV-LESV (Fig. S6) may suggest the presence of a monophyletic alveolate specific form possibly acquired from an archaeplastidal source. Nevertheless, the support for alveolate monophyly remains low (BS=20) as is often the case in such single gene phylogenies.

ESV (early starvation) was first discovered as a starch binding protein in plants which when defective promoted deregulated starch degradation in leaves, thereby leading to a carbon starvation phenotype at the end of the night. Two related proteins were discovered (ESV and LESV (Like early starvation protein)) with similar biochemical properties (*39*, *40*) . When expressed in a yeast system reconstituting the aggregation of starch-like granules, ESV and LESV on their own did promote a significant synthesis of insoluble starch-like granular material within yeasts that otherwise still predominantly synthesize glycogen-like material. However, when expressed together with the heteromultimeric isoamylase of green plants, the recombinant yeasts synthesize significantly more glucose into starch like granules than glycogen (*41*). ESV and LESV have also been shown to interact with crystalline starch catabolism by affecting degradation of the starch crystallites through interactions with the glucan and phosphoglucan water dikinases (GWD and PWD) and through the laforin-like glucan phosphatases (*42*, *43*). ESV and LESV are non-enzymatic proteins that are considered to act as polysaccharide chaperones affecting and modelling the surface of the growing polymers (*39*). While these proteins obviously bind α−glucans, they are not as yet recognized as CAZymes as they lack previously recognized CBMs. The presence of ESV and LESV, unlike GWD, does not correlate perfectly with starch synthesis in eukaryotes as they are entirely absent from red algae. We confirm the absence of these polysaccharide chaperones in all red algae (Fig. S6). Interestingly paraglycogen (amylopectin) accumulating ciliates (Fig. S6) contain ESV-LESV while glycogen synthesizing ciliates such as Paramecium and Tetrahymena do not, thereby mirroring the distribution of GWD. Interestingly, these proteins are found in cryptophytes where they together with the cryptistan host phosphorylase (Chabi et al. submitted) presently define the only candidate protein sequences that could influence starch crystallinity (Figs. 2F and S6). Indeed, the cryptophytes apparently all lack isoamylases, pullulanases and iDBE despite the presence of iDBE in cryptista and Goniomonas (Fig. 1B, and Table S1). This suggests the presence of a novel yet to be characterized CAZyme taking over the indispensable degradation of the α-1,6 branches in these algae. Because they are not considered CAZymes we have not included ESV/LESV in our original target set of gene captures for the characterization of the starch defective mutants of *C. velia* (Chabi et al. submitted).

A striking feature of both the ciliate phosphorylase and the ciliate GT5 glycogen synthase phylogenies is the presence of two clostridia sequences nested within the ciliate phylogeny (Figs. S1, S3). Clostridia are among the very few bacteria documented to accumulate granulose which consists of semi-crystalline amylopectin granules. The diversity of clostridium strains concerned by this LGT is on the low side (two strains) and no other related clostridium sequences affiliated to ciliates could be found in the databases. The topology of the trees, the eukaryotic nature of the sequences and the low diversity of bacteria concerned argue for a transfer from ciliates to Clostridium in a common microaerophilic or anaerobic environment such as the rumen or digestive track of animals where both organisms proliferate. The documented presence of amylopectin in a majority of ciliate clades and the presence of GWD and ESV/LESV in basal ciliates and Myzozoa, argue that the last common ancestor of alveolates (LACA) was synthesizing amylopectin. In addition, the monophyletic alveolate nature of both phosphorylase and iDBE proven in *C. velia* to be responsible for starch granule aggregation (Chabi et al. submitted) further strengthens the argument.

### Five molecular synapomorphies unite Myzozoa starch metabolism

We now report five molecular synapomorphies of starch metabolism established in the last common myzozoan ancestor (LCMA) (see discussion) that distinguish them from ciliates. These synapomorphies are evidence of a considerable amount of evolution starting from the otherwise rather stable platform of storage polysaccharide metabolism witnessed in ciliates. The first two synapomorphies concern two loci encoding branching enzymes. A phylogenetic tree displaying both the prokaryotic GH13_9 (BE1) and eukaryotic GH13_8 (BE2) branching enzymes has been presented and discussed in Chabi et al (submitted). For convenience we now present two separated figures displaying a more detailed phylogeny of BE1 (Fig. 3A and S7) and BE2 (Fig. 2A and S8). A bacterial LGT encoding BE1 (GH13_9) possibly acquired from planctomycetes (Fig. S7) is fulfilling the essential branching enzyme function in Myzozoa as was initially demonstrated by targeted mutagenesis in *Toxoplasma gondii* (*44*) and subsequently confirmed by our mutant selection in *Chromera velia* (Chabi et al. submitted).

**Figure 3:**
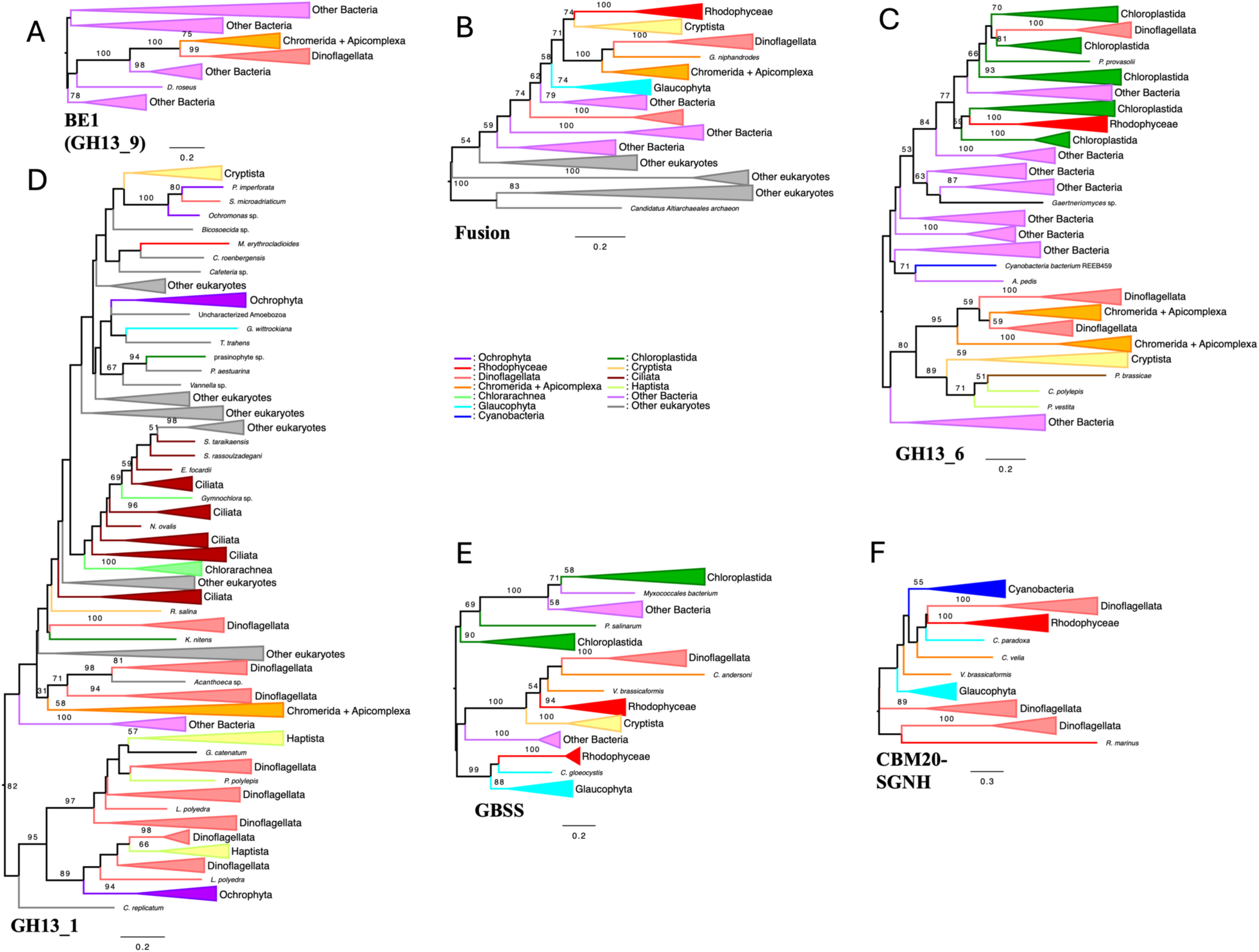
Synapomorphies of storage polysaccharide metabolism in Myzozoa. Phylogenetic trees were all build using IQTREE with LG4X. The bootstrap values are indicated on the branch and are drawn from 100 bootstraps repetition. Sequences are colored depending on their origin with color legend indicated in the figures and the trees are midpoint rooted. (A) BE1 (GH13_9) was possibly acquired from planctomycetes to fulfill the essential branching enzyme function in Myzozoa as was initially demonstrated by targeted mutagenesis in *Toxoplasma gondii* (Yang et al. 2022) and subsequently confirmed by our mutant selection in *Chromera velia* (Chabi et al submitted). (B) The fusion protein for whom we obtained multiple starchless mutations and is essential for polysaccharide synthesis (Chabi et al submitted) is composed a GT4 sequence that disrupt the BE2 (GH13_8) is found in all myzozoa synthesizing starch. The phylogeny of the GT4 part of the protein is monophyletic for all Myzozoa (BS= 100) and suspected to be either of cryptophyte or red alga (BS= 71) or glaucophyte origin (BS= 58). (C) One GH13_6 hydrolase present in all Myzozoa (BS=95) represent also a well supported synapomorphy this time from an unknown either bacterial, rhizarian, stramenopile or cryptophycean source. The enzyme concerns a GH13_6 hydrolase most probably an α-amylase not found in ciliates which was transmitted to the last common Myzozoa ancestor (LCMA). Another GH13_6 α-amylase is seen in dinoflagellates selectively and emerged from green algae (BS=66) representing an LGT unifying dinoflagellates alone. (D) GH13_1 α-amylases show that ciliates conglomerate in the same region of the tree, the nodes uniting them or a large subset of them receive very poor or no bootstrap support. Weak support (BS=31) exists for a node uniting the chromerids to the basal dinoflagellates up to the more recently diverging Kareniaceae. In addition, a well-supported node (BS=95) unites a large diversity of more recently derived dinoflagellates (from Kareniaceae onwards) with haptophytes, however, identity of the donor or recipient is unresolved. (E) The GBSS phylogeny demonstrates that a LGT from red alga or Cryptophyceae happened in the common ancestor of all Myzozoa (BS=54). (F) Finally, presence of a CBM20-containing SGNH hydrolase superfamily carbohydrate modifying enzyme, that once mutated yield starchless phenotype, qualifies as a sixth candidate synapomorphy shared by both photosynthetic dinoflagellates and chromerids within Myzozoa. The phylogeny does not resolve the gene origin of the alveolate sequences which may have received them from glaucophytes, red algae or cyanobacteria.

Both activities are closely intertwined, as the presence of BE1 correlates tightly in all starch/amylopectin synthesizing Myzozoa with a disruption caused by an insertion of a GT4 type of glycosyl transferase within the previous common monophyletic alveolate BE2 gene (GH13_8) (Figs. 2A and S8). The fusion product has thus become unable to complement a selective defect in BE1 generated in either the Toxoplasma or Chromera BE1 mutants (Chabi et al submitted). The disruption by the GT4 transferase effectively separates the function of the N-terminal part of the BE2 protein including its CBM48 (**C**arbohydrate **B**inding **M**odule of CAZy family 48) from the classical branching enzyme activity (Fig. 2A). Indeed, multiple starchless mutations spanning all regions of the BE2/GT4 fusion have been recently selected in *Chromera velia* (Chabi et al. submitted), proving that the BE2/GT4 fusion is essential for polysaccharide synthesis in chromerids and probably all Myzozoa. However, a particular nonsense mutation allowing to produce a truncated protein consisting of the CBM48 and the first few residues of the BE2 enzyme still allows for the synthesis of up to 20% of the wild-type amount of normal starch granules in *C. velia* (Chabi et al. submitted). This occurs despite the complete absence of the GT4 transferase and the likely absence of catalytic BE2 branching activity. We thus surmise that there is a previously unsuspected and unknown essential function of polysaccharide synthesis encoded at the N-terminus of BE2 and that the BE2 and GT4 domains are regulators within this complex gene fusion which through their folding and substrate sensing either prevent or allow for the proper presentation of this essential part of the protein. The prokaryotic BE1 gene does not contain such an essential CBM48-containing region, at its N-terminus (Chabi et al. submitted) and therefore cannot either complement this function of the BE2α-GT4-BE2b gene fusion in the BE2 fusion mutants selected in *Chromera velia*. Both BE1 and BE2 have thus become indispensable to ensure significant starch synthesis in Myzozoa. This rearrangement of branching enzyme genes and function is found in possibly all starch/amylopectin synthesizing Myzozoa including chromerids, apicomplexa and dinoflagellates while it is never found in ciliates. The phylogeny of the GT4 enzyme is presented in Figs. 3B and S9. As suggested by this analysis the alveolate GT4 insertion is suspected to be either of cryptophyte or red alga (BS= 71) or glaucophyte origin (BS= 58).

Another well supported synapomorphy (BS=95, Figs. 3C and S10) common to all Myzozoa also entails an LGT, this time from an unknown either bacterial, rhizarian, haptophyte or cryptophycean source (Fig. S10). The enzyme concerns a GH13_6 hydrolase most probably an α-amylase not found in ciliates which was transmitted to the last common Myzozoa ancestor (LCMA). α-amylase genes are often exchanged between eukaryotic phagotrophs and between the latter and bacteria leading to patchy gene distributions within the eukaryotic tree. Another GH13_6 α-amylase is seen uniting a diverse set of dinoflagellates with green algae (Figs. 3C and S10). These will be further discussed below. To better understand the fate of the ciliate GH13_1 α-amylases we now report this enzyme’s phylogeny (Figs. 3D and S11). In this case while ciliates conglomerate in the same region of the tree, the nodes uniting them or a large subset of them receive very poor or no bootstrap support. Some weak support (BS=31) exists for a node uniting the chromerids to the basal heterotrophic dinoflagellates up to the more recently diverging Kareniaceae which are nevertheless considered by some authors to be among the earliest diverging photosynthetic dinoflagellates (*11*). However, the BS values are too low and the diversity too restricted with no such sequences detected in the apicomplexa to propose this as a synapomorphy for Myzozoa. Nevertheless, a well-supported node (BS=95) unites a large diversity of more recently derived dinoflagellates (from Kareniaceae onwards) with expected haptophytes (*45*). However, it is impossible to propose the identity of the donor or recipient clade as both the haptophyte and dinoflagellate diversity are mixed in a way suggesting multiple gene exchanges between them.

GBSS, the enzyme responsible for amylose synthesis in all species tested, appears to define a fourth synapomorphy uniting all Myzozoa although in both Apicomplexa and chromerids the distribution of this gene is rather patchy with only *Vitrella brassicaformis* and several Cryptosporidia species having been demonstrated to contain such sequences (*16*) within the chromerid/apicomplexa clade, while the gene seems much more widely distributed in dinoflagellates. The topology of the GBSS phylogeny nevertheless demonstrates that a red alga GBSS LGT was common to the evolution of all Myzozoa (Fig. 3E and S12, (BS=54)). To further nail down the correlation of amylose biosynthesis with the presence of GBSS in alveolates we have purified the storage polysaccharide from both Chromera and Vitrella (Fig. 3 and S13). As expected, we find substantial levels of amylose in the very anciently derived chromerid Vitrella and none in Chromera. Cryptosporidia polysaccharides on the other hand have been reported to yield a strong blue stain with iodine which is characteristic for amylose containing starches (*46*). The electron micrographs previously published for *V. brassicaformis* show a very tight association of starch with the Vitrella pyrenoid across the four membranes of the secondary plastid (*47*). This is clearly evidencing starch as a major sink for photosynthate export in the chromerid cytosol. The pyrenoid starch association resembles a similar arrangement found in mesophilic unicellular red algae such as Rhodella with the stromal rhodoplast pyrenoid bulging off the plastid surrounded by starch on the cytosolic side, across the 2 rhodoplast membranes (*48*, *49*). Cryptophyceae have conserved the endosymbiont (red algae) type of association in the periplast and have lost starch metabolism in the host cytosol. We have previously hypothesized the presence of amylose as necessary for obtaining the morphogenetic plasticity of starch that is required to surround the pyrenoid (*14*, *50*). GBSS in all Archaeplastida, Cyanobacteria, Cryptophyta and *Vitrella* displays a conserved N-terminus and a molecular mass around 60 kDa (Figs. 3E and S14). Interestingly dinoflagellates depart from this consensus by displaying N-terminal extensions of variable lengths and sequences.

Finally, the presence of the CBM20-containing SGNH hydrolase superfamily carbohydrate modifying enzyme, qualifies as a fifth candidate synapomorphy shared by both photosynthetic dinoflagellates and chromerids within Myzozoa. Nevertheless, several events of gain and losses remain compatible with the observed phylogeny due to the low resolution of such single gene trees. In that case it would not be a true synapomorphy. This enzyme first identified by our mutant selection in *C. velia* could possibly define a regulatory protein affecting cytosolic starch metabolism in the presence of photosynthesizing plastids. (Figs. 3F and S15) The phylogeny does not resolve the gene origin of the alveolate sequences which may have received them from glaucophytes, red algae or cyanobacteria

### Two candidates for molecular synapomorphies and five enzyme additions of starch metabolism unite dinoflagellates within Myzozoa

The presence of numerous complete full genome sequences in both the Apicomlexa and sister chromerid lineages solidify our above conclusions concerning the myzozan synapomorphies. However, both the gargantuan size of dinoflagellate genomes and the heterotrophic often parasitic nature of the earliest diverging clades do not allow us the same level of confidence. Hence, we will use the term candidate synapomorphy whenever a reasonable sample encompassing the full diversity of available dinoflagellate sequences including the earliest diverging clades support it. We will use the term enzyme additions or acquisition to cover all other situations which will possibly translate into established synapomorphies as the databases expand.

The dinoflagellates have acquired seven additional enzymes affecting storage polysaccharide metabolism which are absent from all other alveolates. The first dinoflagellate synapomorphy concerns the replacement of the ancient unique eukaryotic UDP-Glc-utilizing GT-5 glycogen synthase found in many glycogen accumulating protists including alveolates by several isoforms of a novel glucan synthase, each probably displaying a distinct biochemical function. We have previously documented 5 analogous enzyme isoforms during the annotation of the first completed cryptophyte genome sequence (*16*, *51*). These are the only candidate glycosyl-transferases possibly involved in elongating glucan chains from UDP-Glc presently found in cryptophyte or dinoflagellate genomes save for GBSS whose function is confined to amylose synthesis. In addition, we have previously reported mutants of the heterotrophic dinoflagellate *C. cohnii* that display a reduced and modified soluble starch synthase activity (*52*). This modified soluble starch synthase phenotype cosegregates in crosses with low starch accumulation establishing that these enzymes are indeed controlling starch polysaccharide elongation from UDP-Glc in all dinoflagellates. In all past and present phylogenies since their discovery in 2012, these enzymes appeared united with GBSSI and SSI/SSII from green algae with reasonable bootstrap (BS=68) (Figs. 4A, S16). While we now show the whole GT5 family diversity for the reader’s convenience, we have previously published more detailed phylogenies encompassing more sequence diversity from both cryptophytes and dinoflagellates (*53*). While the interrelation of GBSS SSI/SSII and the cryptophytes or dinoflagellates are not resolved as is often the case in such deep time single gene phylogenies, we can rely on the dating of the nodes uniting all green algae, cryptophytes and dinoflagellates to propose LGT scenarios. Indeed, there is a general agreement, despite ongoing controversies, on the antiquity of primary endosymbiosis. This is dated around 1.5 billion years which is non overlapping with all proposed dates for secondary or higher order plastid endosymbiosis events (*54*). Because all green algae and all plants with no reported exceptions contain SSI, SSII and GBSS we can confidently exclude that the robust nodes defining the phylogeny of these three enzymes are the result of LGTs from either cryptophytes or dinoflagellates as donors. The green lineage must thus be the donor. Three possible scenarios can be nevertheless proposed. In a first scenario, both cryptophytes and dinoflagellates have received independently a copy of a related green alga source. In a second scenario the cryptophytes have received the green gene and in turn donated it to dinoflagellates while in a third scenario the dinoflagellates have selected the green gene LGT and passed it on to cryptophytes. Among the 3 possible sources (GBSS, SSI or SSII) of the green starch synthase genes we have suggested that the anyhow more distantly-related GBSS is the less likely donor. Caution might nevertheless be required before discarding GBSS as there is a suspicion that at least in green algae SSI-SSII which have no close prokaryotic relatives enzymes have themselves evolved from an ancestral archaeplastidal GBSS (*22*).

**Figure 4:**
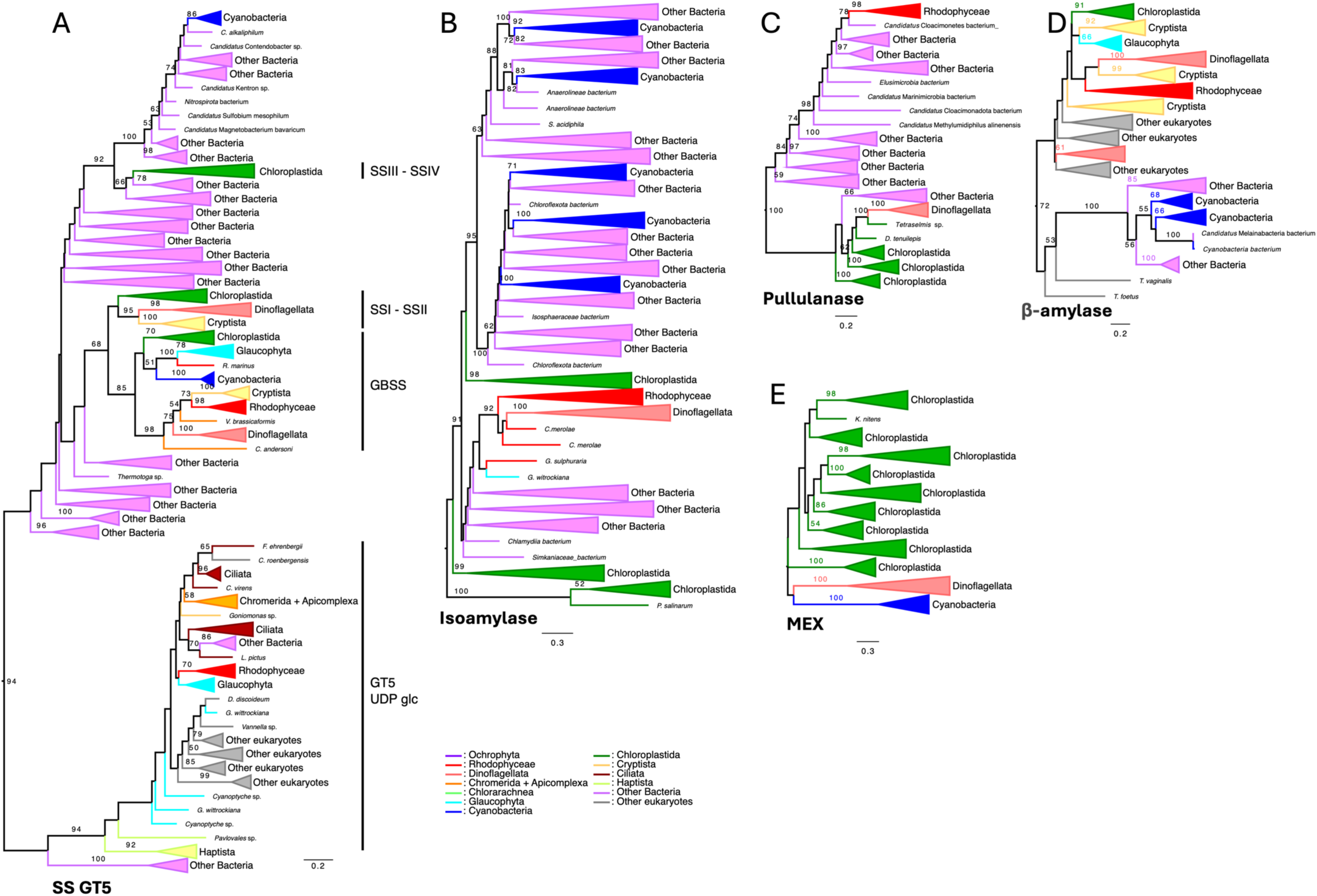
Synapomorphies of storage polysaccharide metabolism in dinoflagellates. Phylogenetic trees were all build using IQTREE with LG4X. The bootstrap values are indicated on the branch and are drawn from 100 bootstraps repetition. Sequences are coloured depending on their origin with colour legend indicated in the figures and the trees are midpoint rooted. (A) In dinoflagellate we already reported presence of SSI/SSII enzyme close to GBSS SSI and SSII sequences from green algae origin (Nielsen et al. 2018) and display here again their phylogeny with reasonable bootstrap (BS=68). The interrelation of GBSS SSI/SSII and the cryptophytes or dinoflagellates is not resolved. Among the 3 possible sources (GBSS, SSI or SSII) of the green starch synthase genes we suggest that the anyhow more distantly-related GBSS is the less likely donor. (B) An isoamylase gene (CAZy subfamily GH13_11) displays a robust node uniting them to cyanidiales and other red algae in our reported phylogenies (BS=92), despite this we cannot rule out completely an unlikely glaucophyte source for this originally prokaryotic gene. This indicates a direction of transfer from Archaeplastida to dinoflagellates. (C) A second green gene encoding pullulanase (GH13_14) defines a third synapomorphy with dinoflagellates grouping with the green algae Tetraselmis with a BS=100. (D) A fourth synapomorphy uniting dinoflagellates within alveolates consists of the acquisition of β-amylase. One set of sequences probably results from exchanges that occurred between eukaryotic phagotrophs while the other set is united as sisters to cryptophytes with low support (BS=37). Both cryptophytes and dinoflagellates are united to red algae with very low support (BS=17). (E) MEX1 (maltose export) is known in archaeplastida to export maltose from the chloroplast to the cytosol. MEX1 acquisition defines the fifth synapomorphy that distinguishes starch metabolism in dinoflagellates from other alveolates.

The finding of the “green” SSI/SSII like starch synthases spanning the whole diversity of dinoflagellates including the earliest diverging *O. marina* qualify these enzymes as a candidate dinoflagellate synapomorphy.

Strikingly we also find a gene fusion between the common alveolate BE2 branching enzyme (see above) and the dinoflagellate SSI/SSII starch synthase. The finding of this gene fusion with a similar domain organization in both the heterotrophic and early diverging *O. marina* and the later diverging photosynthetic *P. glacialis* is further backed by the detection of comparable fusions in a wide diversity of dinoflagellates (Table S1). The fusion consists of an N-terminal BE2 full enzyme sequence complete with its CBM48 N-terminal regions that we have proven essential for starch synthesis in Myzozoa, (Chabi et al. submitted) followed without disruption of the BE2 sequence by the GT5 green SSI/SSII. We found that dinoflagellates carry both the BE2 fused and unfused forms of the GT5 green SSI-SSII starch synthase. Dinoflagellate strains were also found that contain both the BE2 GT4 and GT5 fusions. Because of the wide distribution of the BE2-GT5 gene fusion within dinoflagellates including the earliest branching *Oxyrrhis marina* we can reasonably propose that this defines a second robust synapomorphy of dinoflagellates. Interestingly we were unable to detect analogous fusions in cryptophytes nor were we able to detect fusions between the cryptophyte branching enzyme and the cryptophyte homolog of the GT4 transferase.

A third enzyme addition witnessed in diverse dinoflagellates consists of the acquisition of an isoamylase gene (CAZy subfamily GH13_11). The latter displays a robust node uniting them to cyanidiales and other red algae in both of our reported phylogenies (BS=92 and BS=95) (Figs. 4B, S17 and (*29*). Despite this we cannot rule out completely an unlikely glaucophyte source for this originally prokaryotic gene. Anyhow, because of its restricted distribution in eukaryotes, we can safely assume that the direction of transfer is from Archaeplastida to dinoflagellates. The requirement for isoamylase comes as a surprise in face of the presence in dinoflagellates of the very same phosphorylase and iDBE isoforms that we have recently proven to be involved in amylopectin crystallization in chromerids (Chabi et al. submitted). As discussed below, the model we propose for storage polysaccharide metabolism evolution fully accounts for this surprising result. The distribution of isoamylase in dinoflagellates is wide and includes both Amphidinium and Kareniaceae (*11*, *55*) which according to different authors define the earliest diverging photosynthetic dinoflagellates carrying the classic peridinin plastid. We have presently not found isoamylase sequences in the earliest diverging heterotrophic dinoflagellates. We believe this to be due to lower sequence sampling in these early diverging clades and suspect that isoamylase will likely translate in the future into a dinoflagellate synapomorphy

A fourth enzyme acquisition is a second green gene encoding pullulanase (GH13_14) acquired by LGT. Pullulanase as defined in Fig. 1C is a prokaryote-specific direct debranching enzyme not found in eukaryotic glycogen metabolism networks which like isoamylase directly hydrolyses the α-1,6 linkage branch thereby releasing MOS (maltooligosaccharides) in the cytosol of red algae and in the plastid stroma of green algae and plants. While pullulanase catalyses the same basic reaction as isoamylase, it acts faster on tightly spaced branches and is comparatively less effective on the loosely spaced branches of glycogen which are the preferred substrates of isoamylase. It is thought that pullulanase (also called limit-dextrinase) is an important enzyme of starch catabolism in green algae and plants and may modulate the action of isoamylase during synthesis (*31*). Interestingly in Archaeplastida pullulanase has been transmitted twice independently in the common ancestors of respectively the red and green algae from distinct bacterial sources (*16*). The phylogeny of the dinoflagellate pullulanase presented in Figs. 4C and S18 proves both that the dinoflagellates have acquired the pullulanase gene from early diverging green algae and that the direction of transfer was from green algae to dinoflagellates. We infer that selection of a pullulanase LGT may have been triggered by the prior transfer of isoamylase. In the present restricted sample of dinoflagellate sequences (mostly Suessiales), we were nevertheless able to find species that encoded all three debranching enzymes: the eukaryotic indirect debranching enzyme and the prokaryotic isoamylases and pullulanases. Hence, we believe there is no or little functional overlap and redundancy between these enzymes.

A fifth enzyme addition of dinoflagellates within alveolates consists of the acquisition of β-amylase. This “phagotrophic” enzyme mostly distributed in the eukaryotic domain is shared by many glycogen-accumulating heterotrophic eukaryotes such as Amoebozoa or Metamonada and occasionally by phagotrophs lacking glycogen metabolism such as Naegleria. β-amylase (GH14) is an exo-type of enzyme producing β-maltose from the non-reducing end of an outer chain of glycogen or amylopectin. Maltose accumulation is cytotoxic, especially in the osmotically sensitive eukaryotic cytosol. The phylogenetic tree presented in Fig. 4D and S19 characteristically lacks bacteria and fungi. Close inspection of the tree emphasizes the existence of two groups of dinoflagellate β-amylase sequences. One set of sequences probably results from exchanges that occurred between eukaryotic phagotrophs while the other set is united as sisters to cryptophytes with low support (BS=37). Both cryptophytes and dinoflagellates are united to red algae with very low support (BS=17). Because of the antiquity of the node uniting a large section of the red alga diversity including Cyanidiales, Bangiales, Florideophycidae, and mesophilic unicellular red algae, and because of the topology of the tree we can propose a mesophilic red alga as donor of the cryptophyte/dinoflagellate genes. However, three scenarios are possible. Both dinoflagellates and cryptophytes may have received a copy of this red β-amylase gene independently. Alternatively, the cryptophytes may have received the gene from red algae and transferred it to dinoflagellates, or the dinoflagellates may have received the red alga gene and passed it on to cryptophytes. Both β-amylase forms are widely distributed in dinoflagellates including some of the earliest diverging photosynthetic clades such as Karlodinium. As for isoamylase, we suspect this enzyme gain may translate into a dinoflagellate synapomorphy in the future upon further sampling of very early diverging heterotrophic dinoflagellates.

The Archaeplastida use the DPE2 enzyme (amylomaltase (GH77)) to achieve maltose degradation exclusively in the cytosol. Yet, in green algae and plants, β-amylase degrades starch granules in conjunction with GWD and PWD dikinases in the chloroplasts that typically lack DPE2 activities despite containing another α-1,4 GH77 enzyme unable to process maltose (DPE1). Green algae and plants have solved this problem by evolving a novel plastidial membrane transporter called MEX1 (maltose export or maltose excess) which exports maltose from the chloroplast to the cytosol for degradation by the DPE2 cytosolic amylomaltase. MEX1 acquisition defines the sixth enzyme acquisition that distinguishes starch metabolism in dinoflagellates from other alveolates. The phylogeny displayed in Figs. 4E and S20 does not resolve the origin of the dinoflagellate gene. Nevertheless, the presence of the whole diversity of the green algae and plants enables us to reasonably reject the possibility that the dinoflagellates have donated the gene to the Chloroplastida. The proximity of the dinoflagellates relatively to the green lineage and the restricted diversity and type of cyanobacteria present in the tree does not favour a cyanobacterial origin for the MEX1 gene. We therefore prefer the likely scenario that the dinoflagellates have acquired this transporter from the green algae. Interestingly MEX1 and both β-amylase isoforms exhibit a similar type of wide taxonomic distribution within dinoflagellates including some of the earliest diverging photosynthetic clades which is consistent with their function

Finally, the presence of an isoform of GH13_6 α-amylase of green alga origin (Figs. 3C and S10) is found selectively in dinoflagellates in addition to the common myzozoan isoform, thereby defining a possible seventh enzyme acquisition specific of dinoflagellates that include some early diverging photosynthetic clades

### Acquisition of DPE1-like α-1,4 glucanotransferase distinguishes the chromerids/Apicomplexa from dinoflagellates

By comparison to the dinoflagellates, synapomorphies of starch metabolism that distinguish the chromerids relatively to the dinoflagellates are scarce. The only addition to the starch network consists of DPE1 (DisProportionating Enzyme 1) (Fig. 5 and S21). This enzyme was named inappropriately after the form of enzyme present in the chloroplast of green algae and plants. Plant DPE1 transfers segments of glucan donor chains to recipient glucans, occasionally releasing glucose in the process. The plant enzyme is unable to use the disaccharide maltose as donor for transfers to recipient oligosaccharides, despite the latter being generated in abundance by both α- and β-amylases within the chloroplast. DPE2 (amylomaltase) can do so in the cytosol thanks to the export of maltose from the chloroplast by the MEX1 transporter. In apicomplexans and chromerids an α-1,4 glucanotransferase candidate gene named DPE1 (CAZy GH77) was found in several amylopectin accumulating Apicomplexa and *Vitrella* but has not been found in *Chromera* or *Cryptosporidium*. It is unclear whether this enzyme can break down maltose.

**Figure 5:**
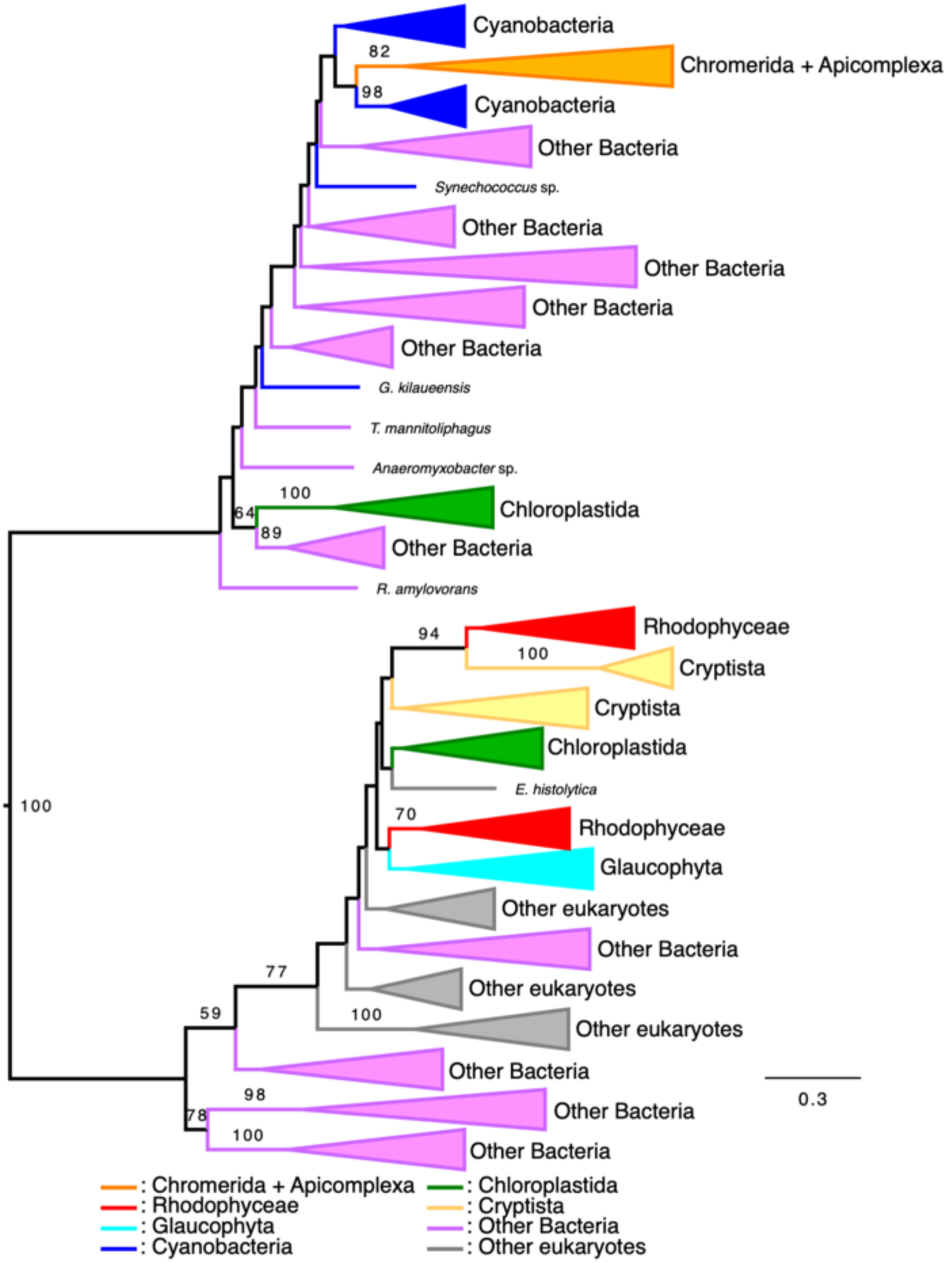
Phylogeny DPE1. Phylogenetic trees were all build using IQTREE with LG4X. The bootstrap values are indicated on the branch and are drawn from 100 bootstraps repetition. Sequences are colored depending on their origin with color legend indicated in the figures and the trees are midpoint rooted. Amylomaltase DPE1-2 (GH77) is the only case involved in starch metabolism that distinguish the chromerids/apicomplexa from the dinoflagellates. We display here the phylogeny of apicomplexa and chromerids α -1,4 glucanotransferase candidate gene named DPE1 (CAZy GH77) found in several amylopectin accumulating apicomplexa and Vitrella. Despite baring the same name as the plant and green alga enzyme which cannot breakdown maltose, the chromerid/apicomplexa enzyme appears more distantly related to such enzymes with a possible affiliation to cyanobacteria. We thus have no reason to suspect that chromerid enzyme has or has not the ability to digest maltose.

### Comparative biochemistry of starch biosynthesis in lineages derived from secondary endosymbiosis of red algae

Among lineages suspected to be derived from secondary endosymbiosis of both red and green algae (*56*) only the red alga secondary endosymbiosis that generated cryptomonads and the secondary or higher order founding endosymbiosis that possibly led to the Myzozoa have maintained complete starch metabolism pathways and have been demonstrated to lead to the accumulation of substantial amounts of this form of polysaccharide (*22*). In Cryptophyceae (photosynthetic cryptomonads) starch is found exclusively in the periplast of the secondary plastid with essentially no polysaccharides left in the secondary endosymbiosis host cytosol (*57*, *58*) . *Goniomonas avonlea* can be distinguished from other heterotrophic cryptomonads by the absence of plastid (*38*, *59*). Genomic sequencing and analysis demonstrate that *G. avonlea* contains a complete and complex pathway of glycogen metabolism including GS (glycogen synthase) of both the GT3 and GT5 forms of the enzyme, glycogen branching enzyme, glycogen phosphorylase, the autoglucosing elongation priming protein glycogenin, indirect debranching enzyme, alpha and beta amylases, the DPE2 maltase and various amylase-like hydrolases (Fig 1B, Fig 1C, Sup Table 2). Most importantly it contains genes encoding glucan water dikinases (GWD). Since the presence of GWD correlates with that of crystalline amylopectin in eukaryotes, we can suggest that Goniomonas synthesizes cytosolic amylopectin granules in yet to be reported specific physiological conditions or in particular stages of its life cycle. We therefore propose that the Goniomonas pathway reflects the status of storage polysaccharide metabolism that existed prior to the secondary red alga endosymbiosis of cryptomonads. By analogy, we believe that the status of storage polysaccharide of LCMA before secondary or tertiary endosymbiosis will be similar to that evidenced in early diverging ciliates.

**Table 2:** Periplastidial starch metabolism network of the cryptophyte *Guillardia theta*.

| ABBREVIATION | ENZYME NAME | N° | GENE<br>TRANSFER | GENE SOURCE | PHYLOGENY |
| --- | --- | --- | --- | --- | --- |
| **SSI-SSII (UDPG)** | Starch Synthase | 5 | LGT | Green algae or Dinoflagellates | **Figure S15** |
| **GLG** | Glycogenin | 2 | - | Host | Not shown |
| **BE(GH13_8)** | Branching enzyme | 1 | - | Host | **Figure S1** |
| **GBSS** | Granule-bound starch synthase | 1 | LGT or EGT | Red algae (EGT) or Myzozoa (LGT) | **Figure S10** |
| **A-AMY (GH13_6)** | α-amylase | 1 | LGT | Bacteria or Myzozoa | **Figure S8** |
| **A-AMY (GH13_10)** | α-amylase | 1 | LGT ? | Green algae | Not shown |
| **A-AMY (GH13_32)** | α-amylase | 1 | LGT | Dinoflagellates/Bacteria | Not shown |
| **PHO (GT35)** | Starch Phosphorylase | 2 | LGT and - | Host and likely green alga (unlikely red) | **Figure S3** |
| **β-AMY (GH14)** | β-amylase | 3 | LGT or EGT or - | 1 red algae (EGT) or LGT (Dinoflagellates)) + 1 green alga or Glaucophyte (LGT) +1 Host or diverse protists (LGT) | **Figure S18** |
| **DPE2** | α-1,4 glucanotransferase | 2 | EGT + LGT | 1 red alga (EGT) + 1 unresolved (LGT) | **Figure S20** |
| **GWD-PWD** | Glucose/phosphoglucose dikinase | 3 | EGT and LGT | Red algae (EGT) and green algae LGT | **Figure S5 and Cenci et al. 2018)** |
| **ESV_LESV** | Early starvation protein | 1 | LGT | Green algae (LGT) | **Figure S13** |

The number of distinct enzyme isoforms is listed in the “N°” column. A mosaic of genes of either red (EGT) or green alga (LGT) or host origin are evidenced. In many (7) cases the low resolution of these single gene trees offers multiple possible gene sources which are all listed. ? indicates possible, yet with uncertain or complicated determination. – in gene transfer column indicates a host origin.

Ciliates, however, lack both β-amylase and their associated DPE2 maltase as well as the GT3 GS which are all present in Goniomonas suggesting that the last common ancestor of either Alveolata or Cryptophyta may indeed be different in this respect. They however also contain glucan water dikinases which is never seen in glycogen accumulating organisms. Hence both heterotophic hosts of secondary (or tertiary) endosymbiosis were already amylopectin synthesizing protists prior to endosymbiosis and thus must have evolved mechanisms that allowed for the crystallization and aggregation of such polysaccharides.

Thanks to the phylogenetic analysis performed in the present work and access to the full genome sequences of both Goniomonas and its photosynthetic derivatives, we can now retrace the events that led to periplastidial starch metabolism enzyme network in cryptophytes. The phylogenies of the enzyme network listed in Fig. 1B and table 2 further prove that cryptophytes are using the red alga’s blueprint of starch metabolism. Yet only a minority of enzymes were recruited as summarized in Table 2 to build upon this blueprint directly from the red alga network (possibly only GBSS one isoform of β-amylase and one isoform of the DPE2 amylomaltase). Several enzymes were also recruited from the host network of cytosolic amylopectin synthesis including at minimum branching enzyme, starch phosphorylase and glycogenins (Table 2). Suggestive evidence for LGTs from either Myzozoa, dinoflagellates, green algae and bacteria has also been gathered (Table 2). Among such LGTs the most surprising features consist in the recruitment of a new type of soluble starch synthase which is abundantly discussed below and the recruitment of ESV/LESV both from a likely green alga source. The ESV gene was lacking from the original starch metabolism network of the red alga symbiont. It was required because of the absence of direct or indirect debranching enzymes in the cryptophyte periplast. By analogy with the function of phosphorylase reported for alveolates (Chabi et al. submitted) the only two genes that are likely candidates to explain the aggregation of semi-crystalline starch granules in cryptophytes consist of the isoform of phosphorylase donated by the cryptistan amylopectin accumulating host and by ESV.

The net result of the impact of secondary endosymbiosis on starch metabolism is a chimeric network of enzymes substituting for the ancient rhodophycean pathway. Our most intriguing finding remains the apparent absence of enzymes documented to be able to hydrolyse the α-1,6 linkage such as isoamylase, pullulanase or indirect debranching enzyme that we confirm in this study, over a decade after first reporting it. Yet the branch-linkage must be catabolized somehow. It is entirely possible that some of the GH13 enzyme subfamilies designated as α-amylase on the basis of a few detailed biochemical literature reports may in fact harbor the missing enzyme(s).

## DISCUSSION

### Presence of a starch accumulating symbiont as the dinoflagellates diverged from other Myzozoa

We have shown that the dinoflagellates can be distinguished by the possession of genes for the SSI/SSII/GBSS subgroup of starch synthases found in green algae and plants (Fig 4 and S16). An analogous form of enzyme was first documented in the cryptophyte *G. theta* (*51*, *53*, *57*, *58*) and is seen in Fig. S16 in sister photosynthetic cryptophyte lineages. All other alveolate lineages synthesizing storage polysaccharides display glycogen/starch synthases belonging to the GT5 CAZy family commonly found in the eukaryotic cytosol where they participate in glycogen synthesis from UDP-Glc (Fig. S1, S16). Interestingly the heterotrophic plastid-less cryptistan Goniomonas has both standard eukaryotic GS (<u>g</u>lycogen synthase) the aforementioned GT5 and the GT3 GS which is the predominant form of GS found in animals and fungi with no traces of the SSI/SSII-like enzyme of its photosynthetic plastid bearing relatives (Fig. 1B, 4 and S16). This further strengthens the previous suggestion that Goniomonas ancestors may not have experienced the secondary red algal endosymbiosis that generated its photosynthetic cryptophyte relatives (*38*, *59*). That the SSI/SSII-like starch synthases from dinoflagellates and cryptophytes also use UDP-Glc is suggested by our previous description of low starch mutants of the heterotrophic dinoflagellate *C. cohnii* (*52*) with altered starch synthase activity as well as our biochemical studies on starch metabolism in *G. theta* (*57*, *58*) and *Crypthecodinium* (*60*). Dinoflagellate SSI/SSII like starch synthase cluster possesses a great diversity of clades from the most basal heterotrophic clades recognized as dinoflagellates to the more recently derived photosynthetic clades. This detection may have been facilitated by the likely presence of several isoforms of this enzyme as we reported initially for the cryptophytes (*51*). The surprising presence of such isoforms can be explained as follows.

During primary and secondary plastid endosymbiosis, as the symbiont is isolated from its free-living relatives, symbiont genes will tend to accumulate deleterious mutations despite natural selection, a process known as Mueller’s ratchet (*61*, *62*). This will lead to fitness erosion of the gene because of the absence of recombination with the symbiont’s free-living sisters and the ensuing absence of possible selection of the best possible genotype afforded by the natural free-living population. EGT, a process by which a duplicated symbiont gene will be integrated into the host nuclear genome and its protein product targeted back to the symbiont, opposes this fitness erosion. Residing in the host genome, the transferred compromised symbiont gene is no longer subjected to the deleterious effects of Mueller’s ratchet as it benefits from the repair and recombination processes adapted to the host’s genes and natural population size (*62*). The protein product will have no problems expressing its function in a compartment where it had been preselected and the original plastidial copy of the gene can thus be readily lost. EGT of cyanobacterial genes can be observed in Archaeplastida while EGTs of red algae or green algae genes in the host of green and red alga secondary endosymbioses is also observed.

However, EGTs can be impeded by both extrinsic and intrinsic factors. First and foremost EGTs are readily generated by cellular processes such as autophagy that lead to the random insertion of large pieces of organelle DNA in the nuclear genome of the host (*63*). However, this recycling of organelles is expected to lead to catastrophic consequences if the secondary or primary endosymbiosis has subsequently generated an organism that has only a single copy of the organelle which thus cannot be subject to autophagy without leading to its loss(*64*). This is however the case for both the green alga *C. reinhardtii* and cryptophyte plastids (*51*, *65*, *66*). Significantly, unlike higher plants where multiple plastids per cell are generally observed, very little or no plastid DNA-derived sequences (NUPTs) are observed in the nuclear genome of the host of both Chlamydomonas and *G. theta* while these are abundant in plants(*51*, *65*). In addition, no copies of nucleomorph derived sequences could be observed in the host nuclear genome of Guillardia further suggesting that EGTs are no longer possible in this system thereby explaining the presence of nucleomorphs which are nuclei relics of the ancient red alga symbiont. The “limited transfer window’ hypothesis” described above which explains EGTs arrest can be considered as an extrinsic factor impeding EGT, as it applies to all ratchet-compromised symbiont proteins irrespective of their nature (*64*). Equally important are intrinsic factors which prevent a minor yet significant portion of compromised symbiont functions to be transferred to the host nucleus. Several variable functional or structural reasons may prevent useful genes to be expressed from the host nucleus and their product targeted to the plastid making them unsuitable candidates for becoming EGTs (*62*, *67*, *68*). Among these are their intrinsic ability or inability to follow the available routes of protein targeting. In the specific case of starch metabolism, it is of relevance that enzymes of starch metabolism of green algae are localized in the chloroplast of these algae while all enzymes of starch metabolism of red algae and glaucophytes are located in the cytosol and have no past record of being able to follow the plastid targeting route (*14*, *21*). They have thus not been subjected to selection for their ability to be taken in charge by chaperones and subsequently renatured in a foreign environment. In particular, the UDP-Glc requiring GT5 glycogen/starch synthase is an enzyme of ancient eukaryotic ancestry(*22*). Unlike other enzymes of the ancestral common cytosolic archaeplastidal starch metabolism, it has not been observed to be targeted to plastids during evolution of the chloroplastida (for a review concerning this issue see (*21*)). Thus, the GT5 starch synthase present in the cytosol of all red algae has likewise failed to generate an EGT. Whenever a compromised symbiont function cannot be rescued by EGTs other solutions may be sought. One of these for the cryptophytes would be for natural selection to recruit an enzyme of similar function from the heterotrophic starch accumulating cryptistan phagotroph host ancestor exemplified here by Goniomonas (*38*). However, both GT3 and GT5 glycogen/starch synthase present in this organism are equally unlikely substrates for plastid targeting. Yet, another possibility would be to recruit a foreign gene by LGT. A green gene encoding either SSI SSII or GBSS from a green alga was thus selected to be targeted to the symbiont (*53*). These genes have themselves evolved from an ancient GBSS gene when the starch pathway was progressively redirected to the chloroplast in the rising green algae (*21*). The glaucophyte GBSS enzyme can efficiently accommodate both UDP-Glc and ADP-Glc(*53*). It is reasonable to assume that this recruitment afforded for the evolution of a UDP-Glc accommodating enzyme from an initially ADP-Glc requiring starch synthase from green algae. The soluble starch synthases of green algae and plants are only partly redundant and accomplish mostly specialized functions, such as those involved in the seeding of novel starch granules, the priming of polysaccharide synthesis, or the synthesis of very small medium or long glucan chains(*69*). The first enzyme that came in, to assist the unique rhodophycean enzyme present in the periplast of the cryptophyte symbiont probably was selected to complement one of the most ratchet compromised functions fulfilled by the unique multiple purpose red alga starch synthase(*51*). After adapting to the use of UDP-Glc the gene encoding this novel enzyme was likely duplicated and a novel enzyme progressively took over the next set of compromised functions of the red alga starch synthase. The process repeated itself and the red alga enzyme retreated into a more and more specialized function until the fifth round of gene duplication covered all the functions previously assumed by the red algal starch synthase. This was swiftly followed by the disappearance of the rhodophycean enzyme. In cryptophytes, such a convoluted process of gene replacement occurred because starch metabolism in the periplast was preferred over the maintenance of the eukaryote starch metabolism in the Goniomonas-like host, which therefore vanished from the cryptistan host cytosol. This comes in stark contrast to the chromerids where the reverse happened and starch vanished from the secondary plastid of these algae leaving starch as the major sink for photosynthetic carbon in the cytosol of the host. The UDP-glucose specific SSI-II related starch synthase from cryptophytes and its dinoflagellates homologs are the sole instances where such enzymes are found in the whole tree of life. The reasons underlying their appearance during endosymbiosis are linked to the maintenance of starch synthesis in the periplast of a photosymbiont containing a red alga GT5 UDP-Glc glycogen/starch synthase of ancient eukaryotic origin. This correlates with the absence or disappearance of starch metabolism in the host cytosol. These conclusions are buttressed by the availability of whole genome sequence for *Guillardia theta* and its comparison to the now available Goniomonas genome sequences (*38*, *51*). Because the phylogeny of the dinoflagellate SSI-SSII encompasses some of the earliest diverging dinoflagellate clades such as *Oxyrrhis marina* (Fig. S16), we argue that a starch containing endosymbiont was also present at the earliest stages of dinoflagellate evolution. Subsequent complex events have since led to disappearance of this plastid genome from the current Myzozoa while the presence of a possible remnant compartment may still be suspected in some dinoflagellate species. Currently the enzymes of starch metabolism have all been rewired to cytosol of most free-living dinoflagellates (Fig. 6) The reasons and mechanism for this rewiring to the cytosol are presently unclear.

**Figure 6:**
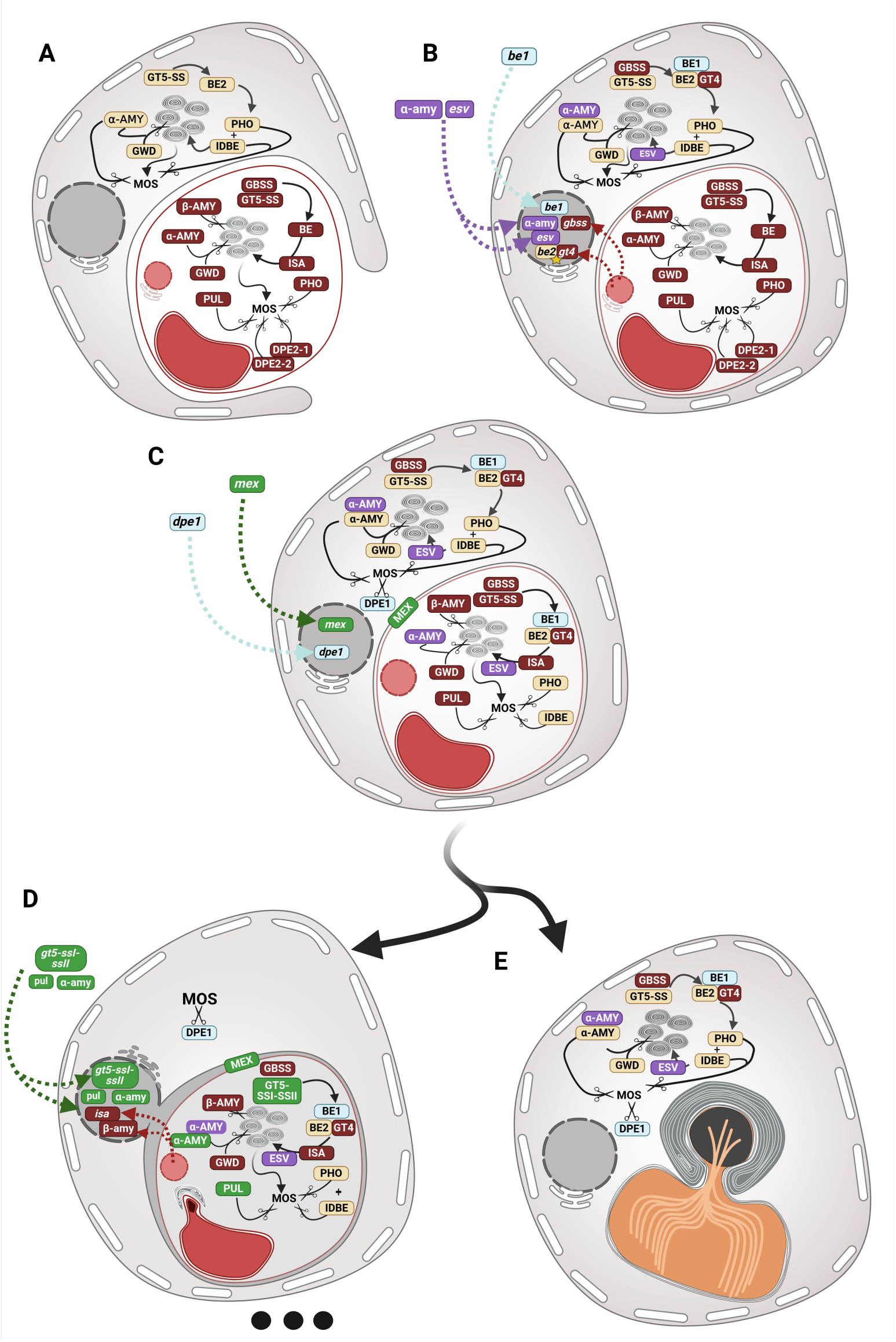
Schematic reconstruction of starch metabolism in Myzozoa. **(A)** describes the initial capture of a red alga symbiont by LCMA. Two distinct networks described in Figure 1B are operating for the synthesis of paraglycogen (amylopectin) in the cytosol of LCMA as described for early diverging ciliates (enzymes displayed in beige) and for the synthesis of starch (enzymes displayed in red) as described in the amylose containing mesophyllic red algae (enzymes displayed in red). Enzymes are represented by their abbreviations listed and defined in table 1. α-amy represents exclusively α-amylases of subfamily GH13_6. The function of each enzyme in storage polysaccharide metabolism is detailed Table 1. Arrows represent biosynthetic steps (including steps required for crystallisation and granule aggregation). Scissors icons are used to represent catabolic steps. Starch granule aggregation (grey granules) is driven by ISA in the symbiont and by PHO and iDBE in the host cytosol. **(B)** represents a series of early LGTs to the host genome by dotted arrows directed to the host nucleus and identifying their nature (table 1) and source. Procaryotic, cryptophyte and glaucophyte sources are respectively represented in light blue, purple and blue. Although we have represented these LGTs for clarity and simplicity as if they first impacted selectively the host cytosolic network, equally viable is the hypothesis that they were first selected because they rescued corresponding fitness compromised symbiont activities due to Muller’s ratchet and thereafter used for the host cytosolic network. EGT of GBSS and of the GT4 glycosyltransferase insertion into BE2 are represented by dotted arrows from the red alga nucleus (nucleomorph) to the host nucleus **(C) :** represents the integration of the early LGTs and EGTs into the symbiont’s network. Integration of enzymes from the host storage network such as PHO and iDBE is also represented (enzymes in beige). Two interdependent LGTs to the host nucleus are depicted encoding an amylomaltase of bacterial origin expressed in the host cytosol and a maltose export transporter of green alga origin localised on the symbiont membranes allowing for export of maltose produced by the symbiont’s α and β amylases following the loss of the red alga dpe2-like α-1,4 glucanotransferases. It is proposed that the cytosolic DPE1 α-1,4 glucanotransferases was acquired by LGT from cyanobacteria at this stage This achieved net export of reduced carbon to the host in the form of maltose, as do plant leaf chloroplasts today, thereby cementing photosymbiosis in LCMA. **(D)** represents the early diverging dinoflagellate lineage. By analogy with cryptophytes, we hypothesize a fusion of the outer membranes of the symbiont with the host ER. This would have favoured the export of photosynthate in the form of symbiont lipid targeting to the ER which will trigger TAG droplets synthesis on the host cytosolic face of the ER using prexisting and optimized host lipid droplet synthesis pathways. Because carbohydrate export is now a minor storage sink for carbon, the host cytosolic starch metabolism was lost. Within the symbiont, fitness erosion due to Muller’s rachets led to the selection of EGTs for the red alga isoamylase and β-amylase and for further LGTs of green algae pullulanase and α-amylase. The single red alga starch synthase was replaced by 5 distinct isoforms of GT5-SSI-SSII starch synthases of green alga origin. The three dots at the bottom symbolizes the complex and unknown evolution that followed our proposed hypothetical cryptophyte like intermediate. This possibly involved photosynthesis loss, as in cryptophytes, followed by complex membrane rearrangements, leading to the separation of the ER from the symbiont and the possible rescue of the periplast to the cytosol. The presence of multiple photosynthesis, plastome and even outright plastid losses are all evidenced in early diverging dinoflagellates, further blurring the traces of the ancient myzozoan endosymbiosis. **(E)** displays a vitrella-like chromerid ancestor whose red alga plastid displayed in panel C was simply replaced by an ochrophyte plastid (in orange). Because ochrophytes do not contain starch or glycogen metabolism, evolution of starch metabolism was arrested. Nevertheless, host cytosolic starch was retained as the major cytosolic carbon sink for photosynthesis as evidenced by the presence of a large starch sheet surrounding the ochrophyte pyrenoid (in black) across the four membranes plastid.

### Stasis in the evolution of storage polysaccharide metabolism in the absence of endosymbiosis and parasitism

Over a billion years of evolution occurred since the red and green algae and glaucophyte diverged(*54*). Despite such a long period of time, abundant genomic and functional biology evidence obtained in the green alga *C. reinhardtii*, in Arabidopsis leaves and cereal grain endosperm (maize, rice, barley and wheat) as well as in potato point to very few innovations since primary plastid endosymbiosis. A similar analogous if not identical network of 30-40 genes required for the synthesis and degradation of only 2 glycosidic linkages is indeed evidenced from the most anciently derived picophytoplanctonic green algae Ostreococcus and Micromonas to the most recently diverged C4 monocot plants (maize) (*14*, *22*). Even major events that gave birth to or affected many important plant pathways such as plant terrestrialisation does not appear to have affected the essential features and basic composition of the highly complex Chloroplastida starch metabolism. Mutation inactivating given components of this network display similar or identical phenotypes when looking at a plant leaf cell experiencing recurrent starch synthesis and degradation or an alga culture experiencing clock-controlled cycles of polysaccharide synthesis and degradation. More variation in mutant phenotype expressivity will be found between leaf and grain endosperm starch or between potato tubers and leaves than between Chlamydomonas growing cultures and potato leaves(*70*). In addition, Chlamydomonas can to some extent reproduce the variations between source and sink tissues of plants by simply arresting growth through nutrient starvation (*71*). Such an astonishing degree of conservation of metabolic network composition and function deserves to be referred to as “stasis”. Evidence for stasis in cytosolic storage polysaccharide metabolism is also shown by comparative genomics in red algae (*14*, *72*, *73*). This pathway shows evidence of pathway streamlining when compared to the glaucophytes(*14*, *74*, *75*). This streamlining was selected to maintain the number of genes required for starch metabolism as low as possible, in agreement with the general trend of nuclear genome compactness and simplicity that characterizes all Rhodophyceae(*76*). In Rhodophyceae the changes are restricted to the loss of GBSS and consequently of amylose synthesis that seemed to have happened twice: once in the common ancestor of cyanidiales and once in the ancestor of multicellular red algae. Of interest is the loss of GWD/PWD in the Cyanidiales *G. sulphuraria* which correlates, in this species only, among all red algae with the accumulation of glycogen and loss of GWD/PWD and pullulanase(*16*). Overall, while mesophyllic unicellular red algae seem slightly enriched in catabolic enzymes the level of stasis seen here is also very significant. Unfortunately, we do not have access to a wealth of functional studies by comparison to plants. Functional studies are indeed largely limited to the study of mutants of *C. merolae* defective in isoamylase and a few biochemical studies (*29*, *77*). As to the glaucophytes, the pathway seems very similar to the standard β-amylase containing version of glycogen metabolism seen in Entamoeba, Dictyostellium or Trichomonas (*78*). Of significance is the presence in all glaucophytes of indirect debranching enzyme which is a relic of eukaryotic glycogen metabolism (Figs. 1B, 2 and S2). Chlamydial TTS effector enzymes such as prokaryotic glycogen/starch synthase and isoamylase are also evidenced(*23*, *74*). Although a significant level of gene duplications reflecting possible interesting cases of enzyme subfunctionalisations have apparently occurred in Cyanophora we have no way of making decisions on how significant and stable they are, in the absence of complete genome sequences from other glaucophyte species. Only one biochemical study dealing with glaucophyte starch metabolism has appeared (*79*). We can only speculate that “stasis” will also be evidenced in the future for the glaucophyte lineage but presently the evidence is unfortunately lacking.

Finally, by looking at comparative genomics of ciliate starch metabolism we see very little evidence for radical changes, save the occasional gain and losses of hydrolytic enzymes involved in prey digestion or, more telling, the loss of the glucan water dikinases and ESV/LESV which correlate with the replacement of amylopectin by standard glycogen accumulation as witnessed in the late diverging ciliates Paramecium and Tetrahymena (Fig. 1A) thereby recapitulating the evolution seen in *G. sulphuraria* among the red algae. By comparison to the stasis seen in extant Archaeplastida and ciliates, the evolution that, following primary plastid endosymbiosis, led from an ancestral glycogen or amylopectin accumulating eukaryotic phagotroph to the starch metabolism witnessed in the 3 Archaeplastida lineages is summarized in Table S1. Although the stepwise complex evolution depicted in Table S1 does not lend itself to accurate quantifications the number of steps involved in the evolution of red algae and glaucophyte metabolism is globally similar to the evolution reported here for the chromerid lineage albeit assuming the likely prior presence of a ciliate-like pathway in the chromerid ancestor (Fig. 3). Strikingly the targeting to the primary plastid of the starch pathway that characterizes storage polysaccharide evolution in the green algae further entails many additional steps building on the previous platform of cytosolic starch metabolism witnessed in glaucophytes and red algae(*21*, *80*). As outlined above, five major acquisitions of enzyme activities typify the evolution of all Myzozoa while this process virtually halted thereafter in the chromerid/apicomplexan lineage as the dinoflagellate diverged (Fig. 3 and S7 to S12 and S15). The presence of a starch containing symbiont is here hypothesized as the sole plausible explanation for the extensive evolution of starch synthases seen at the earliest stages of dinoflagellate divergence (Fig. 6D). In such a context, we believe that the same symbiont triggered the evolution seen in the chromerid/apicomplexan lineages before dinoflagellate divergence. In our view this would account for the extensive evolution common to all Myzozoa.

Nevertheless, when taking a closer look to what happened selectively in the chromerid/apicomplexan lineage, arrest of enzyme acquisition was not followed by the type of stasis exemplified by the green algae and plants or in the red algae. This is due to extensive cases of parasitism evolving in the myzozoan lineages including both the Apicomplexa and the early diverging dinoflagellates. *C. velia* already contains significantly less enzyme isoforms involved in storage polysaccharide metabolism than *V. brassicaformis*. These reductions in enzyme isoform numbers include a single form of UDP-glucose GT5 glycogen/starch synthase and a single BE1 type of branching enzyme compared to 2 and 4 isoforms respectively in *V. brassicaformis* (Fig. 1A). The loss by *C. velia* of both GBSS and DPE1 comes in addition to a significant decrease in the number of amylases and dikinase isoforms by comparison to *V. brassicaformis*. Similar and more dramatic losses are evidenced in the apicomplexa. The total count of candidate CAZymes that potentially affect starch metabolism within the chromerid/apicomplexan lineage ranks *V. brassicaformis* as N°1 with 36 candidate CAZymes followed by *C. velia* (19), Toxoplasma gondii (12) and finally *C. parvum* (7) (Fig. 1A). *C. parvum* holds presently the lowest CAZyme count record among all starch accumulating eukaryotes. Interestingly, we were not able to find any sequence related to the classical UDPG GT5 glycogen synthase of ancient eukaryotic origin in *C. parvum*. One GBSS-like enzyme was however found in other cryptosporidium strains bearing sufficient resemblance with GBSS to cluster with other myzozoan GBSS sequences during phylogenetic analysis albeit with long branches, attesting the presence of a fast-evolving enzyme (Fig. 3 and S12). It must be further noted that some of the most dangerous apicomplexa parasites to humans such as the malaria parasite Plasmodium have lost storage polysaccharide metabolism outright thereby closing the cycle of parasitic reduction of storage polysaccharide metabolism characterising the chromerid/apicomplexan lineage.

### Evidence for both an ancient endosymbiont pathway of starch metabolism and of an ancient cytosolic alveolate pathway in the last myzozoan common ancestor

Although a number of morphological and ultrastructural cell features unite dinoflagellates chromerids, apicomplexans, and their sister lineages, molecular synapomorphies supporting a hypothetical common origin of their plastids uniting all Myzozoa remain scarce and are limited to the presence of an ancient LGT of a gene encoding Rubisco II and of the presence of a potentially common plastidial transcript polyuridinylation machinery (although the genes involved still need to be described) (*6*, *7*).

However, convincing phylogenetic and biochemical evidence point to the presence of distinct ancestral plastids in both the dinoflagellate lineage where haptophytes were initially suspected to have donated the peridinin plastids, and the apicomplexans and chromerids where it was hypothesized that ochrophytes gave rise to the current plastid (*10*, *81*).

Nevertheless, we do not know if these plastids indeed correspond to the first common endosymbiotic event characterized by the two unique synapomorphies described above. Equally viable is a scenario whereby a founding endosymbiosis involving a distinct lineage initiated the complex endosymbiotic history of Myzozoa started (*82*, *83*). Thereafter if the genes encoding the Rubisco II and polyuridinylation machinery were conserved, these nuclear host genes could have been recruited by novel incipient plastids. Our data suggest that starch metabolism pathways were initially present in LCMA in both the alveolate host cytosol as in chromerids, ciliates and Apicomplexa and in the periplastidial space of a complex plastid as in cryptophytes. Thereafter cytosolic starch metabolism was lost in the rising dinoflagellates while the original symbiont was lost in the rising chromerid/apicomplexa and subsequently replaced by an ochrophyte plastid (Fig. 6). Recent evidence gathered from studies performed on Kareniaceae dinoflagellates demonstrate that the outright replacement of a preexisting plastid by plastids from another source while not having been detected in other photosynthetic eukaryotes, is possibly routine in Myzozoa despite the requirement for LGT of many genes required to stabilize the newly acquired organelle (*45*). The dual localisation of starch metabolism in LCMA would have resulted from either a secondary or tertiary endosymbiosis involving a paraglycogen synthesizing alveolate host with respectively a red alga or a cryptophyte endosymbiont, respectively. The rationale for this hypothesis is as follows:

1. Acquisitions of five enzyme (synapomorphies) invovled in starch metabolism is shared by most Myzozoa further suggesting that these acquisitions impacted both the symbiont and the cytosolic enzyme networks of storage polysaccharide metabolism (Fig. 6A). As this amount of evolution was not witnessed after conclusion of primary endosymbiosis either in green or red algae nor is it seen in free living glycogen accumulating protists, we infer that these significant changes were propelled by an ongoing secondary or tertiary endosymbiosis involving a starch containing symbiont common to all Myzozoa. Indeed, they may have resulted from the ongoing process of EGTs and LGTs selection required to transfer compromised symbiont functions to the nuclear host genome. These genes may thus have impacted both the symbiont and host enzyme networks within the short time where both co-existed in the same cells (Fig. 6B)
2. Metabolic integration of the starch containing complex (secondary or higher order) endosymbiont continued selectively in the rising dinoflagellates until all required genes likely to be compromised by Mueller’s ratchet could be rescued by EGTs, LGTs or rescuing genes encoding enzymes from the host starch metabolism network. This led to seven further major additions to the pathway that could only have been completed if starch metabolism had remained functional inside the symbiont throughout the whole process. However, there was no requirement for such maintenance with respect to cytosolic starch metabolism. (Fig. 6C)
3. The dinoflagellates have reproduced very early on after divergence the evolution witnessed in cryptophytes. This evolution consisted in the selection of several distinct isoforms of an entirely different and unique form of starch synthase that evolved to replace the red alga symbiont starch synthase (see discussion above). We argue that this was accompanied, as in cryptophytes, by the loss of starch metabolism in the ancestral host cytosol (Fig. 6D)
4. The rising chromerid/apicomplexa lineages after acquiring the five aforementioned enzymes witness an arrest of such acquisitions. This arrest would have been prompted by either the loss of the symbiont or the replacement of the starch-accumulating plastid by an ochrophyte plastid. Ochrophytes are known to be devoid of starch metabolism both in the plastid or the host cytosol thereby arresting further additions. Starch would thus have remained in the alveolate cytosol where it would have defined the predominant sink for photosynthetic carbon export (Fig. 6E).

### Impact of EGT versus host gene recruitment or LGT on the mechanism of starch granule aggregation in cryptophytes and dinoflagellates

As discussed above both extrinsic and intrinsic factors may impede EGT of important functions for starch metabolism. This may have led to the selection of unexpected alternative solutions resulting in the greatest diversity of genes present in the environment. The selective presence of nucleomorphs in cryptophytes not only bears witness to the red alga identity of the symbiont but it also suggests that the mechanism of reductive genome evolution of the nucleomorph may have been on pause quite early, further complicating recruitment of genes by standard EGT. If the transfer window hypothesis is correct ‘(see the above discussion) this could explain why the cryptophytes did not manage to recruit their own red alga isoamylase gene through EGT. The transfer window may have indeed closed before they had a chancechance or needed to do so. The fact that the Goniomonas-like ancestor was likely an amylopectin producer suggests the presence in the host genome of genes encoding suitable enzymes to fulfil the function supplied for starch granule aggregation by isoamylase in red algae. The closing of the transfer window indeed does not preclude recruitment of host genes or LGTs. We have produced functional evidence from mutant studies in *C. velia* supporting that both phosphorylase and iDBE are required to aggregate amylopectin in ciliates and chromerids/apicomplexan lineage (Chabi et al. submitted). By analogy with the ciliates, we infer that the heterotrophic ancestor of cryptophytes may be using phosphorylase to the same end. It is thus interesting that in *Guillardia theta* a host phosphorylase gene was recruited in addition to that of another presumably green alga inherited isoform. Equally intriguing is the presence of the ESV/LESV protein, defining an LGT of a gene encoding a non-catalytic protein of putative green alga origin (Fig. 1B and S6). This “polysaccharide chaperone” protein may be assisting starch aggregation by the host phosphorylase in the foreign symbiont environment, as supported by its activity in a semi-vitro yeast system for starch granule aggregation. We were unable to find the gene in heterotrophic cryptistan ancestors suggesting it may have been selected to assist a host enzyme in the periplast after endosymbiosis within cryptophytes (Fig. 6B). Although phylogenetic analysis of single enzyme offers, often, insufficient resolution to make definitive statements, the topology of the ESV-LESV tree displayed in Fig. S6 may nevertheless suggest that the ESV-LESV gene may have been selected twice independently from either a green algal source in cryptophytes and from glaucophytes in alveolates.

The process of EGT/host gene recruitment/LGT that characterizes evolution of pathways impacted by endosymbiosis driven metabolic integration offers a straightforward explanation for the recruitment of a red algal isoamylase in the dinoflagellate lineage. If, as proposed in this work, a starch-containing symbiont was being integrated in the rising dinoflagellates with concomitant disappearance of cytosolic starch metabolism in the host, then the genes possibly encoding functions controlling starch granule aggregation in the host would if maintained possibly have acquired plastid targeting sequences and expressed their products in the symbiont. This process however occurs one gene at a time and the selection value of each step must be considered. If, for instance, the host phosphorylase is transferred first to the symbiont, it will be selected for its major and effective individual contribution to polysaccharide degradation in replacement of the ratchet compromised phosphorylase of the symbiont. Despite phosphorylase and iDBE being the two major determinants of starch crystallisation (Chabi et al. submitted), we argue that the individual transfer of phosphorylase had no selection value in this process in the absence of its iDBE partner. Likewise, the transfer the host iDBE to the symbiont in the absence of its phosphorylase partner would have had no selective value with respect to polysaccharide crystallisation. However, iDBE on its own does have selection value as an enzyme supplying a well-established α-1,4 glucanotransferase domain likely to be effective for degrading chains produced by isoamylase. Alveolata indeed characteristically lack α-1,4-glucanotransferases as they do not contain isoamylases, pullulanases or β-amylases which produce subtrates for these enzymes save the dinoflagellates. The iDBE surrogate of such an enzyme was thus highly desirable in that lineage. Hence within the symbiont neither phosphorylase nor iDBE were selected for their complex interplay demonstrated to ensure starch crystallisation in chromerids/apicomplexan lineage and ciliates. A red algal isoamylase was thus required and recruited either as an EGT (if the symbiont was a red alga) or an LGT (if the symbiont was a cryptophyte).

### Conclusion: towards a model explaining the major features of starch metabolism evolution in Myzozoa

In our phylogenetic analysis, four candidate “green genes” have been selected in the starch metabolism pathway of dinoflagellates (Fig. 6C and D). These consist of MEX1 (Figs. 4 and S20), of a GT5 starch synthase derived from the green alga SSI/SSII type of enzymes (Figs. 4 and S16), a pullulanase (a direct debranching enzyme involved in both starch catabolism and isoamylase function) (Fig. 4 and S18) and one of the two GH13_6 α-amylase isoforms (Fig. 3 and S10) related to the plastidial enzymes of green algae. Strikingly no green gene LGTs are found in the chromerid and apicomplexa starch metabolism networks. As explained above, green genes would have been at a selective advantage if the proteins needed to follow the organelle targeting machinery which was obviously not the case in the rising chromerids and apicomplexa if we assume that the starch pathway remained in the host cytosol in these organisms. This further strengthens our suggestion that when the dinoflagellates evolved, they lost alveolate cytosolic starch synthesis and kept starch synthesis in the cytosol of the symbiont (now called the periplast) in a fashion similar to cryptophytes, while the reverse happened in the chromerid lineage. In addition to green genes the dinoflagellates have acquired β-amylase (Fig. 4D, S19) genes of red algal origin as well as two additional red alga genes encoding isoamylase (Fig. 4 and S17) and GBSS (Fig. 3E and S12). Interestingly such enzymes have all been demonstrated to be readily targeted to the evolving chloroplast when the starch pathway was relocated from the cytosol to the chloroplast selectively in the green lineage after primary plastid endosymbiosis (*21*, *29*, *53*). Hence substitution of red genes with green versions of the gene is selectively observed for those functions that already have a past record of failing to be transferred to the chloroplast during evolution of the green algae.

To summarize, comparison of the starch metabolism networks of dinoflagellates with those of the chromerid/apicomplexan lineage suggest that both lineages emerged from a common complex (secondary or higher) endosymbiosis. This is supported by acquisition of a shared EGT of a red alga or cryptophyte GBSS gene and a common BE1 acquisition from bacterial LGT that preceded the common disruption of the alveolate BE2 by a GT4 gene of common symbiogenic origin (either of red alga or cryptophyte origin) (Fig. 6B, 3B and S8 and S9). We suspect that acquisition of the myzozoan GH13_6 hydrolase was selected at that time through LGT from either prokaryotes or eukaryotes to provide suitable nuclear encoded replacements of mutation accumulating periplastidial enzymes of the symbiont (Fig. 3C, 6B and S10). We highly suspect that the initial prokaryotic LGT of BE1 was selected similarly to rescue the BE symbiont enzyme. It then triggered the evolution of BE2 in both the cytosol and the symbiont. All this happened in a context where LCMA acquired photosynthesis as suggested by the common presence of the CBM20-containing SGNH hydrolase superfamily carbohydrate modifying enzyme (Fig. 3F and S15)which is selectively present in photosynthetic organisms and absent or lost from heterotrophic derivatives. At this very early stage both the symbiont and its alveolate host synthesized starch. However, this did not last very long (starch synthesized in distinct cellular compartments has never been reported) (Fig. 6B and C) and natural selection favoured the loss of starch from the symbiont in the chromerid/apicomplexa (Fig. 6E) while the reverse happened in the rising dinoflagellates (Fig. 6D). This signalled an arrest of further endosymbiont driven evolution of starch metabolism in the chromerid/apicomplexan lineage which kept major functional features of the ancient alveolate pathway exemplified by ciliates. On the contrary in the evolving dinoflagellate lineage the process of EGT and LGT associated with plastid genome evolution continued, leading to the acquisition of the green enzymes such as starch synthases, mex, α-amylase and pullulanase and EGTs of red alga isoamylases and β-amylases (Fig. 6D) of either red alga or cryptophyte origin. This led to an entirely different pathway with possibly distinct mechanisms of amylopectin crystallisation relying more classically on isoamylase. Although we have no reason to favour red algae over cryptophytes as the likely initial common symbiont that triggered the diversification of Myzozoa, we have applied the knowledge generated on secondary red alga endosymbiosis by the accessibility of both *G. theta* and *G. avonlea* genome sequences to propose a well-supported possible sequence of events accounting for the early evolution of starch metabolism in Myzozoa. Despite this model, there are, as yet, gaping holes in our understanding of the complex secondary or tertiary plastid evolution in dinoflagellates to fully grasp why the hypothesized ancient myzozoan plastid was eventually lost and how the periplastidial starch metabolism pathway was rescued in the dinoflagellate cytosol. For the sake of clarity in Fig. 6 we have recapitulated the evolution of starch metabolism in Myzozoa. In this Fig. we assume that LCMA has undergone a secondary endosymbiosis with a red alga. Equally viable is, in our view, the alternative hypothesis, i.e., that the founding myzozoan endosymbiosis was a tertiary endosymbiosis involving a cryptophyte. Our results cannot easily accommodate an initial myzozoan endosymbiosis which involved neither an Archaeplastida nor a cryptophyte symbiont. The amount of evolution seen in Myzozoa indeed dwarves that seen in starch accumulating Archaeplastida post primary endosymbiosis or those seen in glycogen-accumulating protists in the absence of endosymbiosis-driven evolution. Strikingly, however, the arrest of starch metabolism evolution in chromerid and apicomplexan plastids calls for the replacement of the original myzozoan plastid by a starchless organelle prior to its completed metabolic integration such as those characterizing, among others, the ochrophytes and the haptophytes. Our results however do not presently discriminate competing hypotheses concerning the origin of the peridinin plastid of dinoflagellates.

## Resource availability

### Lead contact

Further information and requests for resources and reagents should be directed to and will be fulfilled by the lead contact, Steven Ball.

### Data and code availability

All raw sequences, alignments, and trees reported in this paper are available from the lead contact upon request.

## Acknowledgments

This work was supported by the French Agence National de la Recherche (CONVERGE project, grant agreement ANR-21-CE44-0017), the Région Hauts de France: ALIBIOTECH (FEDER), the Centre National de la Recherche Scientifique (CNRS), CHROSME (AO-DBM), the I-site Université de Lille, CDP-PIE.

## Author contributions

S.G.B. conceived the study. U.C., M.C., M.F., P.D., M.L. and C.T. performed the experiments. U.C. and S.G.B. supervised the study. S.G.B. wrote the original draft. M.O., J.M.A. and C.C. reviewed and edited the manuscript.

## References

1. S. M. Adl, D. Bass, C. E. Lane, J. Lukeš, C. L. Schoch, A. Smirnov, S. Agatha, C. Berney, M. W. Brown, F. Burki, P. Cárdenas, I. Čepička, L. Chistyakova, J. Del Campo, M. Dunthorn, B. Edvardsen, Y. Eglit, L. Guillou, V. Hampl, A. A. Heiss, M. Hoppenrath, T. Y. James, A. Karnkowska, S. Karpov, E. Kim, M. Kolisko, A. Kudryavtsev, D. J. G. Lahr, E. Lara, L. Le Gall, D. H. Lynn, D. G. Mann, R. Massana, E. A. D. Mitchell, C. Morrow, J. S. Park, J. W. Pawlowski, M. J. Powell, D. J. Richter, S. Rueckert, L. Shadwick, S. Shimano, F. W. Spiegel, G. Torruella, N. Youssef, V. Zlatogursky, Q. Zhang, Revisions to the Classification, Nomenclature, and Diversity of Eukaryotes. J Eukaryotic Microbiology 66, 4–119 (2019).

2. T. Cavalier-Smith, E. E. Chao, Protalveolate phylogeny and systematics and the origins of Sporozoa and dinoflagellates (phylum Myzozoa nom. nov.). European Journal of Protistology 40, 185–212 (2004).

3. A. Reyes-Prieto, A. Moustafa, D. Bhattacharya, Multiple Genes of Apparent Algal Origin Suggest Ciliates May Once Have Been Photosynthetic. Current Biology 18, 956–962 (2008).

4. J. M. Archibald, “The Evolution of Algae by Secondary and Tertiary Endosymbiosis” in Advances in Botanical Research (Elsevier, 2012; https://linkinghub.elsevier.com/retrieve/pii/B9780123914996000037)vol. 64, pp. 87–118.

5. P. J. Keeling, The Number, Speed, and Impact of Plastid Endosymbioses in Eukaryotic Evolution. Annual Review of Plant Biology 64, 583–607 (2013).

6. D. Morse, P. Salois, P. Markovic, J. W. Hastings, A nuclear-encoded form II RuBisCO in dinoflagellates. Science 268, 1622–1624 (1995).

7. J. Janouškovec, A. Horák, M. Oborník, J. Lukeš, P. J. Keeling, A common red algal origin of the apicomplexan, dinoflagellate, and heterokont plastids. Proc. Natl. Acad. Sci. U.S.A. 107, 10949–10954 (2010).

8. H. S. Yoon, J. D. Hackett, D. Bhattacharya, A single origin of the peridinin- and fucoxanthin-containing plastids in dinoflagellates through tertiary endosymbiosis. Proc Natl Acad Sci U S A 99, 11724–11729 (2002).

9. J. Petersen, A.-K. Ludewig, V. Michael, B. Bunk, M. Jarek, D. Baurain, H. Brinkmann, Chromera velia, Endosymbioses and the Rhodoplex Hypothesis—Plastid Evolution in Cryptophytes, Alveolates, Stramenopiles, and Haptophytes (CASH Lineages). Genome Biology and Evolution 6, 666–684 (2014).

10. T. Ševčíková, A. Horák, V. Klimeš, V. Zbránková, E. Demir-Hilton, S. Sudek, J. Jenkins, J. Schmutz, P. Přibyl, J. Fousek, Č. Vlček, B. F. Lang, M. Oborník, A. Z. Worden, M. Eliáš, Updating algal evolutionary relationships through plastid genome sequencing: did alveolate plastids emerge through endosymbiosis of an ochrophyte? Sci Rep 5, 10134 (2015).

11. T. G. Stephens, M. A. Ragan, D. Bhattacharya, C. X. Chan, Core genes in diverse dinoflagellate lineages include a wealth of conserved dark genes with unknown functions. Sci Rep 8, 17175 (2018).

12. A. M. G. Novák Vanclová, R. G. Dorrell, “Complex Plastids Across the Eukaryotes: An Overview of Inherited and Convergently Evolved Characters” in Endosymbiotic Organelle Acquisition, S. D. Schwartzbach, P. G. Kroth, M. Oborník, Eds. (Springer International Publishing, Cham, 2024; https://link.springer.com/10.1007/978-3-031-57446-7_2), pp. 39–88.

13. M. Oborník, Endosymbiotic Evolution of Algae, Secondary Heterotrophy and Parasitism. Biomolecules 9, 266 (2019).

14. S. Ball, C. Colleoni, U. Cenci, J. N. Raj, C. Tirtiaux, The evolution of glycogen and starch metabolism in eukaryotes gives molecular clues to understand the establishment of plastid endosymbiosis. Journal of Experimental Botany 62, 1775–1801 (2011).

15. A. L. Reysenbach, N. Ravenscroft, S. Long, D. T. Jones, D. R. Woods, Characterization, Biosynthesis, and Regulation of Granulose in *Clostridium acetobutylicum*. Appl Environ Microbiol 52, 185–190 (1986).

16. S. Ball, C. Colleoni, M. C. Arias, “The Transition from Glycogen to Starch Metabolism in Cyanobacteria and Eukaryotes” in Starch, Y. Nakamura, Ed. (Springer Japan, Tokyo, 2015; http://link.springer.com/10.1007/978-4-431-55495-0_4), pp. 93–158.

17. M. Colpaert, M. Chabi, U. Cenci, C. Colleoni, “Storage Polysaccharides in Prokaryotes: Glycogen, Granulose, and Starch-Like Granules” in Bacterial Organelles and Organelle-like Inclusions, D. Jendrossek, Ed. (Springer International Publishing, Cham, 2020; http://link.springer.com/10.1007/978-3-030-60173-7_8)vol. 34 of *Microbiology Monographs*, pp. 177–210.

18. G. Ritte, J. R. Lloyd, N. Eckermann, A. Rottmann, J. Kossmann, M. Steup, The starch-related R1 protein is an α-glucan, water dikinase. Proc. Natl. Acad. Sci. U.S.A. 99, 7166–7171 (2002).

19. M. Stitt, S. C. Zeeman, Starch turnover: pathways, regulation and role in growth. Current Opinion in Plant Biology 15, 282–292 (2012).

20. U. Cenci, F. Nitschke, M. Steup, B. A. Minassian, C. Colleoni, S. G. Ball, Transition from glycogen to starch metabolism in Archaeplastida. Trends in Plant Science 19, 18–28 (2014).

21. P. Deschamps, I. Haferkamp, C. d’Hulst, H. E. Neuhaus, S. G. Ball, The relocation of starch metabolism to chloroplasts: when, why and how. Trends Plant Sci 13, 574–582 (2008).

22. P. Deschamps, C. Colleoni, Y. Nakamura, E. Suzuki, J.-L. Putaux, A. Buléon, S. Haebel, G. Ritte, M. Steup, L. I. Falcón, others, Metabolic symbiosis and the birth of the plant kingdom. Molecular biology and evolution 25, 536–548 (2008).

23. S. G. Ball, A. Subtil, D. Bhattacharya, A. Moustafa, A. P. M. Weber, L. Gehre, C. Colleoni, M.-C. Arias, U. Cenci, D. Dauvillée, Metabolic Effectors Secreted by Bacterial Pathogens: Essential Facilitators of Plastid Endosymbiosis? The Plant Cell 25, 7–21 (2013).

24. S. Ball, H.-P. Guan, M. James, A. Myers, P. Keeling, G. Mouille, A. Buléon, P. Colonna, J. Preiss, From glycogen to amylopectin: a model for the biogenesis of the plant starch granule. Cell 86, 349–352 (1996).

25. G. Mouille, M.-L. Maddelein, N. Libessart, P. Talaga, A. Decq, B. Delrue, S. Ball, Preamylopectin processing: a mandatory step for starch biosynthesis in plants. The Plant Cell 8, 1353–1366 (1996).

26. D. Dauvillée, C. Colleoni, G. Mouille, A. Buléon, D. J. Gallant, B. Bouchet, M. K. Morell, C. d’Hulst, A. M. Myers, S. G. Ball, Two Loci Control Phytoglycogen Production in the Monocellular Green Alga *Chlamydomonas reinhardtii*. Plant Physiology 125, 1710–1722 (2001).

27. F. Wattebled, Y. Dong, S. Dumez, D. Delvallé, V. Planchot, P. Berbezy, D. Vyas, P. Colonna, M. Chatterjee, S. Ball, C. D’Hulst, Mutants of Arabidopsis Lacking a Chloroplastic Isoamylase Accumulate Phytoglycogen and an Abnormal Form of Amylopectin. Plant Physiology 138, 184–195 (2005).

28. S. Streb, T. Delatte, M. Umhang, S. Eicke, M. Schorderet, D. Reinhardt, S. C. Zeeman, Starch granule biosynthesis in Arabidopsis is abolished by removal of all debranching enzymes but restored by the subsequent removal of an endoamylase. Plant Cell 20, 3448–3466 (2008).

29. T. Maeno, Y. Yamakawa, Y. Takiyasu, H. Miyauchi, Y. Nakamura, M. Ono, N. Ozaki, Y. Utsumi, U. Cenci, C. Colleoni, S. Ball, M. Tsuzuki, S. Fujiwara, One of the isoamylase isoforms, CMI294C, is required for semi-amylopectin synthesis in the rhodophyte Cyanidioschyzon merolae. Front. Plant Sci. 13, 967165 (2022).

30. U. Cenci, M. Chabi, M. Ducatez, C. Tirtiaux, J. Nirmal-Raj, Y. Utsumi, D. Kobayashi, S. Sasaki, E. Suzuki, Y. Nakamura, J.-L. Putaux, X. Roussel, A. Durand-Terrasson, D. Bhattacharya, A.-S. Vercoutter-Edouart, E. Maes, M. C. Arias, M. Palcic, L. Sim, S. G. Ball, C. Colleoni, Convergent Evolution of Polysaccharide Debranching Defines a Common Mechanism for Starch Accumulation in Cyanobacteria and Plants. The Plant Cell 25, 3961–3975 (2013).

31. T. A. Hennen-Bierwagen, M. G. James, C. J. Newton, E. M. Juhl, U. Cenci, S. Ball, C. Colleoni, S. L. Shuler, W. F. Tracy, A. T. Culbertson, A. M. Myers, Noncatalytic functions of ISOAMYLASE 1 and 2 affect the proportion of insoluble and soluble α-polyglucans in maize. The Plant Cell 37, koaf220 (2025).

32. B. Pfister, A. Sánchez-Ferrer, A. Diaz, K. Lu, C. Otto, M. Holler, F. R. Shaik, F. Meier, R. Mezzenga, S. C. Zeeman, Recreating the synthesis of starch granules in yeast. eLife 5, e15552 (2016).

33. D. J. Manners, J. F. Ryley, Studies on the metabolism of the Protozoa. 2. The glycogen of the ciliate *Tetrahymena pyriformis* ( *Glaucoma piriformis* ). Biochemical Journal 52, 480–482 (1952).

34. F. Verni, G. Rosati, Preliminary survey of the morphology and cytochemistry of polysaccharides reserve in ciliates. (1980).

35. Y. Guérardel, D. Leleu, A. Coppin, L. Liénard, C. Slomianny, G. Strecker, S. Ball, S. Tomavo, Amylopectin biogenesis and characterization in the protozoan parasite Toxoplasma gondii, the intracellular development of which is restricted in the HepG2 cell line. Microbes and Infection 7, 41–48 (2005).

36. R. G. F. Visser, I. Somhorst, G. J. Kuipers, N. J. Ruys, W. J. Feenstra, E. Jacobsen, Inhibition of the expression of the gene for granule-bound starch synthase in potato by antisense constructs. Molec. Gen. Genet. 225, 289–296 (1991).

37. M. van de Wal, C. D’Hulst, J. P. Vincken, A. Buléon, R. Visser, S. Ball, Amylose is synthesized in vitro by extension of and cleavage from amylopectin. J Biol Chem 273, 22232–22240 (1998).

38. U. Cenci, S. J. Sibbald, B. A. Curtis, R. Kamikawa, L. Eme, D. Moog, B. Henrissat, E. Maréchal, M. Chabi, C. Djemiel, A. J. Roger, E. Kim, J. M. Archibald, Nuclear genome sequence of the plastid-lacking cryptomonad Goniomonas avonlea provides insights into the evolution of secondary plastids. BMC Biol 16, 137 (2018).

39. D. Feike, D. Seung, A. Graf, S. Bischof, T. Ellick, M. Coiro, S. Soyk, S. Eicke, T. Mettler-Altmann, K. J. Lu, M. Trick, S. C. Zeeman, A. M. Smith, The Starch Granule-Associated Protein EARLY STARVATION1 Is Required for the Control of Starch Degradation in *Arabidopsis thaliana* Leaves. Plant Cell 28, 1472–1489 (2016).

40. R. Osman, M. Bossu, D. Dauvillée, C. Spriet, C. Liu, S. C. Zeeman, C. D’Hulst, C. Bompard, LIKE EARLY STARVATION 1 interacts with amylopectin during starch biosynthesis. Plant Physiology 195, 1851–1865 (2024).

41. C. Liu, B. Pfister, R. Osman, M. Ritter, A. Heutinck, M. Sharma, S. Eicke, M. Fischer-Stettler, D. Seung, C. Bompard, M. R. Abt, S. C. Zeeman, LIKE EARLY STARVATION 1 and EARLY STARVATION 1 promote and stabilize amylopectin phase transition in starch biosynthesis. Sci. Adv. 9, eadg7448 (2023).

42. I. Malinova, H. Mahto, F. Brandt, S. AL-Rawi, H. Qasim, H. Brust, M. Hejazi, J. Fettke, EARLY STARVATION 1 specifically affects the phosphorylation action of starch-related dikinases. The Plant Journal 95, 126–137 (2018).

43. A. Singh, J. Compart, S. A. AL-Rawi, H. Mahto, A. M. Ahmad, J. Fettke, LIKE EARLY STARVATION 1 alters the glucan structures at the starch granule surface and thereby influences the action of both starch-synthesizing and starch-degrading enzymes. The Plant Journal 111, 819–835 (2022).

44. J. Yang, Z. He, C. Chen, J. Zhao, R. Fang, Starch Branching Enzyme 1 Is Important for Amylopectin Synthesis and Cyst Reactivation in Toxoplasma gondii. Microbiol Spectr 10, e0189121 (2022).

45. A. M. Novák Vanclová, C. Nef, Z. Füssy, A. Vancl, F. Liu, C. Bowler, R. G. Dorrell, New plastids, old proteins: repeated endosymbiotic acquisitions in kareniacean dinoflagellates. EMBO Rep 25, 1859–1885 (2024).

46. J. R. Harris, M. Adrian, F. Petry, Amylopectin: a major component of the residual body in *Cryptosporidium parvum* oocysts. Parasitology 128, 269–282 (2004).

47. M. Oborník, D. Modrý, M. Lukeš, E. Černotíková-Stříbrná, J. Cihlář, M. Tesařová, E. Kotabová, M. Vancová, O. Prášil, J. Lukeš, Morphology, Ultrastructure and Life Cycle of Vitrella brassicaformis n. sp., n. gen., a Novel Chromerid from the Great Barrier Reef. Protist 163, 306–323 (2012).

48. L. M. Patrone, S. T. Broadwater, J. L. Scott, ULTRASTRUCTURE OF VEGETATIVE AND DIVIDING CELLS OF THE UNICELLULAR RED ALGAE *RHODELLA VIOLACEA* AND *RHODELLA MACULATA* ^1^. Journal of Phycology 27, 742–753 (1991).

49. J. Scott, E.-C. Yang, J. A. West, A. Yokoyama, H.-J. Kim, S. L. De Goer, C. J. O’Kelly, E. Orlova, S.-Y. Kim, J.-K. Park, H.-S. Yoon, On the genus Rhodella, the emended orders Dixoniellales and Rhodellales with a new order Glaucosphaerales (Rhodellophyceae, Rhodophyta). ALGAE 26, 277–288 (2011).

50. A. Izumo, S. Fujiwara, T. Sakurai, S. G. Ball, Y. Ishii, H. Ono, M. Yoshida, N. Fujita, Y. Nakamura, A. Buléon, M. Tsuzuki, Effects of granule-bound starch synthase I-defective mutation on the morphology and structure of pyrenoidal starch in Chlamydomonas. Plant Science 180, 238–245 (2011).

51. B. A. Curtis, G. Tanifuji, F. Burki, A. Gruber, M. Irimia, S. Maruyama, M. C. Arias, S. G. Ball, G. H. Gile, Y. Hirakawa, J. F. Hopkins, A. Kuo, S. A. Rensing, J. Schmutz, A. Symeonidi, M. Elias, R. J. M. Eveleigh, E. K. Herman, M. J. Klute, T. Nakayama, M. Oborník, A. Reyes-Prieto, E. V. Armbrust, S. J. Aves, R. G. Beiko, P. Coutinho, J. B. Dacks, D. G. Durnford, N. M. Fast, B. R. Green, C. J. Grisdale, F. Hempel, B. Henrissat, M. P. Höppner, K.-I. Ishida, E. Kim, L. Kořený, P. G. Kroth, Y. Liu, S.-B. Malik, U. G. Maier, D. McRose, T. Mock, J. A. D. Neilson, N. T. Onodera, A. M. Poole, E. J. Pritham, T. A. Richards, G. Rocap, S. W. Roy, C. Sarai, S. Schaack, S. Shirato, C. H. Slamovits, D. F. Spencer, S. Suzuki, A. Z. Worden, S. Zauner, K. Barry, C. Bell, A. K. Bharti, J. A. Crow, J. Grimwood, R. Kramer, E. Lindquist, S. Lucas, A. Salamov, G. I. McFadden, C. E. Lane, P. J. Keeling, M. W. Gray, I. V. Grigoriev, J. M. Archibald, Algal genomes reveal evolutionary mosaicism and the fate of nucleomorphs. Nature 492, 59–65 (2012).

52. D. Dauvillée, P. Deschamps, J.-P. Ral, C. Plancke, J.-L. Putaux, J. Devassine, A. Durand-Terrasson, A. Devin, S. G. Ball, Genetic dissection of floridean starch synthesis in the cytosol of the model dinoflagellate Crypthecodinium cohnii. Proceedings of the National Academy of Sciences 106, 21126–21130 (2009).

53. M. M. Nielsen, C. Ruzanski, K. Krucewicz, A. Striebeck, U. Cenci, S. G. Ball, M. M. Palcic, J. A. Cuesta-Seijo, Crystal Structures of the Catalytic Domain of Arabidopsis thaliana Starch Synthase IV, of Granule Bound Starch Synthase From CLg1 and of Granule Bound Starch Synthase I of Cyanophora paradoxa Illustrate Substrate Recognition in Starch Synthases. Front. Plant Sci. 9, 1138 (2018).

54. L. Eme, S. Sharpe, M. Brown, A. Roger, On the age of eukaryotes: evaluating evidence from fossils and molecular clocks., Cold Spring Harb Perspect Biol. (2014)p. 8.

55. E. C. Cooney, B. S. Leander, P. J. Keeling, Phylogenomics shows unique traits in Noctilucales are derived rather than ancestral. PNAS Nexus 1, pgac202 (2022).

56. S. J. Sibbald, J. M. Archibald, Genomic Insights into Plastid Evolution. Genome Biology and Evolution 12, 978–990 (2020).

57. P. Deschamps, I. Haferkamp, D. Dauvillée, S. Haebel, M. Steup, A. Buléon, J.-L. Putaux, C. Colleoni, C. d’Hulst, C. Plancke, S. Gould, U. Maier, H. E. Neuhaus, S. Ball, Nature of the Periplastidial Pathway of Starch Synthesis in the Cryptophyte *Guillardia theta*. Eukaryot Cell 5, 954–963 (2006).

58. I. Haferkamp, P. Deschamps, M. Ast, W. Jeblick, U. Maier, S. Ball, H. E. Neuhaus, Molecular and Biochemical Analysis of Periplastidial Starch Metabolism in the Cryptophyte *Guillardia theta*. Eukaryot Cell 5, 964–971 (2006).

59. G. I. McFadden, P. R. Gilson, D. R. A. Hill, *Goniomonas* : rRNA sequences indicate that this phagotrophic flagellate is a close relative of the host component of cryptomonads. European Journal of Phycology 29, 29–32 (1994).

60. P. Deschamps, D. Guillebeault, J. Devassine, D. Dauvillée, S. Haebel, M. Steup, A. Buléon, J.-L. Putaux, M.-C. Slomianny, C. Colleoni, A. Devin, C. Plancke, S. Tomavo, E. Derelle, H. Moreau, S. Ball, The Heterotrophic Dinoflagellate *Crypthecodinium cohnii* Defines a Model Genetic System To Investigate Cytoplasmic Starch Synthesis. Eukaryot Cell 7, 872–880 (2008).

61. N. A. Moran, Accelerated evolution and Muller’s rachet in endosymbiotic bacteria. Proc Natl Acad Sci U S A 93, 2873–2878 (1996).

62. S. Ball, D. Baurain, M. Leleu, I. Lafontaine, U. Cenci, C. Colleoni, O. Vallon, G. Greub, A. Weber, D. Bhattacharya, None. Comptes Rendus. Biologies 348, 107–135 (2025).

63. R. Bock, J. N. Timmis, Reconstructing evolution: Gene transfer from plastids to the nucleus. BioEssays 30, 556–566 (2008).

64. A. C. Barbrook, C. J. Howe, S. Purton, Why are plastid genomes retained in non-photosynthetic organisms? Trends in Plant Science 11, 101–108 (2006).

65. D. L. Lister, J. M. Bateman, S. Purton, C. J. Howe, DNA transfer from chloroplast to nucleus is much rarer in Chlamydomonas than in tobacco. Gene 316, 33–38 (2003).

66. N. Lane, Plastids, Genomes, and the Probability of Gene Transfer. Genome Biology and Evolution 3, 372–374 (2011).

67. J. F. Allen, The CoRR hypothesis for genes in organelles. Journal of Theoretical Biology 434, 50–57 (2017).

68. E. Hagström, S. G. Andersson, The challenges of integrating two genomes in one cell. Current Opinion in Microbiology 41, 89–94 (2018).

69. N. Szydlowski, P. Ragel, T. A. Hennen-Bierwagen, V. Planchot, A. M. Myers, A. Mérida, C. d’Hulst, F. Wattebled, Integrated functions among multiple starch synthases determine both amylopectin chain length and branch linkage location in Arabidopsis leaf starch. Journal of Experimental Botany 62, 4547–4559 (2011).

70. P. Deschamps, S. G. Ball, D. Dauvillée, “The comparative, biochemistry, genetics, and evolution of starch metabolism in Chlamydomonas reinhardtii” in The Chlamydomonas Sourcebook (Elsevier, 2023; https://linkinghub.elsevier.com/retrieve/pii/B9780128214305000031), pp. 23–50.

71. N. Libessart, M. L. Maddelein, NVd. Koornhuyse, A. Decq, B. Delrue, G. Mouille, C. D’Hulst, S. Ball, Storage, Photosynthesis, and Growth: The Conditional Nature of Mutations Affecting Starch Synthesis and Structure in Chlamydomonas. Plant Cell, 1117–1127 (1995).

72. D. Bhattacharya, D. C. Price, C. X. Chan, H. Qiu, N. Rose, S. Ball, A. P. M. Weber, M. C. Arias, B. Henrissat, P. M. Coutinho, A. Krishnan, S. Zäuner, S. Morath, F. Hilliou, A. Egizi, M.-M. Perrineau, H. S. Yoon, Genome of the red alga Porphyridium purpureum. Nat Commun 4, 1941 (2013).

73. J. Collén, B. Porcel, W. Carré, S. G. Ball, C. Chaparro, T. Tonon, T. Barbeyron, G. Michel, B. Noel, K. Valentin, M. Elias, F. Artiguenave, A. Arun, J.-M. Aury, J. F. Barbosa-Neto, J. H. Bothwell, F.-Y. Bouget, L. Brillet, F. Cabello-Hurtado, S. Capella-Gutiérrez, B. Charrier, L. Cladière, J. M. Cock, S. M. Coelho, C. Colleoni, M. Czjzek, C. Da Silva, L. Delage, F. Denoeud, P. Deschamps, S. M. Dittami, T. Gabaldón, C. M. M. Gachon, A. Groisillier, C. Hervé, K. Jabbari, M. Katinka, B. Kloareg, N. Kowalczyk, K. Labadie, C. Leblanc, P. J. Lopez, D. H. McLachlan, L. Meslet-Cladiere, A. Moustafa, Z. Nehr, P. Nyvall Collén, O. Panaud, F. Partensky, J. Poulain, S. A. Rensing, S. Rousvoal, G. Samson, A. Symeonidi, J. Weissenbach, A. Zambounis, P. Wincker, C. Boyen, Genome structure and metabolic features in the red seaweed *Chondrus crispus* shed light on evolution of the Archaeplastida. Proc. Natl. Acad. Sci. U.S.A. 110, 5247–5252 (2013).

74. D. C. Price, C. X. Chan, H. S. Yoon, E. C. Yang, H. Qiu, A. P. M. Weber, R. Schwacke, J. Gross, N. A. Blouin, C. Lane, A. Reyes-Prieto, D. G. Durnford, J. A. D. Neilson, B. F. Lang, G. Burger, J. M. Steiner, W. Loffelhardt, J. E. Meuser, M. C. Posewitz, S. Ball, M. C. Arias, B. Henrissat, P. M. Coutinho, S. A. Rensing, A. Symeonidi, H. Doddapaneni, B. R. Green, V. D. Rajah, J. Boore, D. Bhattacharya, Cyanophora paradoxa Genome Elucidates Origin of Photosynthesis in Algae and Plants. Science 335, 843–847 (2012).

75. D. C. Price, U. W. Goodenough, R. Roth, J.-H. Lee, T. Kariyawasam, M. Mutwil, C. Ferrari, F. Facchinelli, S. G. Ball, U. Cenci, C. X. Chan, N. E. Wagner, H. S. Yoon, A. P. M. Weber, D. Bhattacharya, Analysis of an improved Cyanophora paradoxa genome assembly. DNA Research 26, 287–299 (2019).

76. J. Lee, E. C. Yang, L. Graf, J. H. Yang, H. Qiu, U. Zelzion, C. X. Chan, T. G. Stephens, A. P. M. Weber, G. H. Boo, S. M. Boo, K. M. Kim, Y. Shin, M. Jung, S. J. Lee, H.-S. Yim, J.-H. Lee, D. Bhattacharya, H. S. Yoon, Analysis of the Draft Genome of the Red Seaweed Gracilariopsis chorda Provides Insights into Genome Size Evolution in Rhodophyta. Molecular Biology and Evolution 35, 1869–1886 (2018).

77. I. Pancha, K. Tanaka, S. Imamura, Overexpression of a glycogenin, CmGLG2, enhances floridean starch accumulation in the red alga *Cyanidioschyzon merolae*. Plant Signaling & Behavior 14, 1596718 (2019).

78. F. Serafimova, Detection, purification and partial characterization of β-amylase from trophozoites of Entamoeba invadens. Molecular and Biochemical Parasitology 83, 175–181 (1996).

79. C. Plancke, C. Colleoni, P. Deschamps, D. Dauvillée, Y. Nakamura, S. Haebel, G. Ritte, M. Steup, A. Buléon, J.-L. Putaux, D. Dupeyre, C. d’Hulst, J.-P. Ral, W. Löffelhardt, S. G. Ball, Pathway of cytosolic starch synthesis in the model glaucophyte Cyanophora paradoxa. Eukaryot Cell 7, 247–257 (2008).

80. N. J. Patron, P. J. Keeling, COMMON EVOLUTIONARY ORIGIN OF STARCH BIOSYNTHETIC ENZYMES IN GREEN AND RED ALGAE^1^. Journal of Phycology 41, 1131–1141 (2005).

81. Z. Füssy, M. Oborník, “Chromerids and Their Plastids” in Advances in Botanical Research (Elsevier, 2017; https://linkinghub.elsevier.com/retrieve/pii/S0065229617300575)vol. 84, pp. 187–218.

82. S. A. Muñoz-Gómez, C. H. Slamovits, “Plastid Genomes in the Myzozoa” in Advances in Botanical Research (Elsevier, 2018; https://linkinghub.elsevier.com/retrieve/pii/S0065229617300873)vol. 85, pp. 55–94.

83. J. F. H. Strassert, I. Irisarri, T. A. Williams, F. Burki, A molecular timescale for eukaryote evolution with implications for the origin of red algal-derived plastids. Nat Commun 12, 1879 (2021).

