## supplemental figures for "Characterization of starch metabolism pathways in dinoflagellates, chromerids and apicomplexans highlights the presence of a starch accumulating symbiont in their common ancestor"

### SUPPLEMENTARY FIGURES

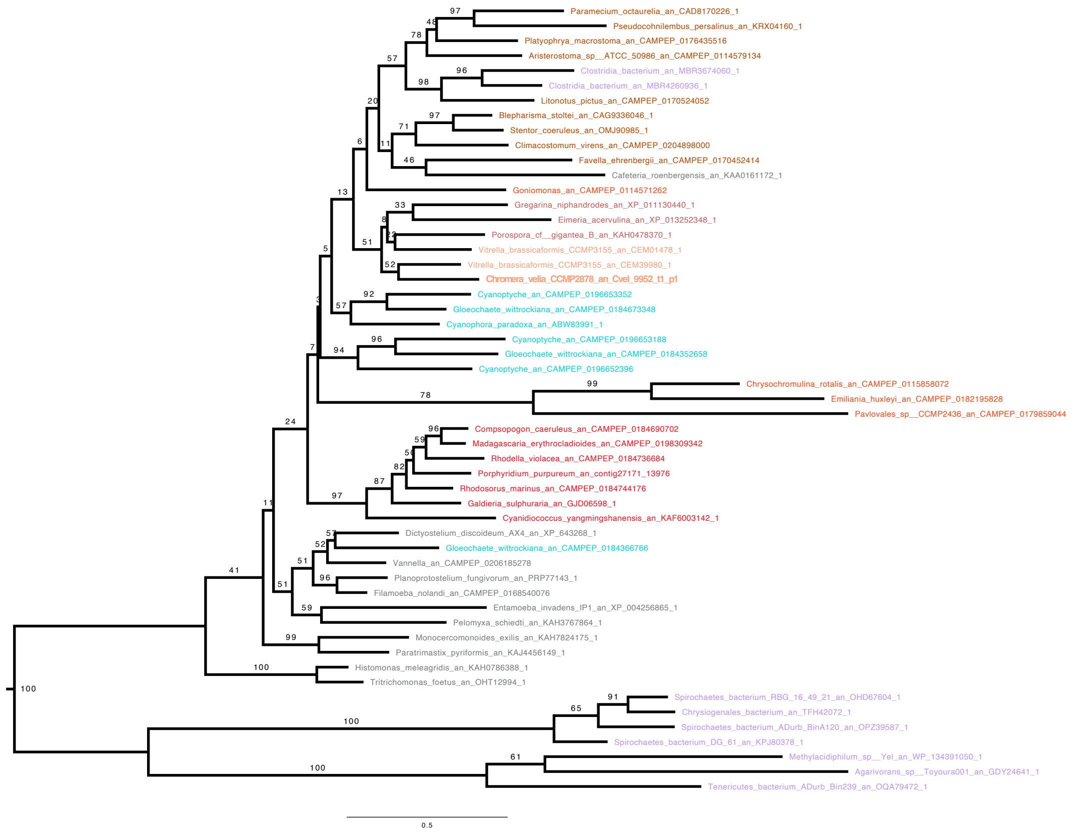


**Figure S1: UDP-glucose Starch/glycogen synthesis (GT5) phylogenetic tree**. The phylogenetic tree was built with IQTREE using LG4X matrix. The bootstrap values are indicated on the branch and are drawn from 100 bootstraps repetition. Sequences are coloured depending on their origin: Rhodophyceae are in Red, Bacteria are in purple, Glaucophyta in light blue, Cryptophyceae in orange, Myzozoa are in light brown except for Chromerids which are in orange and Ciliophora which are in dark brown. Finally other Eukaryotes are in grey. The phylogeny of the GT5 glycogen synthase displays a topology compatible with a monophyletic origin of this enzyme for the ciliates and chromerid/apicomplexa albeit with very low or no support (BS=13). Note the presence of clostridium sequences within the ciliates.

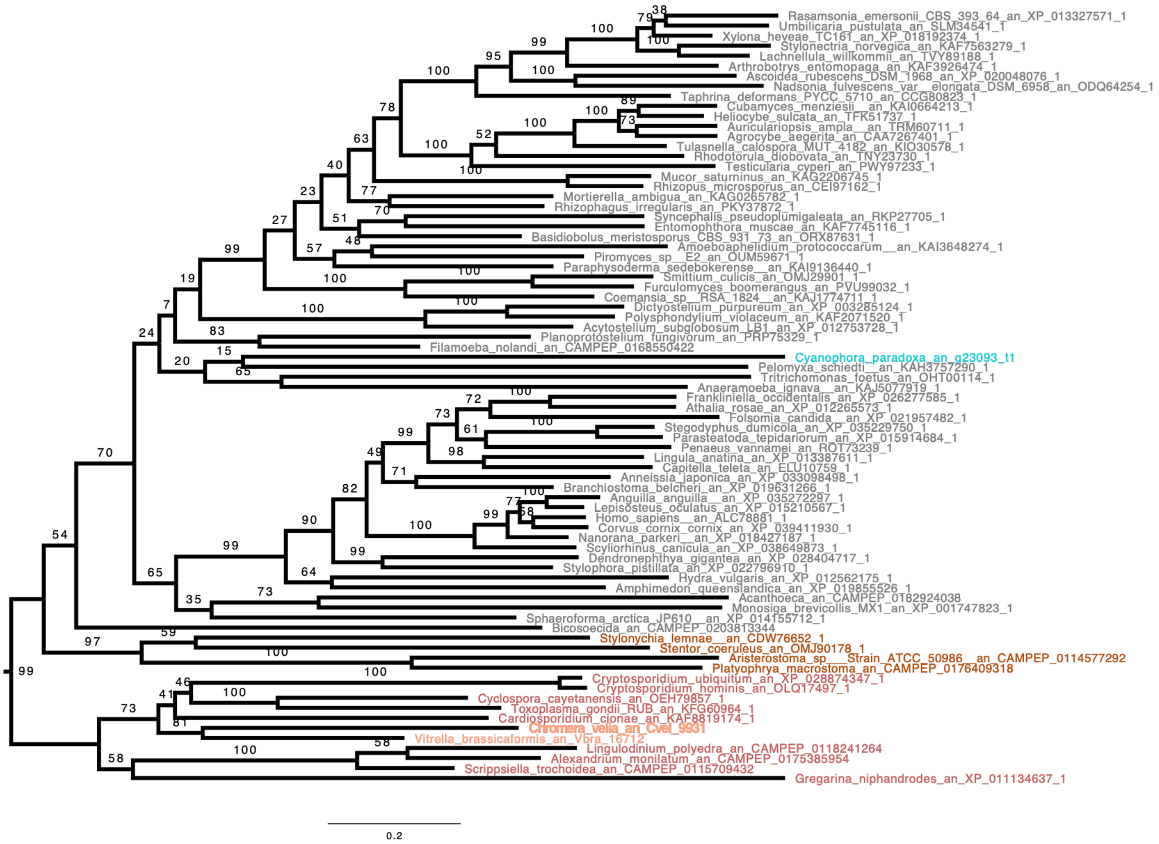


**Figure S2: Indirect debranching enzyme (iDBE) (GH13_25-GH133) phylogenetic tree**. The phylogenetic tree was built with IQTREE using LG4X matrix. The bootstrap values are indicated on the branch and are drawn from 100 bootstrap repetitions. Sequences are coloured depending on their origin: Chloroplastida are in green, Rhodophyceae are in Red, except for Cyanobacteria in black, Bacteria are in purple, Glaucophyta in light blue, Cryptophyceae in orange, Myzozoa are in light brown except for Chromerids which are in light orange and Ciliophora are in dark brown. Finally other Eukaryotes are in grey. *Chromera velia* iDBE groups with other alveolates with BS=54, and with other Myzozoa with BS=99. Interestingly, iDBE is absent from cryptophyceae despite its presence in Goniomonas which begs the question on how such organisms achieve hydrolysis of the α-1,6 linkage. Note the complete absence of prokaryotes in the tree.

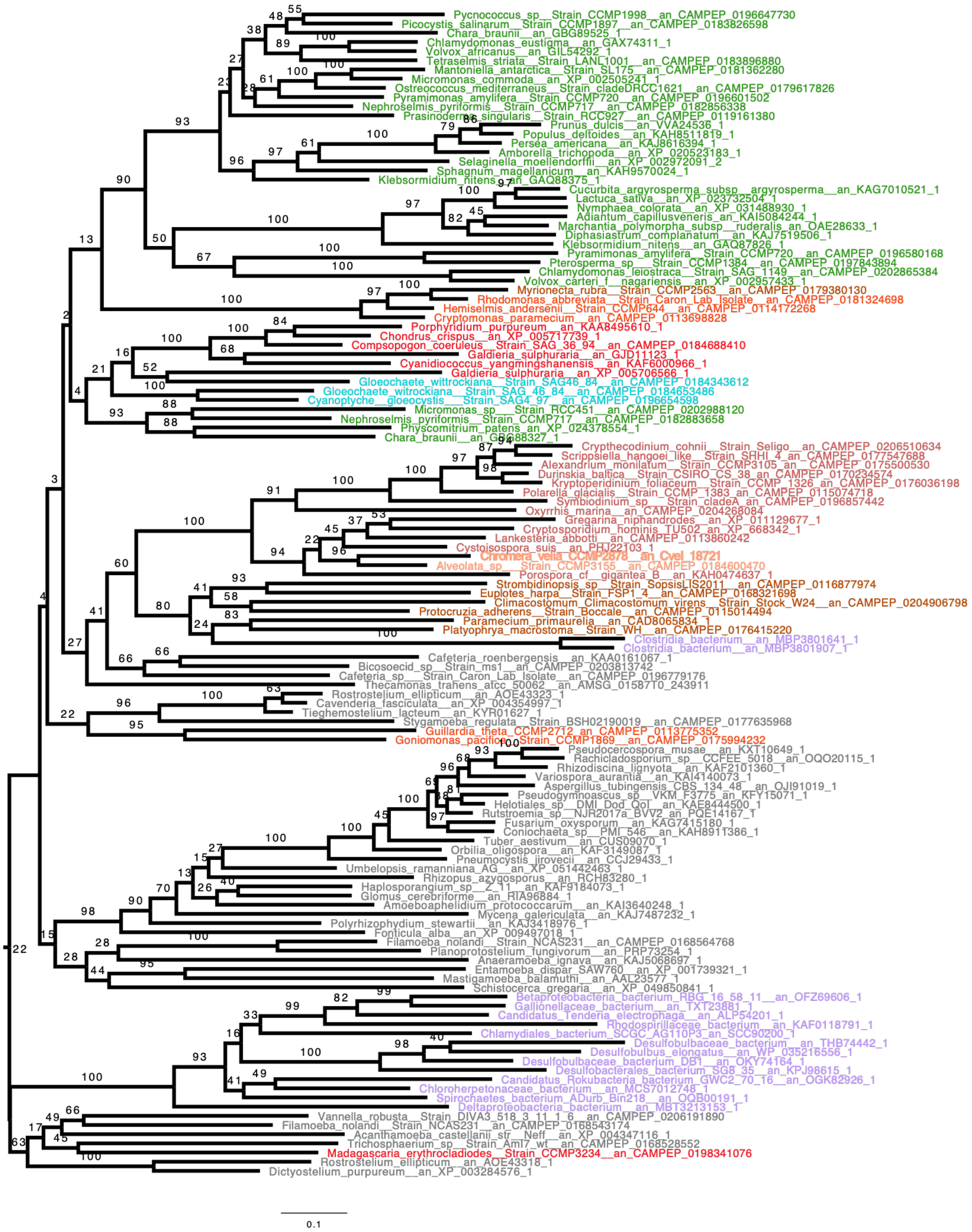


**Figure S3: Starch/glycogen phosphorylase (PHO) (GT35) phylogenetic tree centered on Cvel_18721**. The phylogenetic tree was built with IQTREE using LG4X matrix. The bootstrap values are indicated on the branch and are drawn from 100 bootstraps repetition. Sequences are coloured depending on their origin: Chloroplastida are in green, Rhodophyceae are in Red, except for Cyanobacteria in black, Bacteria are in purple, Glaucophyta in light blue, Cryptophyceae in orange, Myzozoa are in light brown except for Chromerids which are in light orange and Ciliophora are in dark brown. Finally other Eukaryotes are in grey. The phosphorylase in *Chromera velia* is derived from the eukaryote host as the sequence groups with other alveolates, including ciliates, with BS of 60. Interestingly, the cryptophytes equally gained one of its starch phosphorylases from the Goniomonas-like host ancestor. However, a distinct isoform originated from an unresolved archaeplastidal source during the secondary endosymbiosis of cryptophytes. Note the presence of clostridium sequences within the ciliates.

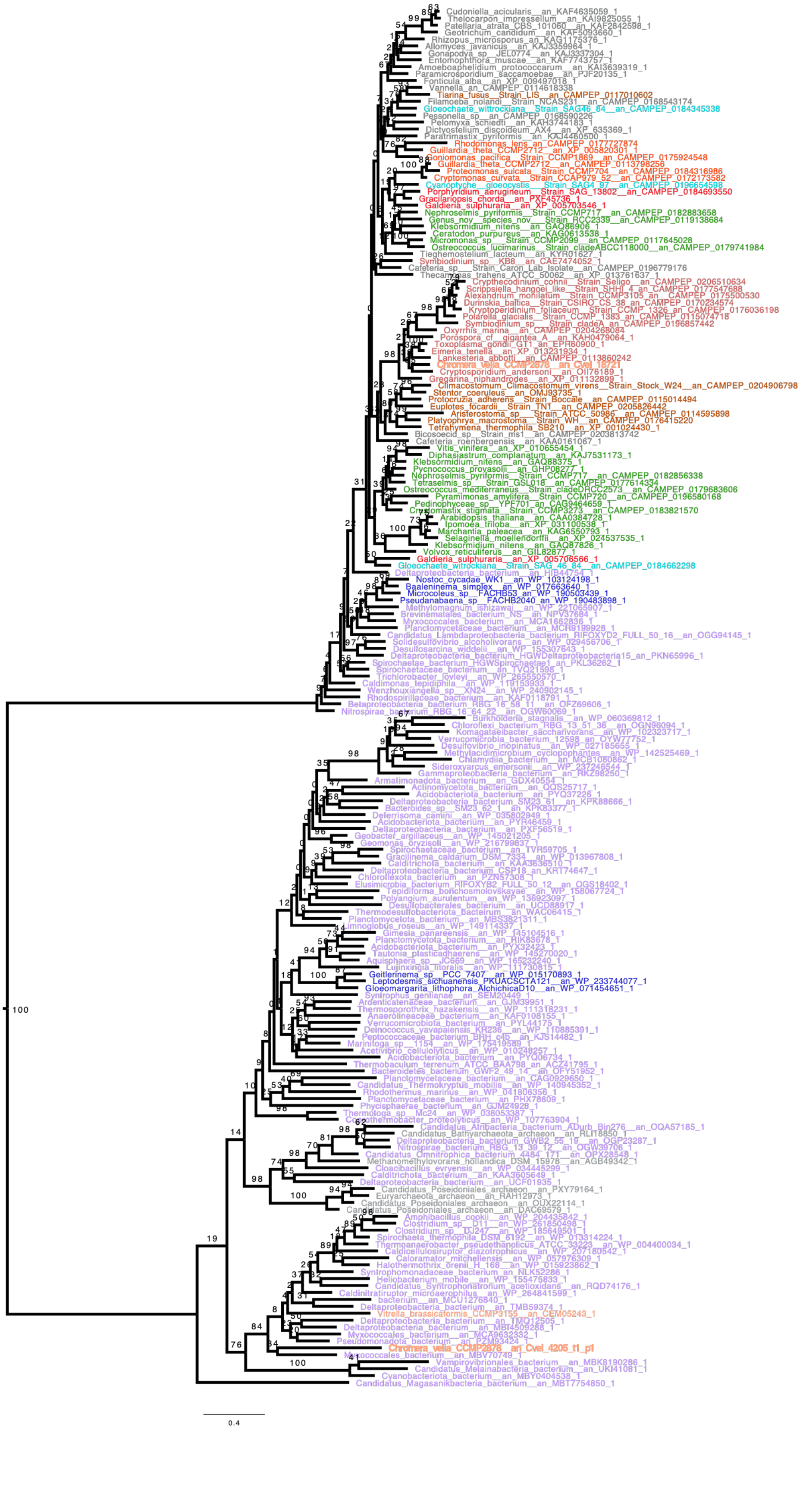

Cudoniella_acicularis__an_KAF4635059_1
Thelocarpon_impressellum__an_KAI9825055_1
Patellaria_atrata_CBS_101060__an_KAF2842598_1
Geotrichum_candidum__an_KAF5093660_1
Rhizopus_microsporus_an_KAG1175376_1
Allomyces_javanicus__an_KAJ3359964_1
Gonapodya_sp__JEL0774__an_KAJ3337304_1
Entomophthora_muscae__an_KAF7743757_1
Amoeboaphelidium_protococcarum__an_KAI3639319_1
Paramicrosporidium_saccamoebae__an_PJF20135_1
Fonticula_alba__an_XP_009497018_1
Vannella_an_CAMPEP_0114618338
Tiarina_fusus__Strain_LIS__an_CAMPEP_0117010602
Filamoeba_nolandi__Strain_NCAS231__an_CAMPEP_0168543174
Gloeochaete_wittrockiana__Strain_SAG46_84__an_CAMPEP_0184345338
Pessonella_sp__an_CAMPEP_0168590226
Pelomyxa_schiedti__an_KAH3744183_1
Dictyostelium_discoideum_AX4__an_XP_635369_1
Paratrimastix_pyriformis__an_KAJ4460500_1
Rhodomonas_lens_an_CAMPEP_0177727874
Guillardia_theta_CCMP2712__an_XP_005820301_1
Goniomonas_pacifica__Strain_CCMP1869__an_CAMPEP_0175924548
Guillardia_theta_CCMP2712__an_CAMPEP_0113798256
Proteomonas_sulcata__Strain_CCMP704__an_CAMPEP_0184316986
Cryptomonas_curvata__Strain_CCAP979_52__an_CAMPEP_0172173582
Cyanoptyche__gloeocystis__Strain_SAG4_97__an_CAMPEP_0196654598
Porphyridium_aerugineum__Strain_SAG_13802__an_CAMPEP_0184693550
Gracilariopsis_chorda__an_PXF45736_1
Galdieria_sulphuraria__an_XP_005703546_1
Nephroselmis_pyriformis__Strain_CCMP717__an_CAMPEP_0182883658
Genus_nov__species_nov__Strain_RCC2339__an_CAMPEP_0119138684
Klebsormidium_nitens__an_GAQ86906_1
Ceratodon_purpureus__an_KAG0613538_1
Micromonas_sp__Strain_CCMP2099__an_CAMPEP_0117645028
Ostreococcus_lucimarinus__Strain_cladeABCC118000__an_CAMPEP_0179741984
Tieghemostelium_lacteum__an_KYR01627_1
Symbiodinium_sp__KB8__an_CAE7474052_1
Cafeteria_sp__Strain_Caron_Lab_Isolate__an_CAMPEP_0196779176
Thecamonas_trahens_ATCC_50062__an_XP_013761837_1
Crypthecodinium_cohnii__Strain_Seligo__an_CAMPEP_0206510634
Scrippsiella_hangoei_like__Strain_SHHI_4_an_CAMPEP_0177547688
Alexandrium_monilatum__Strain_CCMP3105_an__CAMPEP_0175500530
Durinskia_baltica__Strain_CSIRO_CS_38_an_CAMPEP_0170234574
Kryptoperidinium_foliaceum__Strain_CCMP_1326_an_CAMPEP_0176036198
Polarella_glacialis__Strain_CCMP_1383_an_CAMPEP_0115074718
Symbiodinium_sp__Strain_cladeA_an_CAMPEP_0196857442
Oxyrrhis_marina__an_CAMPEP_0204268084
Porospora_cf__gigantea_A__an_KAH0479064_1
Toxoplasma_gondii_GT1_an_EPR60900_1
Eimeria_tenella__an_XP_013231834_1
Lankesteria_abbotti__an_CAMPEP_0113860242
Chromera_velia_CCMP2878__an_Cvel_18721
Cryptosporidium_andersoni__an_OII76189_1
Gregarina_niphandrodes__an_XP_011132899_1
Climacostomum_Climacostomum_virens__Strain_Stock_W24__an_CAMPEP_0204906798
Stentor_coeruleus__an_OMJ93735_1
Protocruzia_adherens__Strain_Boccale__an_CAMPEP_0115014494
Euplotes_focardii__Strain_TN1__an_CAMPEP_0205826442
Aristerostoma_sp__Strain_ATCC_50986__an_CAMPEP_0114595898
Platyophrya_macrostoma__Strain_WH__an_CAMPEP_0176415220
Tetrahymena_thermophila_SB210__an_XP_001024430_1
Bicosoecid_sp__Strain_ms1__an_CAMPEP_0203813742
Cafeteria_roenbergensis__an_KAA0161067_1
Vitis_vinifera__an_XP_010655454_1
Diphasiastrum_complanatum__an_KAJ7531173_1
Klebsormidium_nitens__an_GAQ88375_1
Pycnococcus_provasolii__an_GHP08277_1
Nephroselmis_pyriformis__Strain_CCMP717__an_CAMPEP_0182856338
Tetraselmis_sp__Strain_GSL018__an_CAMPEP_0177614334
Ostreococcus_mediterraneus__Strain_cladeDRCC2573__an_CAMPEP_0179683606
Pyramimonas_amylifera__Strain_CCMP720__an_CAMPEP_0196580168
Pedinophyceae_sp__YPF701_an_CAG9464659_1
Cryptomastix_stigmata__Strain_CCMP3273__an_CAMPEP_0183821570
Arabidopsis_thaliana__an_CAA0384728_1
Ipomoea_triloba__an_XP_031100538_1
Marchantia_paleacea__an_KAG6550793_1
Selaginella_moellendorffii__an_XP_024537535_1
Klebsormidium_nitens__an_GAQ87826_1
Volvox_reticuliferus__an_GIL82877_1
Galdieria_sulphuraria__an_XP_005706566_1
Gloeochaete_wittrockiana__Strain_SAG_46_84__an_CAMPEP_0184662298
Deltaproteobacteria_bacterium__an_HIB44754_1
Nostoc_cycadae_WK1__an_WP_103124198_1
Baaleninema_simplex__an_WP_017663640_1
Microcoleus_sp__FACHB53_an_WP_190503439_1
Pseudanabaena_sp__FACHB2040_an_WP_190483898_1
Methylomagnum_ishizawai__an_WP_221065907_1
Brevinematales_bacterium_NS__an_NPV37684_1
Myxococcales_bacterium__an_MCA1662836_1
Planctomycetaceae_bacterium__an_MCR9199928_1
Candidatus_Lambdaproteobacteria_bacterium_RIFOXYD2_FULL_50_16__an_OGG94145_1
Solidesulfovibrio_alcoholivorans__an_WP_029456706_1
Desulfosarcina_widdelii__an_WP_155307843_1
Deltaproteobacteria_bacterium_HGWDeltaproteobacteria15_an_PKN65996_1
Spirochaetae_bacterium_HGWSpirochaetae1_an_PKL36262_1
Spirochaetaceae_bacterium__an_TVQ21598_1
Trichlorobacter_lovleyi__an_WP_265550570_1
Caldimonas_tepidiphila__an_WP_119153933_1
Wenzhouxiangella_sp__XN24__an_WP_240902145_1
Rhodospirillaceae_bacterium__an_KAF0118791_1
Betaproteobacteria_bacterium_RBG_16_58_11__an_OFZ69606_1
Nitrospirae_bacterium_RBG_16_64_22__an_OGW60069_1
Burkholderia_stagnalis__an_WP_060369812_1
Chloroflexi_bacterium_RBG_13_51_36__an_OGN96094_1
Komagataeibacter_saccharivorans__an_WP_102323717_1
Verrucomicrobia_bacterium_12598__an_OYW77752_1
Desulfovibrio_inopinatus__an_WP_027185655_1
Methylacidimicrobium_cyclopophantes__an_WP_142525469_1
Chlamydia_bacterium__an_MCB1080862_1
Sideroxyarcus_emersonii__an_WP_237246544_1
Gammaproteobacteria_bacterium__an_RKZ98250_1
Armatimonadota_bacterium__an_GDX40554_1
Actinomycetota_bacterium__an_QQS25717_1
Acidobacteriota_bacterium__an_PYQ37226_1
Deltaproteobacteria_bacterium_SM23_61__an_KPK88666_1
Bacteroides_sp__SM23_62_1__an_KPK83377_1
Deferrisoma_camini__an_WP_035802949_1
Acidobacteriota_bacterium__an_PYR46459_1
Deltaproteobacteria_bacterium__an_PXF56519_1
Geobacter_argillaceus__an_WP_145021205_1
Geomonas_oryzisoli__an_WP_216799837_1
Spirochaetaceae_bacterium__an_TVR59705_1
Gracilinema_caldarium_DSM_7334__an_WP_013967808_1
Calditrichota_bacterium__an_KAA3636510_1
Deltaproteobacteria_bacterium_CSP18_an_KRT74647_1
Chloroflexota_bacterium__an_PZN57308_1
Elusimicrobia_bacterium_RIFOXYB2_FULL_50_12__an_OGS18402_1
Tepidiforma_bonchosmolovskayae__an_WP_158067724_1
Polyangium_aurulentum__an_WP_136923097_1
Desulfobacterales_bacterium__an_UCD88917_1
Thermodesulfobacteriota_bacterium__an_WAC06415_1
Planctomycetota_bacterium__an_MBS3821311_1
Limnoglobus_roseus__an_WP_149114337_1
Gimesia_panareensis__an_WP_145104516_1
Planctomycetota_bacterium__an_RIK63678_1
Acidobacteriota_bacterium__an_PYX32423_1
Tautonia_plasticadhaerens__an_WP_145270020_1
Aquisphaera_sp__JC669__an_WP_165232240_1
Lujinxingia_litoralis__an_WP_111730815_1
Geitlerinema_sp__PCC_7407__an_WP_015170893_1
Leptodesmis_sichuanensis_PKUACSCTA121__an_WP_233744077_1
Gloeomargarita_lithophora_AlchichicaD10__an_WP_071454651_1
Syntrophus_gentianae__an_SEM20449_1
Ardenticatenaceae_bacterium__an_GJM39951_1
Thermosporothrix_hazakensis__an_WP_111318231_1
Anaerolineaceae_bacterium__an_KAF0106155_1
Verrucomicrobiota_bacterium__an_PYL44175_1
Deinococcus_yavapaiensis_KR236__an_WP_110885391_1
Peptococcaceae_bacterium_BRH_c4b__an_KJS14482_1
Marinitoga_sp__1154__an_WP_175419589_1
Acetivibrio_cellulolyticus__an_WP_010248257_1
Acidobacteriota_bacterium__an_PYQ06734_1
Thermobaculum_terrenum_ATCC_BAA798_an_ACZ41795_1
Bacteroidetes_bacterium_GWF2_49_14__an_OFY51952_1
Planctomycetaceae_bacterium__an_CAG0929650_1
Candidatus_Thermokryptus_mobilis__an_WP_140945352_1
Rhodothermus_marinus__an_WP_041806356_1
Planctomycetaceae_bacterium__an_PHX78609_1
Phycisphaerae_bacterium__an_GJM24928_1
Thermotoga_sp__Mc24__an_WP_038053387_1
Coprothermobacter_proteolyticus__an_WP_107763904_1
Candidatus_Atribacteria_bacterium_ADurb_Bin276__an_OQA57185_1
Candidatus_Bathyarchaeota_archaeon__an_RLI18850_1
Deltaproteobacteria_bacterium_GWB2_55_19__an_OGP23287_1
Nitrospirae_bacterium_RBG_13_39_12__an_OGW39706_1
Candidatus_Omnitrophica_bacterium_4484_171__an_OFX28548_1
Methanomethylovorans_hollandica_DSM_15978__an_AGB49342_1
Cloacibacillus_evryensis__an_WP_034445299_1
Calditrichota_bacterium__an_KAA3605649_1
Deltaproteobacteria_bacterium__an_UCF01935_1
Candidatus_Poseidoniales_archaeon__an_PXY79164_1
Euryarchaeota_archaeon__an_RAH12973_1
Candidatus_Poseidoniales_archaeon__an_OUX22114_1
Candidatus_Poseidoniales_archaeon__an_DAC69579_1
Amphibacillus_cookii__an_WP_204435842_1
Clostridium_sp__D11__an_WP_261850498_1
Clostridium_sp__DJ247__an_WP_185649501_1
Spirochaeta_thermophila_DSM_6192__an_WP_013314224_1
Thermoanaerobacter_pseudethanolicus_ATCC_33223__an_WP_004400034_1
Caldicellulosiruptor_diazotrophicus__an_WP_207180542_1
Caloramator_mitchellensis__an_WP_057976309_1
Halothermothrix_orenii_H_168__an_WP_015923862_1
Syntrophomonadaceae_bacterium_an_NLK52286_1
Heliobacterium_mobile__an_WP_155475833_1
Candidatus_Syntrophonatronum_acetioxidans__an_RQD74176_1
Caldinitratiruptor_microaerophilus__an_WP_264841599_1
bacterium__an_MCU1276840_1
Deltaproteobacteria_bacterium__an_TMB59374_1
Vitrella_brassicaformis_CCMP3155__an_CEM05243_1
Deltaproteobacteria_bacterium__an_TMQ12505_1
Deltaproteobacteria_bacterium__an_MBI4509288_1
Myxococcales_bacterium__an_MCA9632332_1
Pseudomonadota_bacterium__an_PZM83424_1
Chromera_velia_CCMP2878__an_Cvel_4205_t1_p1
Myxococcales_bacterium__an_MBV70749_1
Vampirovibrionales_bacterium__an_MBK8190286_1
Candidatus_Melainabacteria_bacterium__an_UKI41081_1
Cyanobacteriota_bacterium__an_MBY0404538_1
Candidatus_Magasanikbacteria_bacterium__an_MBT7754850_1
0.4

**Figure S4: Starch/glycogen phosphorylase (PHO) (GT35) phylogenetic tree displaying all *C. velia* proteins.** The phylogenetic tree was built with IQTREE using LG4X matrix. The bootstrap values are indicated on the branch and are drawn from 100 bootstraps repetition. Sequences are coloured depending on their origin: Chloroplastida are in green, Rhodophyceae are in Red, except for Cyanobacteria in black, Bacteria are in purple, Glaucophyta in light blue, Cryptophyceae in orange, Myzozoa are in light brown except for Chromerids which are in light orange and Ciliophora are in dark brown. Finally other Eukaryotes are in grey. The two GT35 from *C. velia* are present in the tree and indicates separated origins. GT35 from Cvel_4205 displays a bacterial origin.

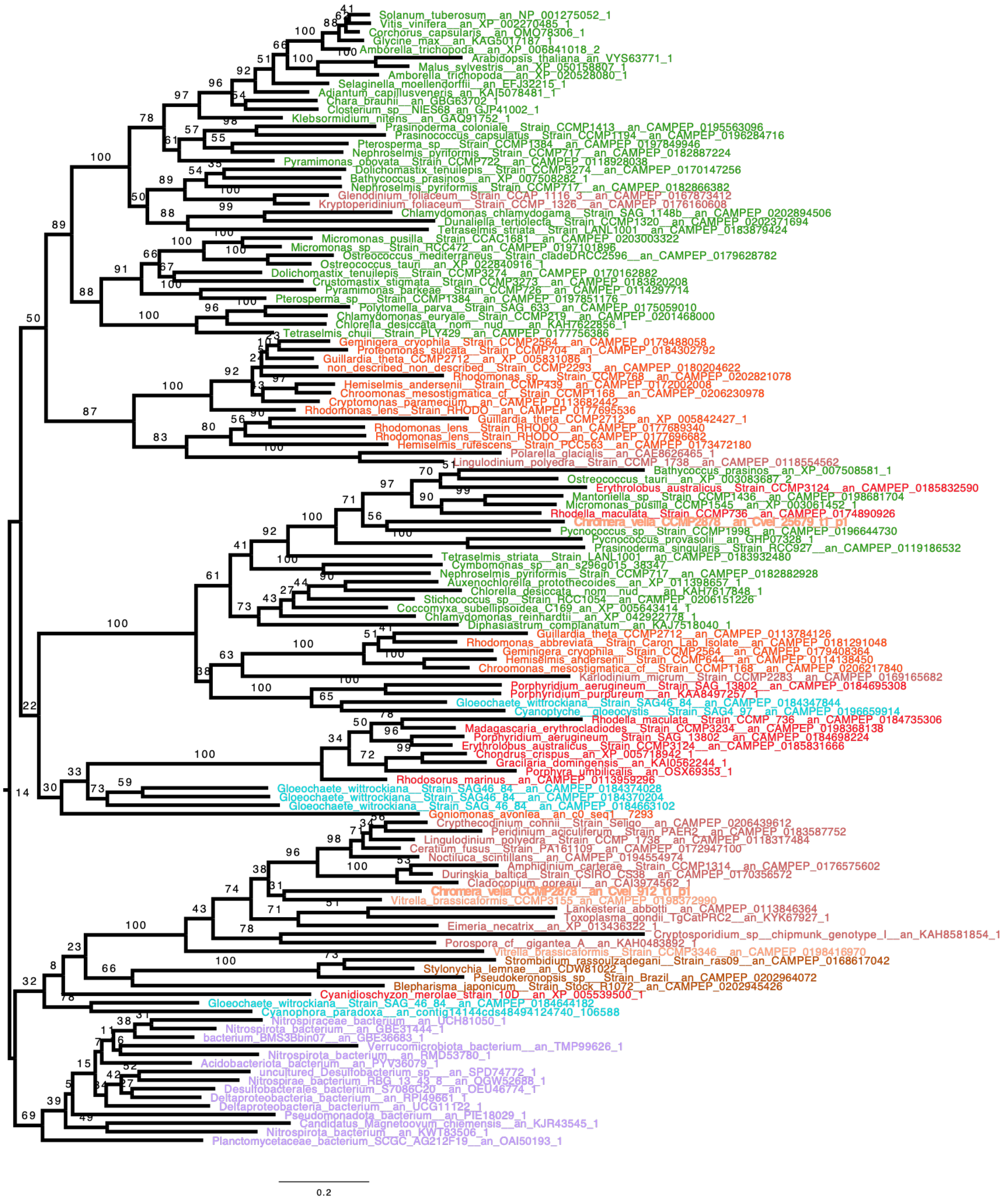


**Figure S5: GWD (CBM45-PPG) phylogenetic tree.** The phylogenetic tree was built with IQTREE using LG4X matrix. The bootstrap values are indicated on the branch and are drawn from 100 bootstrap repetitions (BS). Sequences are coloured depending on their origin: Chloroplastida are in green, Rhodophyceae are in Red, Bacteria are in purple, Glaucophyta in light blue, Cryptophyceae in orange, Myzozoa are in light brown except for Chromerids which are in light orange and Ciliophora which are in dark brown. Finally other Eukaryotes are in grey. In this single gene phylogeny, we observe a strong monophyly of most of the Dinoflagellates (BS=100) and a weaker monophyly that retrieves most of the alveolates together (BS=23). We nevertheless suggest a common archaeplastidal (red alga or glaucophyte) origin for this alveolate enzyme (BS=32). Note that the cryptophytes can be excluded as a possible source for the alveolate common GWD.

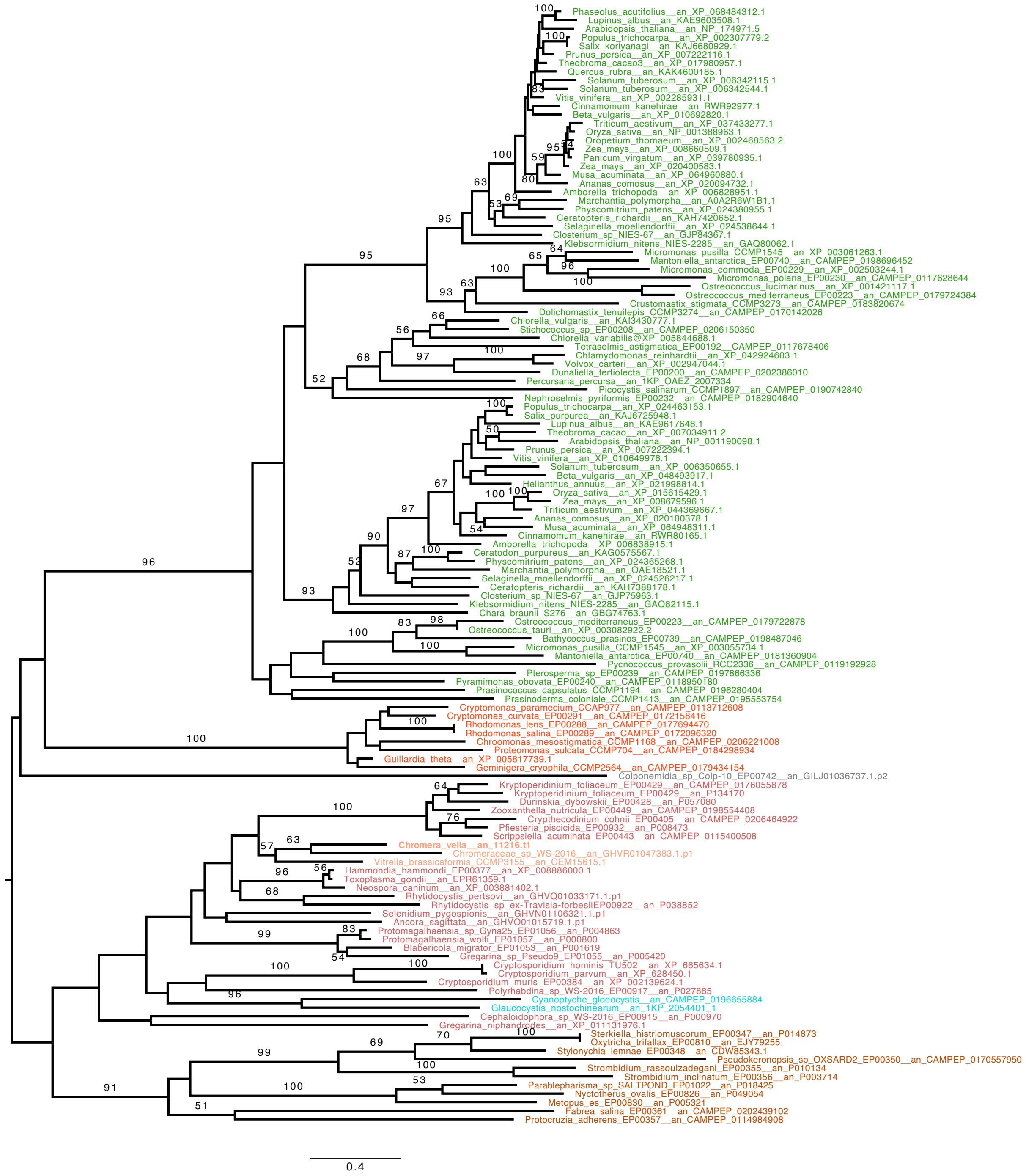


**Figure S6: ESV-LESV phylogeny. ESV-LESV phylogeny.** The phylogenetic tree was built with IQTREE using 100 bootstraps repetition with LG4X matrix. Sequences are coloured depending on their origin: Chloroplastida are in green, Rhodophyceae are in Red, except for Cyanobacteria in dark blue, Bacteria are in purple, Glaucophyta in light blue, Cryptophyceae in orange, Myzozoa are in light brown except for Chromerids which are in light orange. ESV-like sequences are found in *Chromera velia* and other dinoflagellate and apicomplexa and appear monophyletic with good support (BS=97). We observe the absence of these polysaccharide chaperones in all red algae as well as paraglycogen (amylopectin) accumulating ciliates. Interestingly these proteins are found in cryptophytes where they presently define the only candidate protein sequences that could influence starch crystallinity. ESV-LESV has possibly been gained by Myzozoa from the more closely related sequences of Glaucophyta (BS=64). However, more distantly related cryptophyte or green alga sources cannot be discounted.

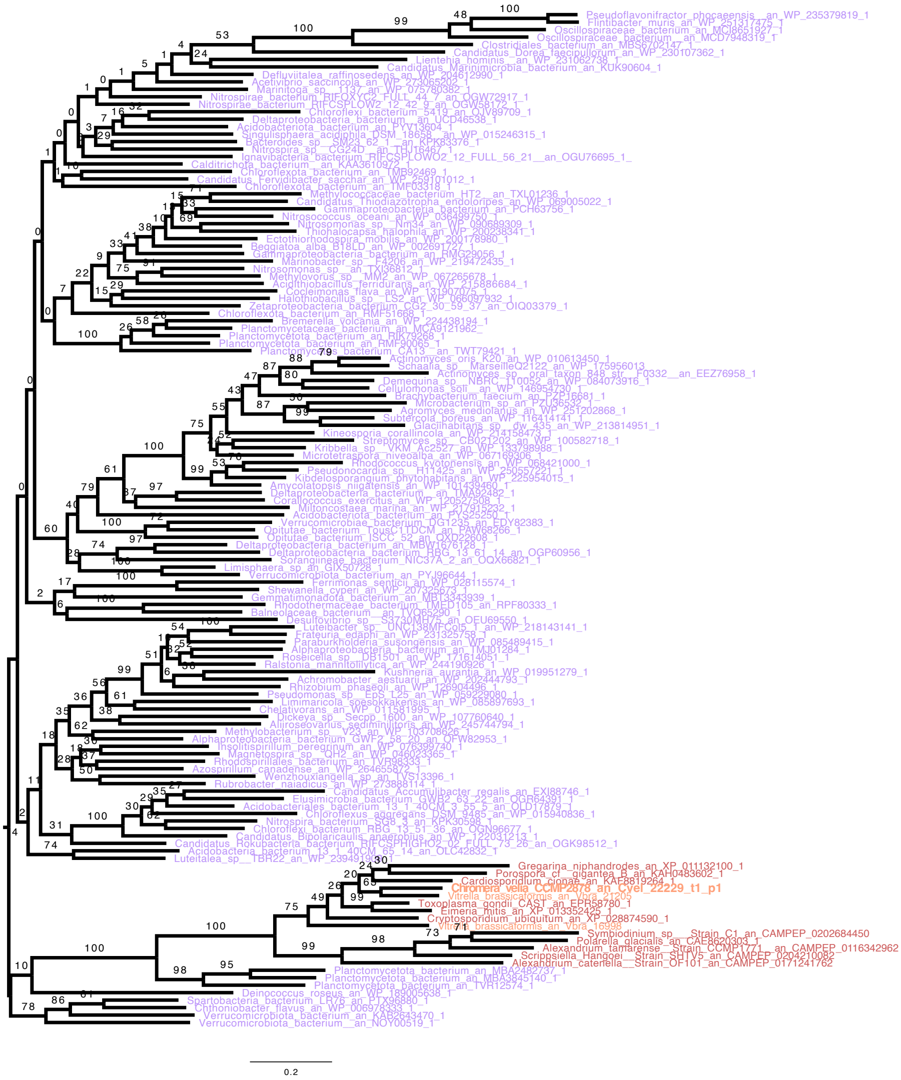

**Figure S7: Branching enzyme 1 (BE1) (GH13_9) phylogenetic tree.** The phylogenetic tree displayed here was built with IQTREE using LG4X matrix. The bootstrap values are indicated on the branch and are drawn from 100 boostrap repetitions (BS). Sequences are coloured depending on their origin: Bacteria are in purple, Myzozoa are in light brown except for Chromerids which are in orange. We observe a bacterial LGT encoding BE1 (GH13_9) possibly acquired from planctomycetes (BS=100) known to fulfil the essential branching enzyme function in Myzozoa as demonstrated in *Toxoplasma gondii* (*1*) and confirmed by our mutant selection in *Chromera velia* (Chabi et al submitted). Interestingly, the presence of BE1 correlates tightly in all alveolates with a remodelling of the BE2 locus by an insertion of a GT4 type of glycosyl transferase sequence (Fig. S6).

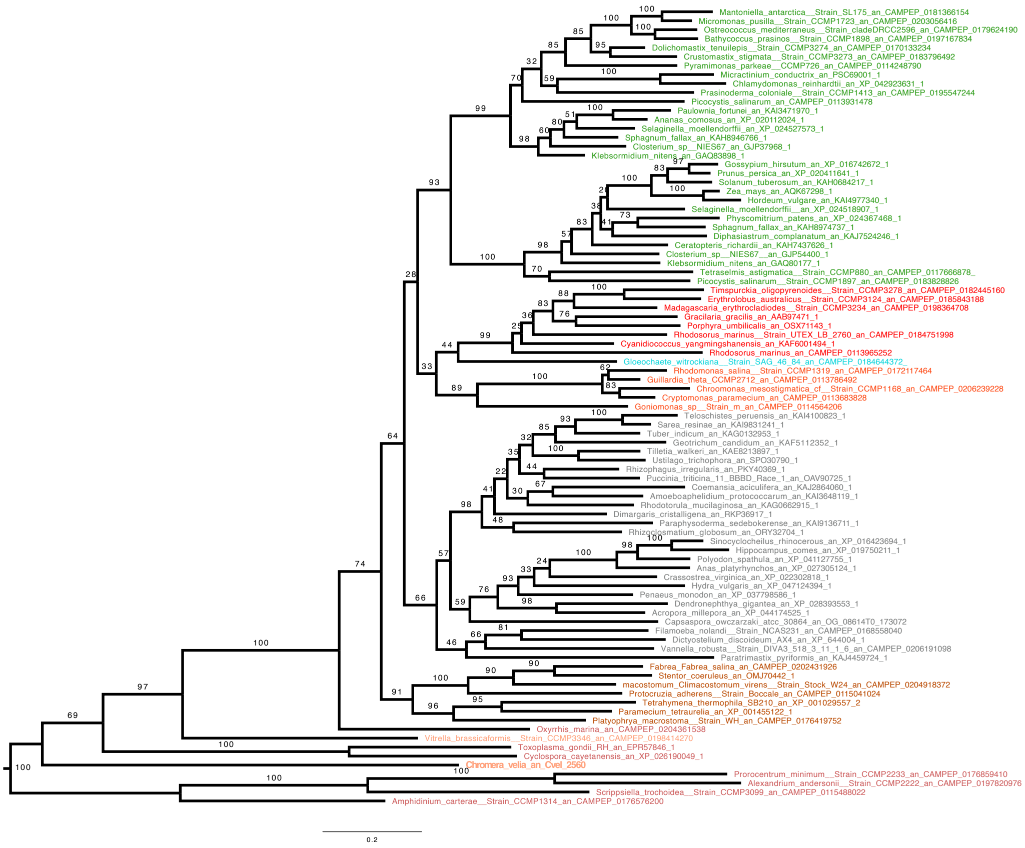


**Figure S8: Branching enzyme 2 (BE2) (GH13_8) phylogenetic tree.** The phylogenetic tree was built with IQTREE using LG4X matrix. The bootstrap values are indicated on the branch and are drawn from 100 boostraps repetition (BS). Sequences are coloured depending on their origin: Chloroplastida are in green, Rhodophyceae are in Red, except for Cyanobacteria in black, Glaucophyta in light blue, Cryptophyceae in orange, Myzozoa are in light brown except for Chromerids which are in orange and Ciliophora are in dark brown. Finally other Eukaryotes are in grey. The BE2 in myzoza is interrupted by a GT4 transferase separating the N-terminal from the C-terminal parts and the presence of the CBM48, however, have not changed. This probably led to neo/subfunctionalization thereby explaining the long branch of this enzyme found at the bottom of the tree, however and except for Oxyrrhis where we have detected a distinct fusion organization involving the dinoflagellate-specific GT5 starch synthase in place of the GT4 glycosyl transferase, all myzozoa group together with a BS=100, indicating a single event of gene fusion. We do not know if Oxyrrhis contains both types of fusion organisations.

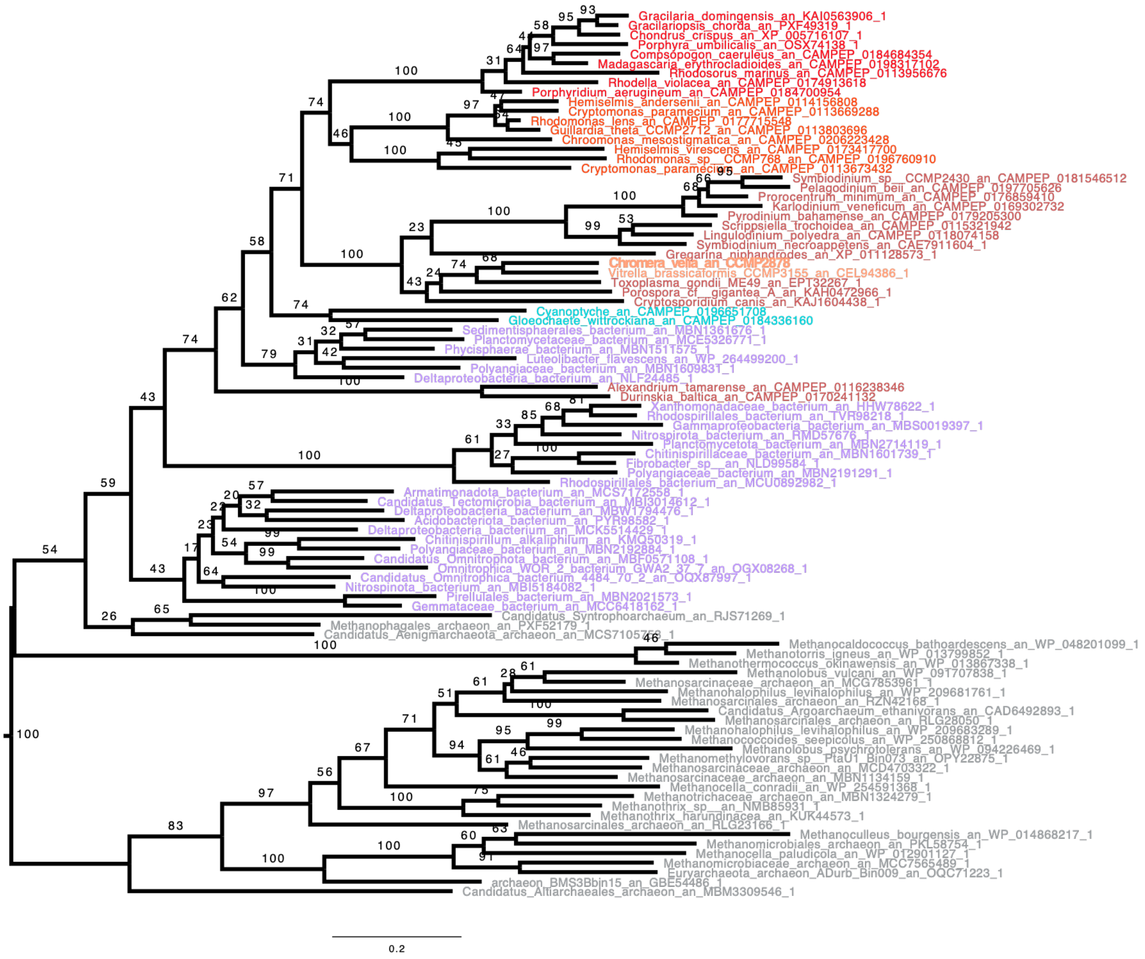


**Figure S9: GT4 phylogenetic tree.** The phylogenetic tree was built with IQTREE using LG4X matrix. The bootstrap values are indicated on the branch and are drawn from 100 bootstrap repetitions. Sequences are coloured depending on their origin: Rhodophyceae are in red, Archae grey, Bacteria are in purple, Glaucophyta in light blue, Cryptophyceae are in light red and Rhodophyceae are in red, Dinoflagellates and Apicomplexa are in dark red and Chromerids are in orange with Chromera in bold. The phylogeny and distribution of the GT4 sequence inserted into the former alveolate BE2 structural gene is never found in ciliates and can be seen as an LGT to Myzozoa. However, insufficient resolution prevents to make definitive statements as to the possible gene donors to the myzozoa but does restrict them to 3 possibilities: glaucophytes, red algae or cryptophytes.

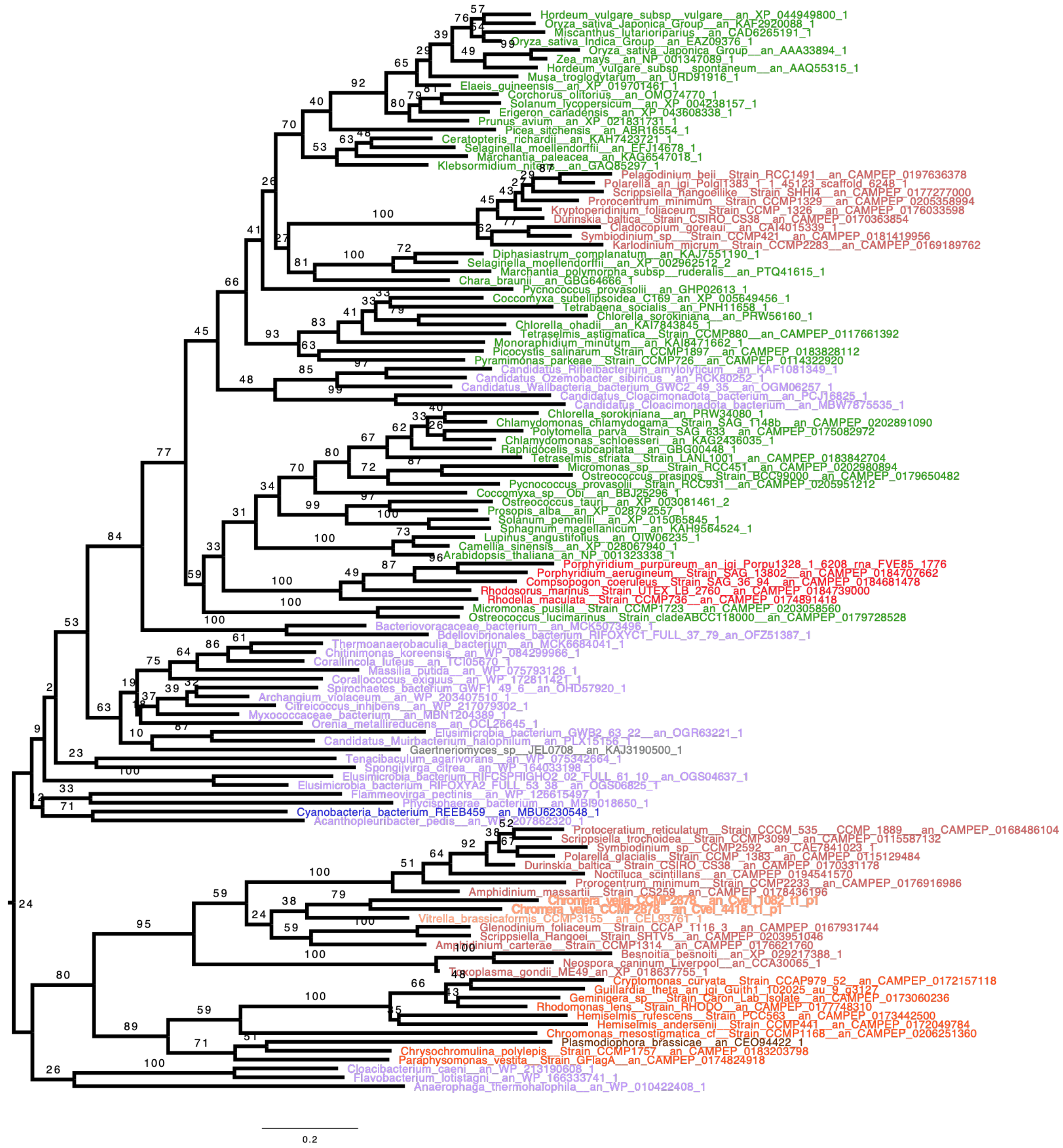

**Figure S10: α-amylase (GH13_6) phylogenetic tree.** The phylogenetic tree displayed here was built with IQTREE using LG4X matrix. The bootstrap values are indicated on the branch and are drawn from 100 bootstrap repetitions. Sequences are coloured depending on their origin: Chloroplastida are in green, Rhodophyceae are in Red, except for Cyanobacteria in dark blue, Bacteria are in purple, Glaucophyta in light blue, Cryptophyceae in orange, Myzozoa are in light brown except for Chromerids which are in light orange and Ciliophora which are in dark brown. Finally other Eukaryotes are in grey. The *Chromera velia* sequence is found with other Myzozoa which all branch as a sister group of Cryptophyceae with good support, thus this sequence seems to have been inherited by either an unidentified bacterial or more likely a cryptophycean source. In addition, we observe the presence of a group of α-amylase from the GH13_6 family acquired from green algae which distributes selectively in dinoflagellates in addition to the common myzozoan isoform.

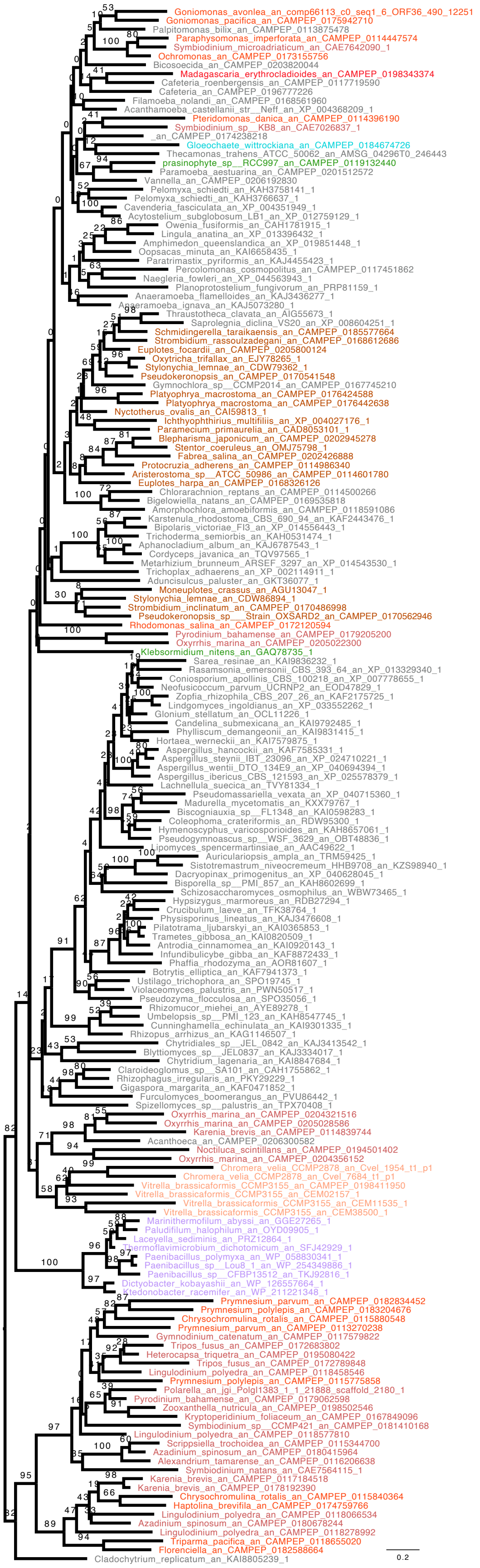

**Figure S11: α-amylase (GH13_1) phylogenetic tree**. The phylogenetic tree was built with IQTREE using LG4X matrix. The bootstrap values are indicated on the branch and are drawn from 100 bootstrap repetitions. Sequences are coloured depending on their origin: Chloroplastida are in green, Rhodophyceae are in Red, Bacteria are in purple, Glaucophyta in light blue, Cryptophyceae in orange, Myzozoa are in light brown except for Chromerids which are in light orange and Ciliophora which are in dark brown. Finally other Eukaryotes are in grey. The α-amylase from Chromera is shared with Vitrella but also other and early diverging Dinoflagellates and is most likely from unknown eukaryotic origin. Several isoforms of GH13_1 sequences are witnessed in the tree including eukaryotic clades not known to accumulate glycogen or starch such as as Rhizaria, haptophytes or percolozoa arguing for some function of this isoform in prey digestion. Of note is the large distribution of a form of dinoflagellate GH13_1 which are exclusively related to haptophyte sequences.

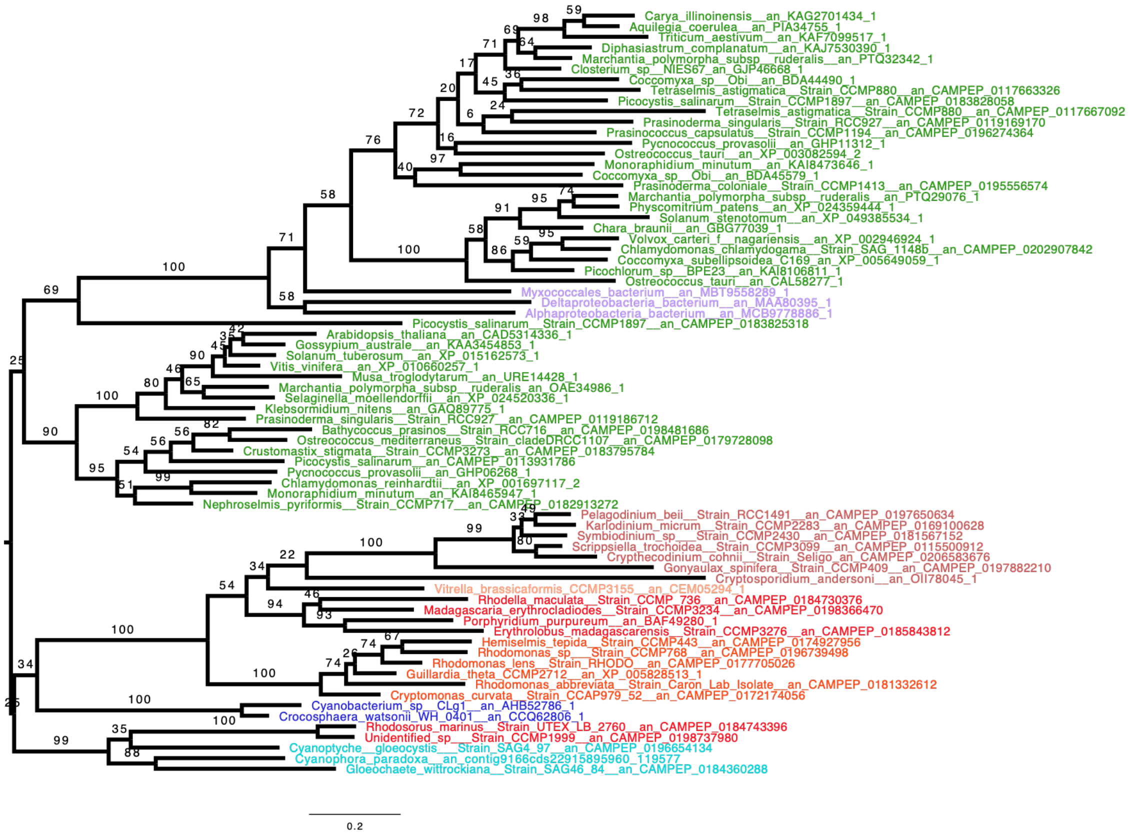


**Figure S12: GBSS (GT5) phylogenetic tree.** The phylogenetic tree was built with IQTREE using LG4X matrix. The bootstrap values are indicated on the branch and are drawn from 100 bootstrap repetitions. Sequences are coloured depending on their origin: Chloroplastida are in green, Rhodophyceae are in Red, except for Cyanobacteria in dark blue, Bacteria are in purple, Glaucophyta in light blue, Cryptophyceae in orange, Myzozoa are in light brown except for Chromerids which are in light orange and Ciliophora which are in dark brown. Finally other Eukaryotes are in grey. GBSS are found in dinoflagellates as well as in the chromerid *Vitrella brassicaformis*, and a few apicomplexa (*e.g.* Cryptosporidium). *Chromera velia* and several apicomplexa store amylopectin granules devoid of GBSS and consequently of amylose. Interestingly, this phylogenetic tree suggests the presence of an analogous red alga source for this GBSS and those of dinoflagellates and cryptophytes (BS=100). Despite its affiliation with red algae (BS=54), we cannot exclude a possible cryptophycean source for the myzozoan enzyme (*2*).

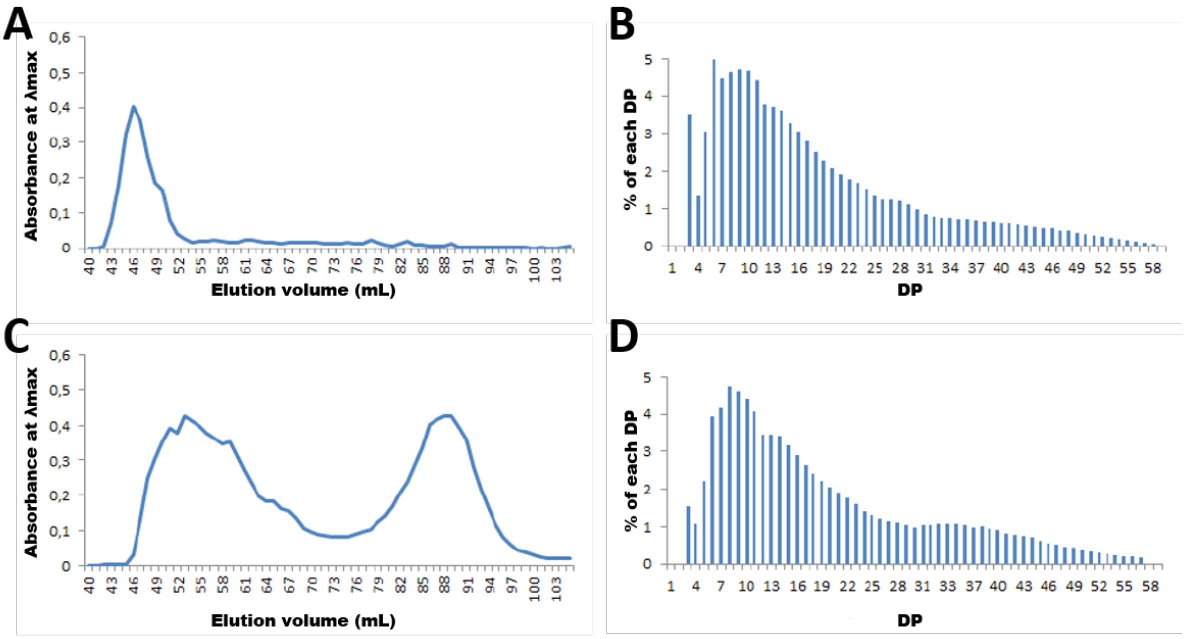


**Figure S13: Structural characterization of *C. velia* and *V. brassicaformis* storage polysaccharides.** For this experiment we performed size exclusion chromatography using TSKHW75 columns on starch solubilized in 10% DMSO from *C. velia* **(A)** and *V. brassicaformis* **(C)**. At variance with *C. velia*, we observe using *Vitrella brassicaformis* starch granules the presence of smaller size amylose eluting as a second polysaccharide peak with iodine staining properties identical to plant amylose. The fractions containing amylopectin were pooled and a CLD (Chain-length distribution) using HPAEC-PAD analysis was performed for *C. velia* **(B)** and *V. brassicaformis* **(D)**.

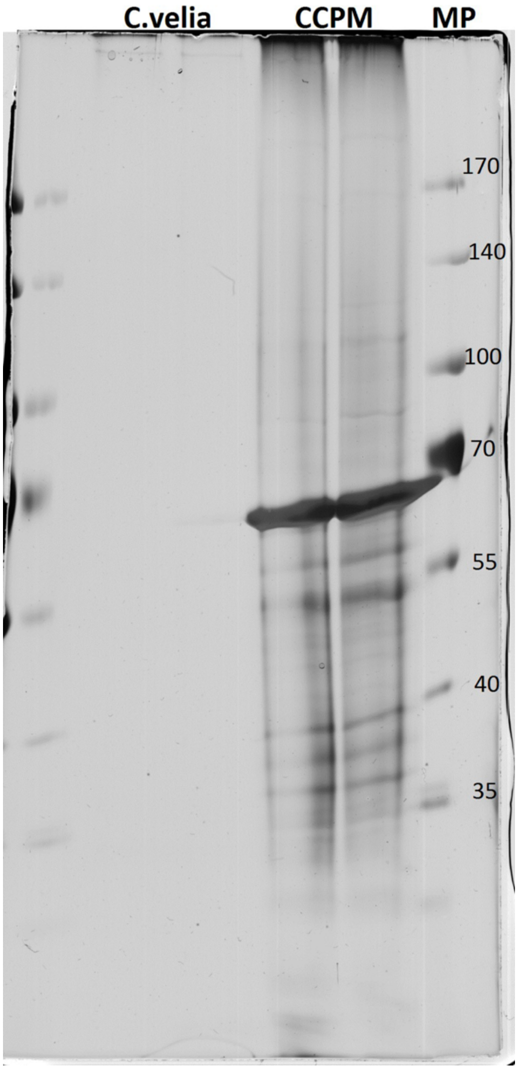


**Figure S14: Picture of SDS-PAGE gel from starch granule extraction**. CV :*Chromera velia*, CCPM : *Vitrella brassicaformis*, MP : *Weight marker*. Proteins extracted from starch granules were denatured and separated on a SDS-PAGE gel. We can see a clear band for *V. brassicaformis* of around 60kDa that corresponds to the expected size of GBSS.

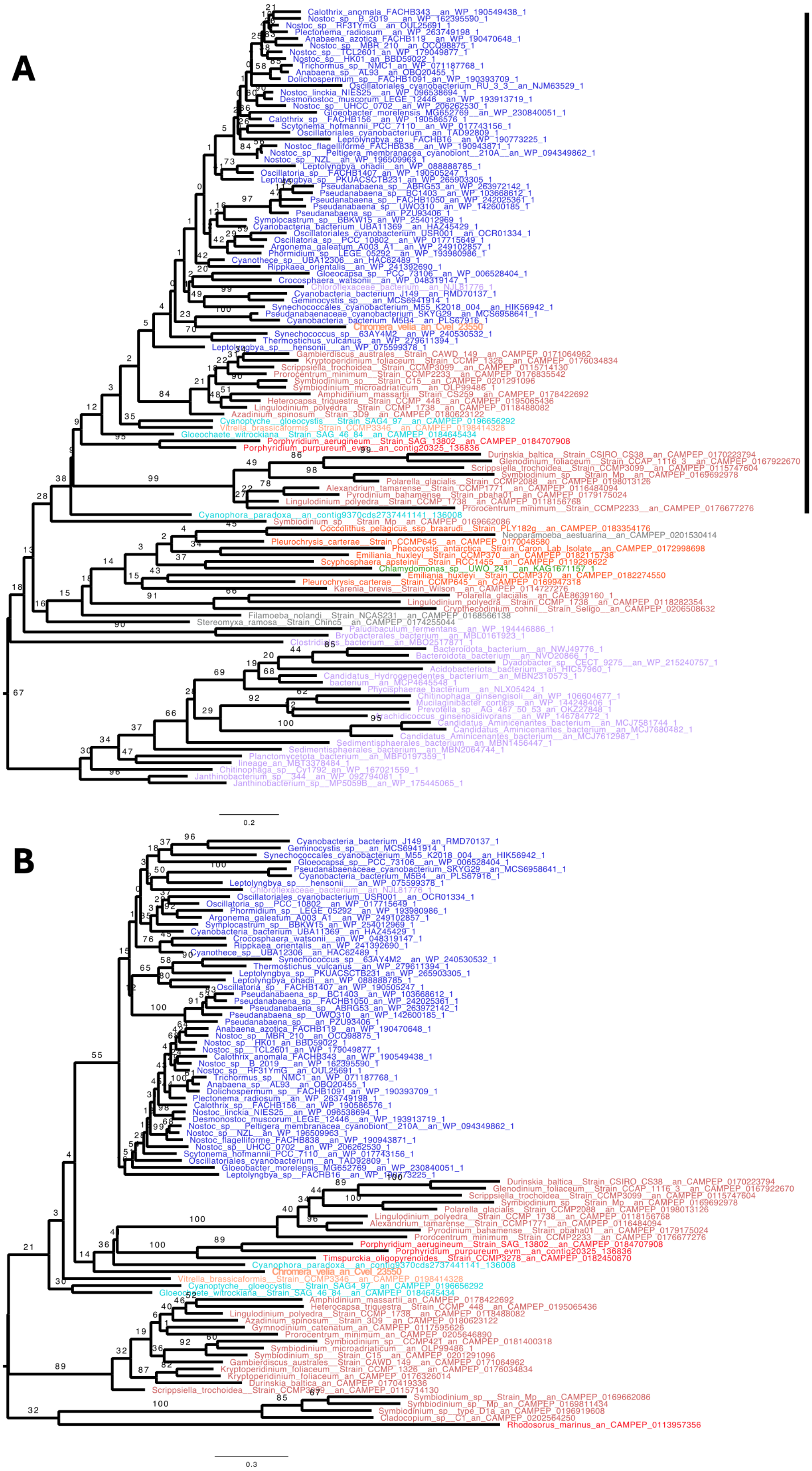


**Figure S15: CBM20-SGNH phylogenetic tree.** The phylogenetic tree was built with IQTREE using LG4X matrix. The bootstrap values are indicated on the branch and are drawn from 100 bootstrap repetitions. Sequences are coloured depending on their origin: Chloroplastida are in green, Rhodophyceae are in Red, Cyanobacteria in dark blue, Bacteria are in purple, Glaucophyta in light blue,

Myzozoa are in light brown except for Chromerids which are in orange. Finally other Eukaryotes are in grey. **(A)** The distribution of this protein among alveolates seems limited to the dinoflagellates and chromerids where it assumes an essential function in starch synthesis that has not been maintained in apicomplexa parasites. All eukaryote strains that contain both the CBM20 and SGHN domains in the corresponding protein are unicellular and photosynthetic and accumulate starch in the cytosol. In addition, we failed to recover this protein within the available dinoflagellate heterotrophic clade sequences. All eukaryotes and non-cyanobacterial prokaryotes displayed here do not contain the CBM20 domain **(B)** To better appreciate relationship between these sequences we also built a phylogenetic tree focussed on all sequences containing the CBM20 domain.

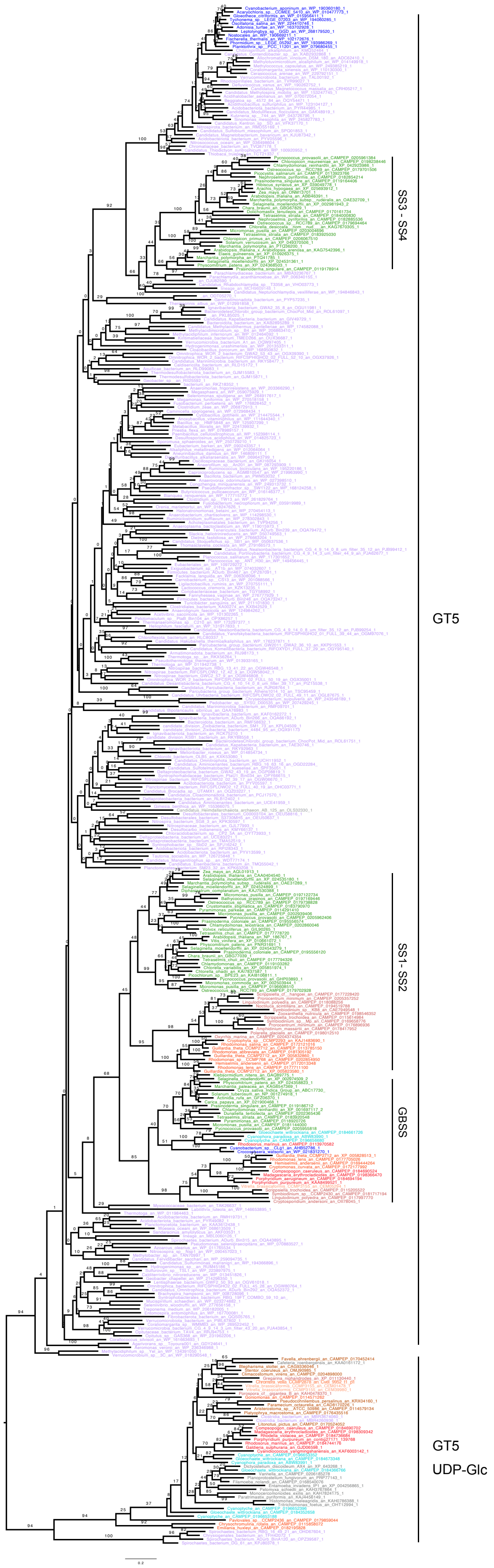
SS3 - SS4
GT5
SS1- SS2
GBSS
GT5
UDP-Glc

**Figure S16: Starch synthase / Glycogen synthase (GT5) phylogenetic tree.** The phylogenetic tree was built with IQTREE using LG4X matrix. The bootstrap values are indicated on the branch and are drawn from 100 bootstrap repetitions. Sequences are coloured depending on their origin: Chloroplastida are in green, Rhodophyceae are in Red, except for Cyanobacteria in dark blue, Bacteria are in purple, Glaucophyta in light blue, Cryptophyceae and haptophytes in orange, Myzozoa are in light brown except for Chromerids which are in light orange and Ciliophora which are in dark brown. Archaea are in dark grey. Finally other Eukaryotes are in grey. The phylogenetic analysis confirms that the dinoflagellates, compared to other alveolates, can be distinguished by the presence of starch synthases related to the SSI/SSII subgroup of starch synthases also found in green algae and plants (Figure 1B). An analogous form of enzyme was first documented in the cryptophyte *Guillardia theta* (*2–5*). This cryptophyte enzymes are sisters to those found in Dinoflagellates (BS=95). All other alveolate lineages synthesizing storage polysaccharides display glycogen/starch synthases belonging to the GT5 CAZy family commonly found in the eukaryotic cytosol where they participate in glycogen synthesis from UDP-Glc. This glycogen/starch synthase at the bottom of the tree is monophyletic and found in many diverse eukaryotes (BS of 94). Interestingly, Goniomonas, the heterotrophic plastid-less cryptophyte contains the standard eukaryotic GT5 GS (glycogen synthase) with no traces of the SSI/SSII-like enzyme of its photosynthetic plastid bearing relatives.

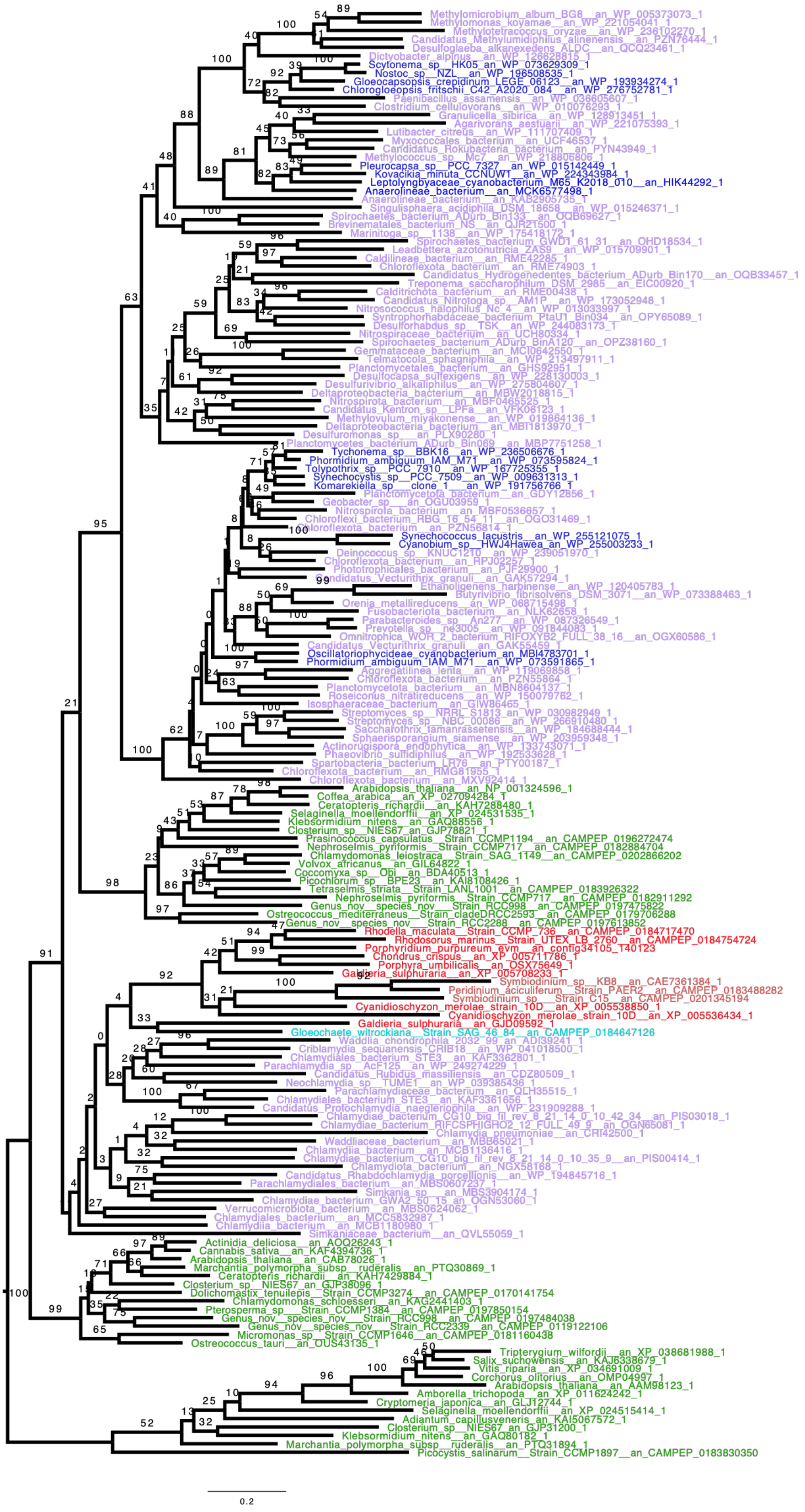

**Figure S17: Isoamylase (GH13_11) phylogenetic tree.** The phylogenetic tree was built with IQTREE using LG4X matrix. The bootstrap values are indicated on the branch and are drawn from 100 bootstrap repetitions. Sequences are colored depending on their origin: Chloroplastida are in green, Rhodophyceae are in Red, except for Cyanobacteria in dark blue, Bacteria are in purple, Glaucophyta in light blue, Cryptophyceae in orange, Myzozoa are in light brown except for Chromerids which are in light orange. Ciliophora are in dark brown. Finally other Eukaryotes are in grey. While isoamylase is distributed among all Archaepastida its presence in other eukaryotes is limited to the dinoflagellates which acquired it most probably from red algae. While glaucophytes are not excluded as candidate source for the dinoflagellate isoamylase in this phylogeny our previous analysis (*6*) with better sampled glaucophytes does exclude them.

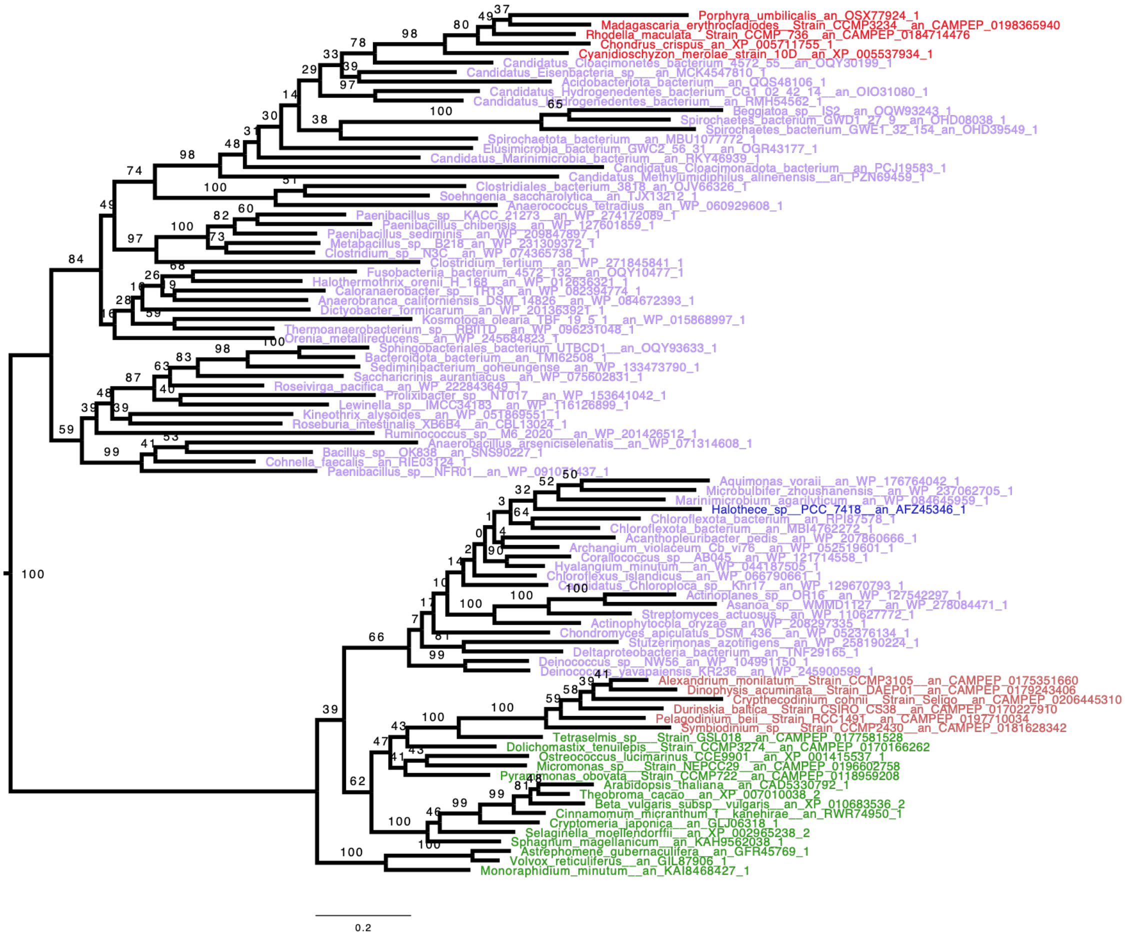


**Figure S18: Pullulanase phylogenetic tree.** The phylogenetic tree displayed here was built with IQTREE using LG4X matrix. The bootstrap values are indicated on the branch and are drawn from 100 bootstrap repetitions. Sequences are colored depending on their origin: Chloroplastida are in green, Rhodophyceae are in Red, Bacteria are in purple, Glaucophyta in light blue, Cryptophyceae in orange, Myzozoa are in light brown except for Chromerids which are in light orange. Ciliophora which are in dark brown. Finally other Eukaryotes are in grey. Within alveolates Pullulanase is only found in Dinoflagellates. The dinoflagellate enzyme is very likely derived from a green alga gene version (BS=100).

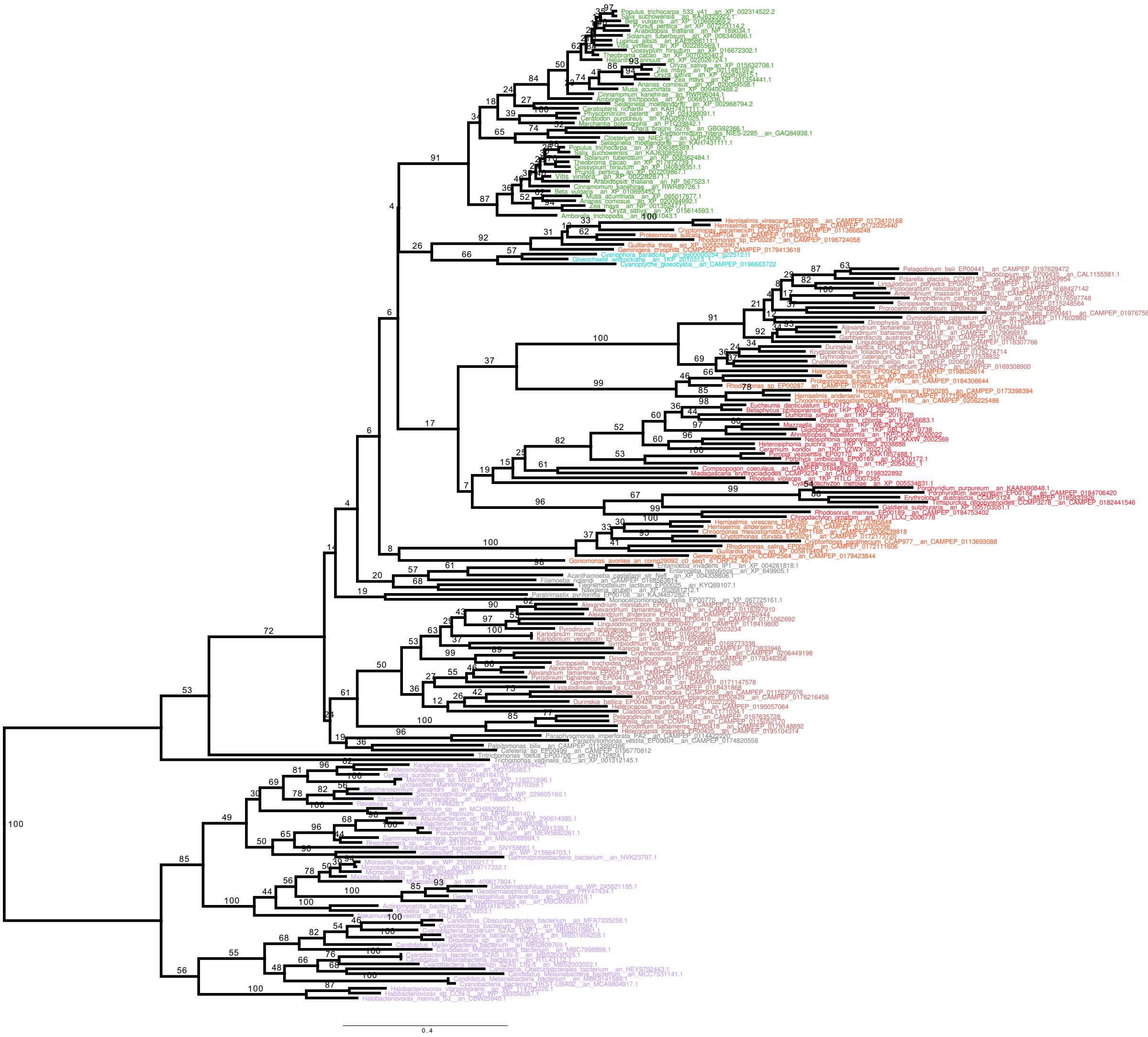

**Figure S19: β-amylase (BAM) (GH14) phylogenetic tree.** The phylogenetic tree displayed here was built with IQTREE using LG4X matrix. The bootstrap values are indicated on the branch and are drawn from 100 bootstraps. Sequences are colored depending on their origin: Chloroplastida are in green, Rhodophyceae are in Red, Bacteria in dark pink, Glaucophyta in light blue, Cryptista in purple, Myzozoa are in brown. Finally other Eukaryotes are in black. β-amylase genes within alveolates are seen exclusively in dinoflagellates. Two distinct isoforms display two separate origins consisting on the one hand of either red algae or cryptophyte and unknown protists on the other. In addition to cryptophytes and Archaeplastida, β-amylases are also found in glycogen accumulating protists. Furthermore, a few heterotrophic phagotrophs not known to synthesize α-glucans have this activity, such as the excavate Naegleria (*7*), which is accordingly suspected to be involved in prey digestion. It should be stressed that this β-maltose producing exo-enzyme is most often found together with the GH77 amylomaltase enzyme DPE2 which is able to metabolize its reaction product (maltose) (*8*). Note the complete absence of bacterial candidate sequences.

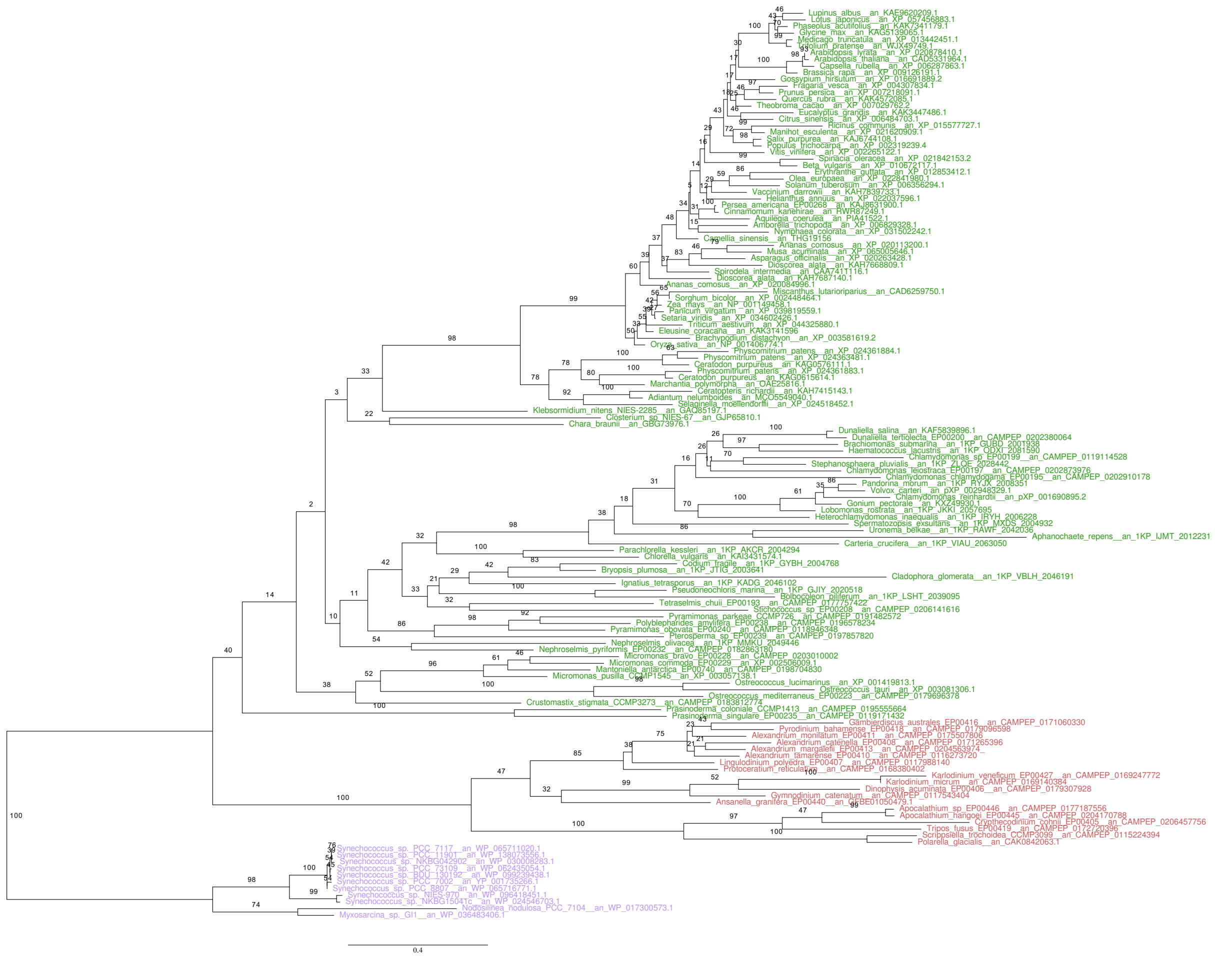

**Figure S20: Maltose Excess transporter (MEX) phylogenetic tree**. The phylogenetic tree displayed here was built with IQTREE using LG4X matrix. The bootstrap values are indicated on the branch and are drawn from 100 bootstraps. Sequences are colored depending on their origin: Chloroplastida are in green, Rhodophyceae are in Red, Bacteria in dark pink, Glaucophyta in light blue, Cryptista in purple, Myzozoa are in brown. Finally other Eukaryotes are in black. Dinoflagellates possess MEX1 (**M**altose **Ex**port), a maltose transporter identified in green algae and plant chloroplast responsible to export to the cytosol the maltose generated by plastidial α- and β-amylases. The single gene phylogeny of MEX1 does not allow to distinguish either between a green alga or a cyanobacterial source for the gene.

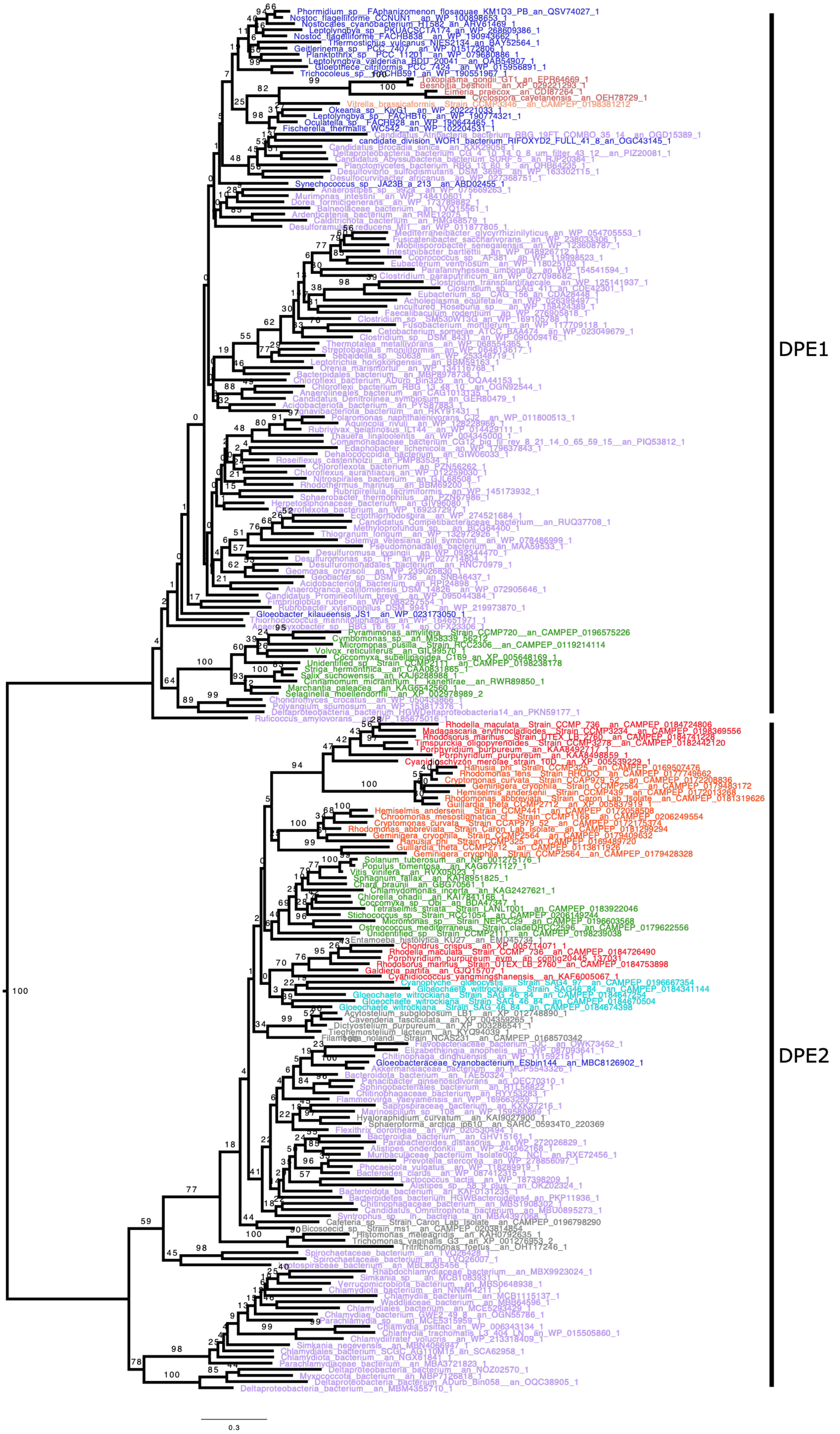
DPE1
DPE2
0.3

**Figure S21: Disproportionate enzyme 1-2 (DPE1-2) (GH77) phylogenetic tree**. The phylogenetic tree displayed here was built with IQTREE using LG4X matrix. The bootstrap values are indicated on the branch and are drawn from 100 boostrap repetitions. Sequences are coloured depending on their origin: Chloroplastida are in green, Rhodophyceae are in Red, except for Cyanobacteria in black, Bacteria are in purple, Glaucophyta in light blue, Cryptophyceae in orange, Myzozoa are in light brown except for Chromerids which are in orange and Ciliophora are in dark brown. Finally other Eukaryotes are in grey. DPE2 is found in several eukaryotes, at variance with DPE1 which is more widely distributed in Bacteria. Interestingly, Chromerid sequences branching with apicomplexa support a likely LGT from Cyanobacteria, however with low statistical support (BS=26). DPE2 is absent from all alveolates despite the presence of b-amylases in dinoflagellates

| Steps | Chloroplastida | Rhodophyta | Glaucophyta |
| --- | --- | --- | --- |
| 1 | + GT5-SS + BE + iDBE + PHO + b-AMY + DPE2 + a-AMY6 + a-AMY10 | + GT5-SS + BE + iDBE + PHO + b-AMY + DPE2 + a-AMY6 + a-AMY10 | + GT5-SS + BE + iDBE + PHO + b-AMY + DPE2 + a-AMY6 + a-AMY10 |
| 2 | + ISA + SSIII/IV + GBSS + GWD/PWD* | + ISA + SSIII/IV + GBSS + GWD/PWD* | + ISA + SSIII/IV + GBSS + GWD/PWD* |
| 3 | -iDBE -a-AMY10 | -iDBE -a-AMY10 | -a-AMY6 |
| 4 | | | Isoform Subfunctionalization |
| 5 | | Isoform number reduction, -SSIII/IV | |
| 6 | | + PUL | |
| 7 | MOS :+PHO +SSIII/IV +DPE1 | | |
| 8 | GLYCOGEN: +BE + PHO + SSIII/IV + SSI/II + ISA | | |
| 9 | STARCH: +ISA + BE + PHO + SSIII/IV + SSI/II + GBSS + GWD/PWD* + a-AMY6 + b-AMY + MEX* + PUL | | |
| 10 | Loss of all cytosolic starch enzymes save DPE2 and PHO | | |

**Table S2: Simplified view of Storage Polysaccharide Metabolism evolution following primary plastid endosymbiosis**. Starting from eukaryotic glycogen metabolism 10 steps involve multiple additions (symbolized by + preceding the enzyme name) of novel enzymes as well as a few cases of enzyme losses (symbolized by -). The identity of the gene source is in each case represented by the colour used for the enzyme. Orange/brown is used for host genes, red for chlamydial genes, blue for genes of cyanobacterial origin and black for genes of other bacterial origin. Novel genes evolved by the host following primary plastid endosymbiosis are indicated by an asterisk. Step 1 represents the status of eukaryotic glycogen metabolism in the host cytosol before plastid endosymbiosis as it is still found today in diverse protists such as *Entamoeba histolytica* or *Trichomonas vaginalis* (*9*). This status can be derived by examining the nature and phylogeny of these enzymes in Archaeplastida. a-AMY6 and a-AMY10 represent two distinct types of GH13 a amylase enzymes of respectively subfamilies 6 and 10. Step 2 represents the early transition from glycogen to starch that occurred after plastid endosymbiosis. This required LGT of the chlamydial effector gene encoding isoamylase. Progressive transition to starch through ISA required de novo coevolution of GWD/PWD. Acquisition of the chlamydial effector SSIII/IV was required both to establish photosymbiosis through ADP-Glc assimilation and also possibly to ensure starch granule synthesis priming (*7*, *8*, *10*). Acquisition of GBSS from a cyanobacterial source terminates the evolution of glycogen into starch metabolism in the cytosol. Step 3 display the losses of enzymes of the cytosolic network experienced from the 3 archaeplastida lineages. Step 4 represent the duplication of genes followed by the specialization of distinct isoforms into distinct functions which characterizes the glaucophyte lineage. In contrast to the gene number expansion seen in glaucophytes, step 5 and 6 represent a very strong opposite trend seen in red algae in line with the genomic simplification experienced by these algae where all catalytic activies were reduced to a single enzyme isoform with the noticeable exception of a-1,4 glucanotransferases of the DPE2 type and isoamylase. Loss of the chlamydial effector gene SSIII/IV and acquisition of pullulanase from bacteria terminates the evolution of the rhodophyte starch metabolism network. Evolution of novel LHCs in the rising green lineage (Chloroplastida) was accompanied by the requirement for progressively more abundant storage polysaccharide pools within the chloroplast (*11*). This requirement was met by increasingly larger pools of MOS (malto-oligosaccharides), glycogen and starch (steps 7 to 9). This led to a complex pattern of gene duplications from the preexisting cytosolic starch metabolism network that were required for building successively MOS (step 7) then glycogen (step 8) and finally starch (step 9) within the chloroplast (for review see (*8*, *11*). The transition from glycogen to starch in the chloroplast signalled the disappearance of starch metabolism from the cytosol. Plastidial starch metabolism correlates with the evolution of a new maltose transporter named MEX (maltose export) exporting maltose to the host cytosol which has kept a cytosolic form of phosphorylase and DPE2 for assimilation of this disaccharide.

#### References


1. J. Yang, Z. He, C. Chen, J. Zhao, R. Fang, Starch Branching Enzyme 1 Is Important for Amylopectin Synthesis and Cyst Reactivation in Toxoplasma gondii. *Microbiol Spectr* **10**, e0189121 (2022).

2. M. M. Nielsen, C. Ruzanski, K. Krucewicz, A. Striebeck, U. Cenci, S. G. Ball, M. M. Palcic, J. A. Cuesta-Seijo, Crystal Structures of the Catalytic Domain of Arabidopsis thaliana Starch Synthase IV, of Granule Bound Starch Synthase From CLg1 and of Granule Bound Starch Synthase I of Cyanophora paradoxa Illustrate Substrate Recognition in Starch Synthases. *Front. Plant Sci.* **9**, 1138 (2018).

3. P. Deschamps, I. Haferkamp, D. Dauvillée, S. Haebel, M. Steup, A. Buléon, J.-L. Putaux, C. Colleoni, C. d'Hulst, C. Plancke, S. Gould, U. Maier, H. E. Neuhaus, S. Ball, Nature of the Periplastidial Pathway of Starch Synthesis in the Cryptophyte *Guillardia theta*. *Eukaryot Cell* **5**, 954–963 (2006).

4. I. Haferkamp, P. Deschamps, M. Ast, W. Jeblick, U. Maier, S. Ball, H. E. Neuhaus, Molecular and Biochemical Analysis of Periplastidial Starch Metabolism in the Cryptophyte *Guillardia theta*. *Eukaryot Cell* **5**, 964–971 (2006).

5. B. A. Curtis, G. Tanifuji, F. Burki, A. Gruber, M. Irimia, S. Maruyama, M. C. Arias, S. G. Ball, G. H. Gile, Y. Hirakawa, J. F. Hopkins, A. Kuo, S. A. Rensing, J. Schmutz, A. Symeonidi, M. Elias, R. J. M. Eveleigh, E. K. Herman, M. J. Klute, T. Nakayama, M. Oborník, A. Reyes-Prieto, E. V. Armbrust, S. J. Aves, R. G. Beiko, P. Coutinho, J. B. Dacks, D. G. Durnford, N. M. Fast, B. R. Green, C. J. Grisdale, F. Hempel, B. Henrissat, M. P. Höppner, K.-I. Ishida, E. Kim, L. Kořený, P. G. Kroth, Y. Liu, S.-B. Malik, U. G. Maier, D. McRose, T. Mock, J. A. D. Neilson, N. T. Onodera, A. M. Poole, E. J. Pritham, T. A. Richards, G. Rocap, S. W. Roy, C. Sarai, S. Schaack, S. Shirato, C. H. Slamovits, D. F. Spencer, S. Suzuki, A. Z. Worden, S. Zauner, K. Barry, C. Bell, A. K. Bharti, J. A. Crow, J. Grimwood, R. Kramer, E. Lindquist, S. Lucas, A. Salamov, G. I. McFadden, C. E. Lane, P. J. Keeling, M. W. Gray, I. V. Grigoriev, J. M. Archibald, Algal genomes reveal evolutionary mosaicism and the fate of nucleomorphs. *Nature* **492**, 59–65 (2012).

6. T. Maeno, Y. Yamakawa, Y. Takiyasu, H. Miyauchi, Y. Nakamura, M. Ono, N. Ozaki, Y. Utsumi, U. Cenci, C. Colleoni, S. Ball, M. Tsuzuki, S. Fujiwara, One of the isoamylase isoforms, CMI294C, is required for semi-amylopectin synthesis in the rhodophyte Cyanidioschyzon merolae. *Front. Plant Sci.* **13**, 967165 (2022).

7. S. G. Ball, C. Colleoni, D. Kadouche, M. Ducatez, M.-C. Arias, C. Tirtiaux, Toward an understanding of the function of Chlamydiales in plastid endosymbiosis. *Biochimica et Biophysica Acta (BBA) - Bioenergetics* **1847**, 495–504 (2015).

8. S. Ball, C. Colleoni, U. Cenci, J. N. Raj, C. Tirtiaux, The evolution of glycogen and starch metabolism in eukaryotes gives molecular clues to understand the establishment of plastid endosymbiosis. *Journal of Experimental Botany* **62**, 1775–1801 (2011).

9. S. Ball, C. Colleoni, M. C. Arias, "The Transition from Glycogen to Starch Metabolism in Cyanobacteria and Eukaryotes" in *Starch*, Y. Nakamura, Ed. (Springer Japan, Tokyo, 2015; http://link.springer.com/10.1007/978-4-431-55495-0_4), pp. 93–158.

10. S. G. Ball, A. Subtil, D. Bhattacharya, A. Moustafa, A. P. M. Weber, L. Gehre, C. Colleoni, M.-C. Arias, U. Cenci, D. Dauvillée, Metabolic Effectors Secreted by Bacterial Pathogens: Essential Facilitators of Plastid Endosymbiosis? *The Plant Cell* **25**, 7–21 (2013).

11. P. Deschamps, I. Haferkamp, C. d'Hulst, H. E. Neuhaus, S. G. Ball, The relocation of starch metabolism to chloroplasts: when, why and how. *Trends Plant Sci* **13**, 574–582 (2008).
